# A taxonomy of transcriptomic cell types across the isocortex and hippocampal formation

**DOI:** 10.1101/2020.03.30.015214

**Authors:** Zizhen Yao, Thuc Nghi Nguyen, Cindy T. J. van Velthoven, Jeff Goldy, Adriana E. Sedeno-Cortes, Fahimeh Baftizadeh, Darren Bertagnolli, Tamara Casper, Kirsten Crichton, Song-Lin Ding, Olivia Fong, Emma Garren, Alexandra Glandon, James Gray, Lucas T. Graybuck, Daniel Hirschstein, Matthew Kroll, Kanan Lathia, Boaz Levi, Delissa McMillen, Stephanie Mok, Thanh Pham, Qingzhong Ren, Christine Rimorin, Nadiya Shapovalova, Josef Sulc, Susan M. Sunkin, Michael Tieu, Amy Torkelson, Herman Tung, Katelyn Ward, Nick Dee, Kimberly A. Smith, Bosiljka Tasic, Hongkui Zeng

## Abstract

The isocortex and hippocampal formation are two major structures in the mammalian brain that play critical roles in perception, cognition, emotion and learning. Both structures contain multiple regions, for many of which the cellular composition is still poorly understood. In this study, we used two complementary single-cell RNA-sequencing approaches, SMART-Seq and 10x, to profile ∼1.2 million cells covering all regions in the adult mouse isocortex and hippocampal formation, and derived a cell type taxonomy comprising 379 transcriptomic types. The completeness of coverage enabled us to define gene expression variations across the entire spatial landscape without significant gaps. We found that cell types are organized in a hierarchical manner and exhibit varying degrees of discrete or continuous relatedness with each other. Such molecular relationships correlate strongly with the spatial distribution patterns of the cell types, which can be region-specific, or shared across multiple regions, or part of one or more gradients along with other cell types. Glutamatergic neuron types have much greater diversity than GABAergic neuron types, both molecularly and spatially, and they define regional identities as well as inter-region relationships. For example, we found that glutamatergic cell types between the isocortex and hippocampal formation are highly distinct from each other yet possess shared molecular signatures and corresponding layer specificities, indicating their homologous relationships. Overall, our study establishes a molecular architecture of the mammalian isocortex and hippocampal formation for the first time, and begins to shed light on its underlying relationship with the development, evolution, connectivity and function of these two brain structures.

## INTRODUCTION

The cerebral cortex occupies a large portion of the mammalian brain (Azevedo et al., 2009) and is uniquely situated to execute multiple functions, from sensory perception and generation of voluntary behavior to emotion, cognition, learning and memory. The cortex is partitioned into multiple areas, each having specific input and output connections with many subcortical and other cortical regions (Rakic, 2009; Van Essen and Glasser, 2018). This area specialization is likely a major contributing factor to its functional diversity (Clowry et al., 2018).

Developmentally, the cortex originates from pallium, a main part of the telencephalon, which can be divided into several parts (Pessoa et al., 2019). Medial pallium gives rise to the hippocampal formation (also called archicortex), ventral pallium gives rise to the olfactory cortex (also called paleocortex), and, in mammals, dorsal pallium gives rise to the isocortex (also called neocortex). Archicortex and paleocortex (collectively also called allocortex) are evolutionarily older structures, whereas iso/neocortex emerged later and substantially expanded in vertebrate evolution, culminating in its current form in the mammalian brain (Rakic, 2009). It is generally thought that neurons in the archicortex and paleocortex are arranged in 3 to 5 layers, whereas those in iso/neocortex populate 6 layers. However, laminar structure varies substantially across different cortical areas, and the traditional cytoarchitecture-based delineation of layers likely underrepresents the cellular diversity both within and between different cortical regions.

Numerous molecular, anatomical, and physiological studies have revealed a broad variety of neuronal cell types in different cortical and hippocampal regions. These cell types display a correspondingly large variety of cellular properties that are likely related to specific functions in the cortical circuits they are embedded in (Zeng and Sanes, 2017). But absent of a systematic study, we still do not have a complete picture of the number of cell types in the cortex and how they are distributed across different regions. This knowledge is an essential prerequisite toward understanding how different cortical regions interact with the rest of the brain and how they carry out their individual functions.

Functional areas of the neocortex (∼180 in humans and ∼30 in mice) (Van Essen and Glasser, 2018) tile the cortical sheet and include primary and higher-order sensory areas across all sensory modalities, primary and secondary motor areas, and multiple associational areas in the frontal, medial, and lateral parts of the neocortex that perform a variety of integrative functions. Extensive connectivity tracing studies and *in vivo* neural imaging studies have shown that these neocortical areas together form a hierarchical neural network with functionally distinct modules and feedforward and feedback pathways both within and between modules (Coogan and Burkhalter, 1993; Felleman and Van Essen, 1991; Harris et al., 2019; Markov et al., 2014). The two major classes of neurons in the isocortex, glutamatergic and GABAergic, can be further divided into multiple subclasses and types (Tasic et al., 2018). Glutamatergic excitatory neuronal types are organized by layers and their long-range projection patterns (Harris and Shepherd, 2015), whereas GABAergic inhibitory interneuron types are organized by their firing characteristics and local connectivity patterns (Tremblay et al., 2016). The tremendous diversity in interareal axon-projection patterns of specific glutamatergic neuron types from specific isocortical areas is thought to form the structural basis of the hierarchical network (D’Souza et al., 2016; Harris et al., 2019).

The hippocampal formation is also a complex multi-areal structure, which includes the hippocampal region, the subicular complex, and the medial and lateral entorhinal cortex. Like in the isocortex, glutamatergic excitatory and GABAergic inhibitory neurons are the two major classes of neurons, and each contains multiple types. Hippocampal and isocortical GABAergic interneuron types exhibit similar molecular, anatomical, and physiological properties (Pelkey et al., 2017; Tremblay et al., 2016). The glutamatergic neuron types in all subregions of the hippocampal formation have not been fully categorized, but many exhibit specific connections within the hippocampal formation and with other brain regions (Bienkowski et al., 2018; van Strien et al., 2009). Together, these diverse neuron types form a network that underlies the many functions of the hippocampal formation – learning, memory, spatial navigation, and regulation of emotions. In particular, there is a prominent functional transition along the dorsal-ventral axis of the hippocampal formation likely built upon the differential networks of cell types and connections, with the dorsal network mainly mediating spatial navigation and the ventral network mainly mediating emotional behaviors (Cembrowski and Spruston, 2019).

We and others have used the single-cell RNA-sequencing (scRNA-seq) approach to systematically characterize and classify cell types in individual regions of the isocortex and hippocampus (Harris et al., 2018; Hodge et al., 2019; Tasic et al., 2016, 2018; Zeisel et al., 2015) (Yao et al, https://www.biorxiv.org/content/10.1101/2020.02.29.970558v2). In our previous study, we analyzed ∼24,000 single-cell transcriptomes to define 133 cell types (61 GABAergic, 56 glutamatergic and 16 non-neuronal) in two distantly-located isocortical areas, the primary visual cortex (VISp) and anterolateral motor cortex (ALM) in mice (Tasic et al., 2018). We found that the two areas have shared GABAergic interneuron types but distinct glutamatergic neuron types. Many of these transcriptomic types are correlated with previously identified isocortical cell types by well-characterized marker genes and transgenic driver lines. More recently, using the Patch-seq approach, we simultaneously profiled the transcriptomic, electrophysiological, and morphological properties of a large set of GABAergic interneurons from the mouse visual cortex and found excellent correspondence among the three modalities (Gouwens et al., https://www.biorxiv.org/content/10.1101/2020.02.03.932244v1). Similar findings were made in another Patch-seq study focused on the mouse primary motor cortex (Scala et al., https://www.biorxiv.org/content/10.1101/2020.02.03.929158v1). These studies demonstrate the validity and power of using the highly scalable scRNA-seq approach to generate a comprehensive census of cell types as a foundation for further structural and functional studies of brain circuits.

Here we have extended our scRNA-seq analysis to cover all regions of the adult mouse isocortex (CTX) and hippocampal formation (HPF), analyzing ∼1.2 million cells with two different scRNA-seq platforms (10x and SMART-seq). We developed a consensus clustering approach to combine the two datasets and derived a cell type taxonomy comprising 379 transcriptomic types, many of which are new. We found that most GABAergic interneuron types are shared across all regions, but also identified several interneuron types specific to the hippocampus. On the other hand, isocortex and hippocampal formation have largely distinct but homologous sets of glutamatergic excitatory neuron types that collectively define a highly complex landscape. Many isocortical glutamatergic types are shared across multiple areas and exhibit gene expression variations along the cortical sheet. Similarly, hippocampal and subicular glutamatergic types are organized along multiple spatial dimensions. The comprehensive catalog of transcriptomic cell types presented here creates many opportunities for cell type-specific investigation of the function of these brain circuits.

## RESULTS

### A taxonomy of transcriptomic cell types across isocortex and hippocampal formation

To conduct large-scale single-cell transcriptomic characterization of cell types in a standardized, comprehensive and efficient manner, we used two complementary pipelines: SMART-Seq v4 (SSv4) (Tasic et al., 2018), and 10x Genomics Chromium platform based on version 2 chemistry (10xv2). All newly acquired transcriptomes were obtained from cells isolated by fluorescence-activated cell sorting (FACS) and combined with the previous VISp/ALM SSv4 dataset (Tasic et al., 2018) to arrive at the final datasets (**Fig. S1A, S2**).

We used Allen Mouse Brain Common Coordinate Framework version 3 (CCFv3) ontology (http://atlas.brain-map.org/) to define brain regions for profiling and boundaries for dissections. We chose sampling at mid-ontology level and covered all regions of isocortex (CTX) and hippocampal formation (HPF), listed here for reference. The covered areas in CTX are frontal pole (FRP), primary motor (MOp), secondary motor (MOs), primary somatosensory (SSp), supplemental somatosensory (SSs), gustatory (GU), visceral (VISC), auditory (AUD), primary visual (VISp), anterolateral visual (VISal), anteromedial visual (VISam), lateral visual (VISl), posterolateral visual (VISpl), posteromedial (VISpm), laterointermediate (VISli), postrhinal (VISpor), anterior cingulate (ACA), prelimbic (PL), infralimbic (ILA), orbital (ORB), agranular insular (AI), retrosplenial (RSP), posterior parietal association (PTLp), temporal association (TEa), perirhinal (PERI), and ectorhinal (ECT) areas. The covered regions in HPF are divided into two main parts, the hippocampal region (HIP) itself which includes fields CA1, CA2, CA3, and dentate gyrus (DG), and the retrohippocampal region (RHP) which includes lateral entorhinal area (ENTl), medial entorhinal area (ENTm), parasubiculum (PAR), postsubiculum (POST), presubiculum (PRE), subiculum (SUB), and prosubiculum (ProS). The few remaining small regions in HPF, i.e., fasciola cinereal (FC), induseum griseum (IG), hippocampo-amygdalar transition area (HATA), and area prostriata (APr), were included in the dissection of their neighboring regions.

For tissue dissections, we chose to combine some neighboring regions (**Fig. 1A**, **S1B, Table S1**), to avoid microdissections of very small regions and to satisfy the requirement for sufficient numbers of cells for profiling, especially for 10xv2. For example, due to the close proximity with each other, PAR, POST and PRE were combined into a single dissection region named PPP for SSv4 profiling, and PPP was further combined with SUB and ProS into a joint dissection region named PPP-SP for 10xv2 profiling. Note that the manual microdissections are imperfect, but despite the presence of cells from neighboring regions, the dissections still contain substantial enrichment of cells for the targeted regions. In total, for 10xv2, we profiled 12 joint regions for CTX and 3 joint regions for HPF with 30,000-300,000 cells per region (**Fig. S1B, D, Table S1**). For SSv4, we profiled 17 regions for CTX and 5 regions for HPF with 1,100-14,000 cells per region (**Fig. S1B, C, Table S1**). For comparative reasons (see below), we also included ∼1,000 SSv4 cells from the claustrum (CLA), a region that does not belong to either CTX or HPF.

**Figure 1.**
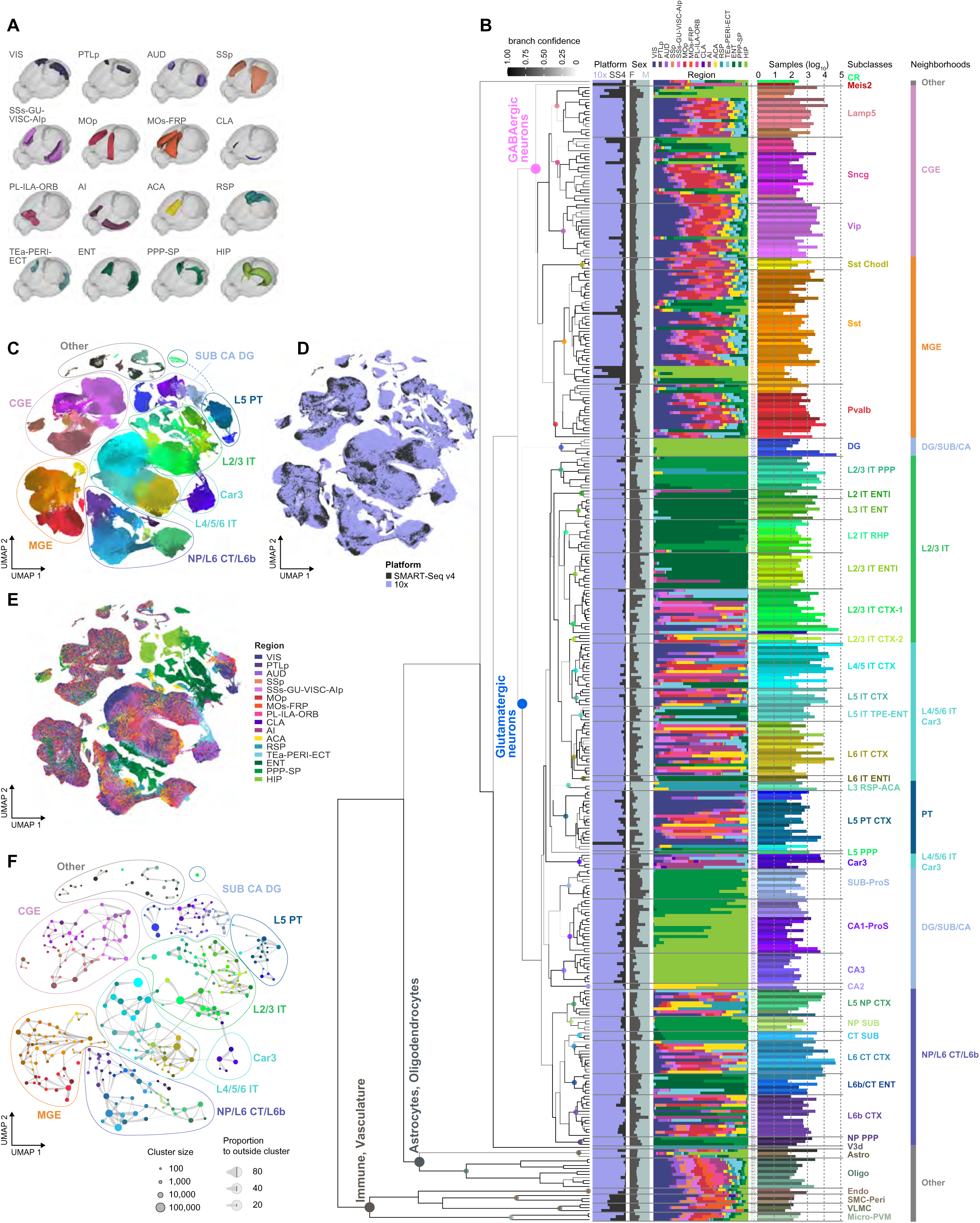
Overall cell type taxonomy of the isocortex (CTX) and hippocampal formation (HPF). (A) Overview of sampled brain regions rendered in Allen Mouse Brain Common Coordinate Framework v3 (CCFv3). Included CTX areas are: visual (VIS), posterior parietal association (PTLp), auditory (AUD), primary somatosensory (SSp), supplemental somatosensory (SSs), gustatory (GU), visceral (VISC), agranular insular posterior part (AIp), primary motor (MOp), secondary motor (MOs), frontal pole (FRP), prelimbic (PL), infralimbic (ILA), orbital (ORB), agranular insular (AI), anterior cingulate (ACA), retrosplenial (RSP), association (TEa), perirhinal (PERI), and ectorhinal (ECT). Included HPF regions are: entorhinal cortex (ENT), presubiculum, parasubiculum, postsubiculum, subiculum, and prosubiculum (PPP-SP), and hippocampus (HIP). A non-cortical region, claustrum (CLA), is also included. **(B)** The transcriptomic taxonomy tree of 379 cell types (clusters) organized in a dendrogram derived from a joint analysis of single cell transcriptomes obtained by 10x Chromium version 2 (10xv2; *n* = 1,089,022) and SMART-Seq v4 (SSv4; *n* = 74,705). The dendrogram was constructed by hierarchical clustering of median cluster gene expression for 4,963 differentially expressed genes. Cell classes are labeled at each branch point on the left. Cell subclasses are represented with colored dots on the tree and matching labels to the right. Subclasses are grouped into neighborhoods (further right). Bar plots represent fractions of cells profiled according to platform, sex, and region, and the total number of cells per cluster on a log_10_ scale. **(C-E)** UMAP representation of the same dataset as in **B**, colored by cluster (**C**), platform (**D**) or region (**E**). In **C**, Clusters are grouped into eight neighborhoods: (1) CGE-derived neurons, (2) MGE-derived neurons, (3) L2/3 IT neurons, (4) L4/5, L5, L6 IT and Car3 neurons, (5) L5 PT neurons, (6) L5 NP, L6 CT, and L6b neurons, (7) subiculum, prosubiculum, CA1, CA2, CA3, and dentate gyrus neurons, (8) Other, which includes Cajal-Retzius (CR) cells, Meis2 cells, 3^rd^ ventricle cells (V3d), and non-neuronal cells. The UMAP was calculated for cells in **B** based on 59 principal components (PCs) and single cell expression of 4,963 differentially expressed (DE) genes (**Methods**). **(F)** Constellation plot of the same dataset as in **B-E**. Each cluster is represented by a dot, positioned at the cluster centroid in UMAP coordinates in **C**, with dot size proportional to the number of cells in the cluster. Connecting lines represent the proportion of nearest neighbors between pairs of clusters (**Methods**). Clusters are grouped by neighborhood.

We used transgenic driver lines for fluorescence-positive cell isolation to enrich for neurons. We used mice of both sexes (**Fig. S1E-F**), in congenic C57BL/6J background, euthanized at postnatal day (P)53-56 (>90% of 586 mice total), with the vast majority being transgenic Cre driver lines crossed to the *Ai14*-tdTomato reporter (Madisen et al., 2010) (**Table S2**). All 10xv2 cells were isolated from the pan-neuronal *Snap25-IRES2-Cre* line crossed to *Ai14* (**Fig. S1H**). For SSv4, transgenic mice, dissection schemes and sampling rates varied by regions, and cells labeled by Retro-seq were also included (**Methods**; **Fig. S1G, Table S2**).

Overall, we obtained 1,138,179 single-cell 10xv2 transcriptomes and 76,307 single-cell SSv4 transcriptomes after the quality control (QC) process (**Fig. S2**). We first clustered 10xv2 and SSv4 cells separately, resulting in 372 10xv2 clusters and 291 SSv4 clusters (excluding low-quality clusters) (**Fig. S3A, Methods**). To perform consensus clustering, we developed an anchor-based integrative clustering algorithm across multiple data modalities, available via the **i_harmonize** function of the **scrattch.hicat** package (https://github.com/AllenInstitute/scrattch.hicat; **Methods**; **Fig. S3B**). Integrative clustering of the 10xv2 and SSv4 datasets resulted in 379 consensus clusters after excluding the low-quality ones (**Fig. 1, S3A, Table S3**).

Post-clustering, we constructed a taxonomy tree (**Fig. 1B**) by hierarchical clustering of transcriptomic clusters based on the average gene expression per cluster of 4,963 differentially expressed (DE) genes (top 50 DE genes in each direction between every pair of clusters). We explored the relationships among the 379 clusters by visualizing cells belonging to them with UMAP projections (**Fig. 1C-E**) and constellation plots (**Fig. 1F**). For scalability reasons, our current implementation of constellation plots does not rely on core and intermediate cells as in previous work (Tasic et al., 2016, 2018) but a new nearest-neighbor approach. Briefly, for each cell, we identify 15 nearest neighbors in the reduced dimension space, and then at the cluster level (represented by a dotted node) we use edges to show the fraction of nearest neighbors that are cells outside the cluster (**Methods; Fig. 1F**).

These different approaches to explore the cell type landscape, which is defined by a combination of discrete and continuous gene expression variation, provide a holistic description of the taxonomy. For example, taxonomical trees are simple but artificially discrete: they do not preserve all the multidimensional relationships among types, but they provide a hierarchical view of the relationships which are less clear in UMAP representations. UMAP representations and constellation plots enable visualization of continuity in addition to discreteness. Based on all these, we label sets of cells within the taxonomy from coarse to fine categories as: class, neighborhood, subclass, supertype, type (**Fig. 1B, C, F**). Some of these categories are discrete, but, especially at the level of types (i.e., individual clusters that are leaf nodes or leaves in taxonomy), they are not.

To be able to interpret and understand this taxonomy which contains many new transcriptomic types, we collated sets of DE genes selective to each cluster as well as each branch of the taxonomy to represent different levels of granularity, and examined the anatomical expression patterns of these genes using the Allen Brain Atlas (ABA) *in situ* hybridization (ISH) data (Lein et al., 2007). Based on such extensive anatomical annotation as well as prior knowledge, we assigned the 379 clusters into 4 classes, 8 neighborhoods, 43 subclasses, and 90 supertypes. For each of the four major cell classes: the GABAergic neuronal class contains 6 subclasses and 118 clusters; the glutamatergic neuronal class contains 28 subclasses and 237 clusters; the astrocyte/oligodendrocyte non-neuronal class contains 2 subclasses and 13 clusters; and the immune/vascular non-neuronal class contains 4 subclasses and 11 clusters (**Fig. 1B, Table S3**). Partly based on the distance relationships revealed in our analysis and partly for practical purposes, we grouped the subclasses into 8 neighborhoods, 2 GABAergic (CGE and MGE), 5 glutamatergic (L2/3 IT, L4/5/6 IT/Car3, L5 PT, NP/CT/L6b, and SUB/CA/DG), and one ‘Other’ neighborhood (**Fig. 1B, C, F**).

Detailed analyses of the neuronal neighborhoods are presented in the following sections. We briefly mention here the ‘Other’ neighborhood. This neighborhood included all non-neuronal subclasses, as well as three neuronal types, Cajal-Retzius (CR) (cluster #1, glutamatergic, mostly in layer 1), Meis2 (cluster #2, GABAergic, mostly in white matter), and V3d (cluster #355, glutamatergic, containing cells isolated from RSP-specific dissections that were likely from the third ventricle), because in the taxonomy tree they formed three distant branches preceding the major glutamatergic/GABAergic split (**Fig. 1B**), as shown before (Tasic et al., 2018). For the astro/oligo class, we identified 3 astrocyte clusters (#356-358) and 10 oligodendrocyte clusters (#359-368), 2 of which were oligodendrocyte progenitor cell (OPC) clusters (#359-360). For the immune/vascular class, there were 2 endothelial cell clusters (#369-370), 3 smooth muscle cell (SMC) and pericyte clusters (#371-373), 3 vascular/leptomeningeal cell (VLMC) clusters (#374-376) and 3 microglia/perivascular macrophage (PVM) clusters (#377-379). Overall, since the cell isolation in this study was aimed towards enrichment for neurons, we only had limited sampling of non-neuronal cells (12,513 10xv2 cells and 1,958 SSv4 cells after QC). Thus, the description of non-neuronal cell diversity presented here is likely incomplete, and we intend to address this issue in future studies.

Despite the large difference in gene detection per cell for the two platforms (7,593 genes per cell for SSv4 and 4,132 for 10xv2; **Fig. S4A-B**), there was good correlation for the numbers of genes detected at each cluster level between the two methods (**Fig. S4D-E**). We found that the 10xv2 and SSv4 cells usually commingle in nearly all clusters, except 13 clusters with only 10xv2 cells and 2 with only SSv4 cells (**Fig. 1B, D, Table S3**). We also found a high degree of concordance between the consensus clusters and clusters derived from SSv4-only or 10xv2-only data (**Fig. S3C, S5**). In addition, we found generally good, albeit variable, correspondence between our current transcriptomic taxonomy and previously published ones from either us (Tasic et al., 2018) or others (Cembrowski et al., 2018; Harris et al., 2018; Saunders et al., 2018; Zeisel et al., 2018) (**Fig. S3C, S6**). We provide a web interface to present summarized information about each cell type in the form of a cell type card (https://taxonomy.shinyapps.io/ctx_hip_browser/; **Fig. S7**). Each cell type card includes information about cluster annotation, sampling metrics, marker gene expression, relationship to other clusters within its neighborhood and the equivalent cluster in previously published datasets (if present).

### GABAergic cell type taxonomy

The GABAergic inhibitory neuronal class is divided into two neighborhoods that correlate with distinct developmental origins: caudal ganglionic eminence (CGE) (**Fig. 2A-F**) and medial ganglionic eminence (MGE) (**Fig. 2G-L**). Note that some of the CGE cell types (e.g., the neurogliaform cells) may, in fact, be developmentally derived from the nearby preoptic region (PO) (Niquille et al., 2018). Each neighborhood is further divided into 3 subclasses: Lamp5, Sncg and Vip in CGE (**Fig. 2A**), and Sst Chodl, Sst and Pvalb in MGE (**Fig. 2G**).

**Figure 2.**
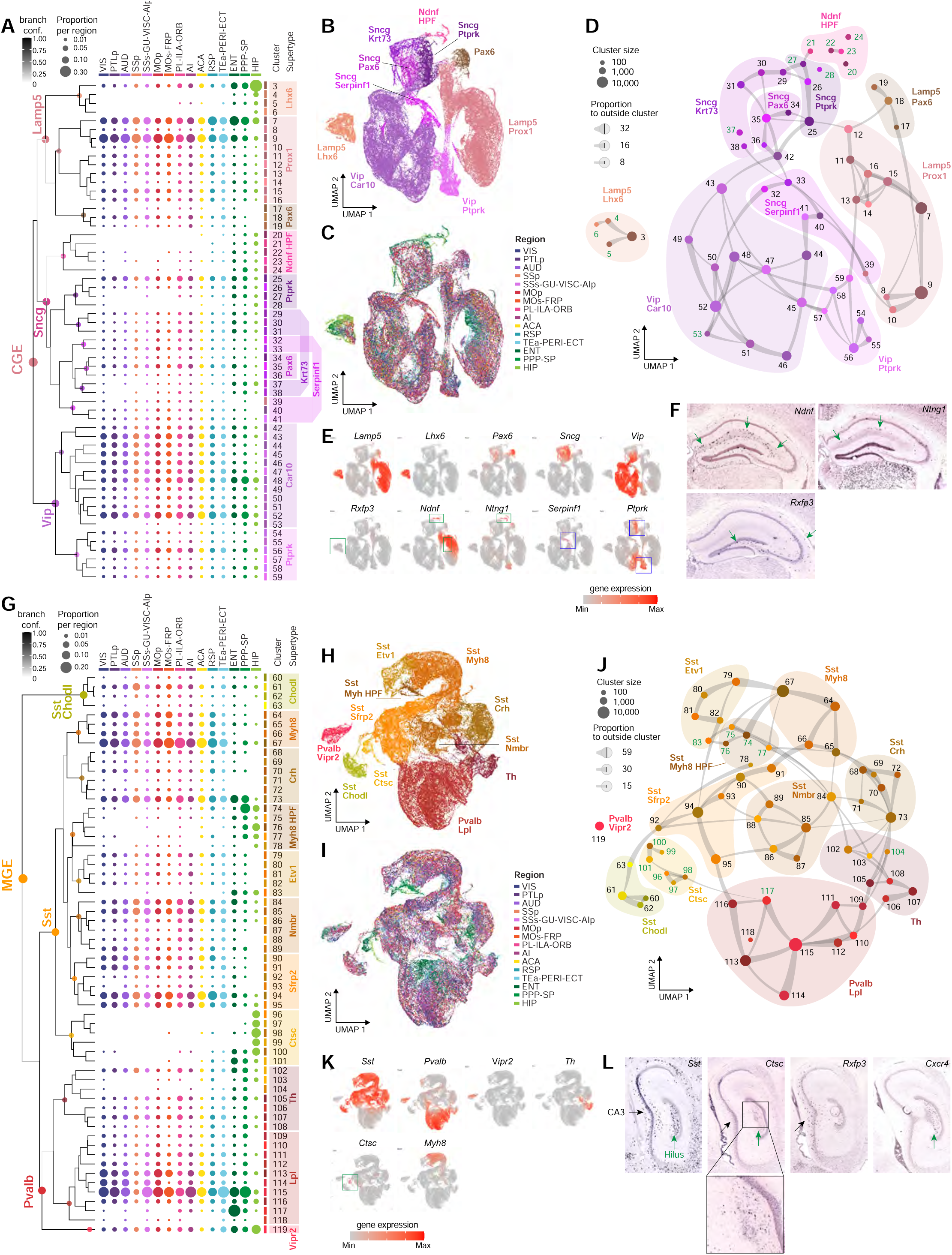
GABAergic cell types of CTX and HPF. **(A)** Dendrogram of 57 CGE types followed by the proportion of cells (dot size) within each cluster derived from each region noted at the top of the graph (the values in each region/column add up to 100%). The dendrogram is a part of the dendrogram in Figure 1B focusing on the CGE-derived neuronal types. Clusters are grouped into 10 supertypes on the right, which represent the level of organization below subclass. Cells included: 10xv2: *n* = 91,663; SSv4: *n* = 12,682. **(B-C)** UMAP representation of CGE types, colored by supertype and annotated with major supertype marker genes (**B**) or by region (**C**). This UMAP was calculated for cells in **A** based on 17 PCs derived from 4,963 DE genes. **(D)** Constellation plot for CGE types indicating relatedness of clusters. Each cluster is represented by a dot, positioned at the cluster centroid in UMAP coordinates in **B**. Clusters are grouped into supertypes. Cluster labels colored green are clusters where more than 80% of cells are derived from HPF. **(E)** UMAP representation of CGE types, as in **B**, showing expression of select supertype marker genes in red. The green boxes highlight HPF specific clusters within UMAP. The blue boxes highlight transitions between supertypes. **(F)** RNA ISH from the Allen Mouse Brain Atlas (ABA) for select markers expressed in the HPF-enriched CGE supertypes. The *Ntng1* gene is specifically expressed by the Ndnf HPF supertype, whereas the *Rxfp3* gene is specific to HPF-enriched clusters in the Lamp5 Lhx6 supertype. **(G)** Dendrogram of 60 MGE types followed by the proportion of cells (dot size) within each cluster derived from each region where the values in each region (column) add up to 100%. The clusters are grouped into 11 supertypes. Cells included: 10xv2: *n* = 75,304; SSv4: *n* = 9,891. **(H-I)** UMAP representation of MGE types, colored by supertype and annotated with major supertype marker genes (**H**) or by region (**I**). This UMAP was calculated for cells in **G** based on 19 PCs derived from 4,963 DE genes. **(J)** Constellation plot of the global relatedness between MGE types, grouped into supertypes. Cluster labels colored green are clusters where more than 80% of cells are derived from HPF. **(K)** UMAP representation of MGE types, as in **H**, showing expression of select supertype marker genes in red. **(L)** RNA ISH for select markers expressed in the HPF-enriched MGE supertypes. Sst Ctsc is a supertype highly enriched in HPF, located mostly in CA3 (marked by *Rxfp3*) and the polymorphic layer of DG (DG-po) (marked by *Cxcr4*).

The current CTX-HPF GABAergic taxonomy is largely consistent with our previous transcriptomic taxonomy derived from two cortical areas, VISp-ALM (Tasic et al., 2018), at the supertype level but exhibits some notable differences at the type/cluster level (**Fig. S6A, S8**).

The current cell type taxonomy is also consistent with the large body of literature about the cortical GABAergic interneurons (Lim et al., 2018; Pelkey et al., 2017; Tremblay et al., 2016), as well as with a large-scale Patch-seq study described in our companion paper (Gouwens et al., https://www.biorxiv.org/content/10.1101/2020.02.03.932244v1).

In the CGE neighborhood, the Lamp5 subclass (mostly neurogliaform cells) is divided into 3 supertypes, Lhx6 (clusters #3-6), Prox1 (#7-16), and Pax6 (#17-19). The Sncg subclass is divided into a highly distinct supertype, Ndnf HPF (#20-24), and several more closely related supertypes, Ptprk (#25-28), Krt73 (#29-31, 37-38), Pax6 (#34-36), and Serpinf1 (#32-33, 39-41). The Vip subclass is divided into 2 supertypes, Car10 (#42-53) and Ptprk (#54-59) (**Fig. 2A-F, S9**). In the MGE neighborhood, the Sst Chodl subclass (#60-63) remains as one group (representing long-range projecting Sst cells). The Sst subclass is divided into 7 supertypes, Myh8 (#64-67), Crh (#68-73), Myh8 HPF (#74-78), Etv1 (#79-83), Nmbr (#84-89), Sfrp2 (#90-95), and Ctsc (#96-101). The Pvalb subclass is divided into 3 supertypes, Th (#102-108, which contains both *Sst*+ clusters and *Pvalb*+ clusters), Lpl (#109-118; containing basket and related cells), and the highly distinct Vipr2 (#119; representing chandelier cells) (**Fig. 2G-L, S10**).

As shown in dot plots (**Fig. 2A, G**) and UMAPs (**Fig. 2C, I**), most clusters are shared by all isocortical areas, consistent with our previous observations (Tasic et al., 2018), and also by retrohippocampal (RHP) regions. Even the estimated relative proportions of cells in these clusters, as revealed from the large number of 10xv2 cells, appear consistent across the different regions (**Fig. 2A, G**). At the same time, we also observed a set of clusters that are specific to or highly enriched in HPF. These include the Lamp5 Lhx6, Sncg Ndnf HPF, Sst Myh8 HPF, and Sst Ctsc supertypes, as well as select clusters within shared supertypes (i.e., Sncg Ptprk #27-28, Sncg Krt73 #37-38, Sst Etv1 #83, Sst Th #103-104, and Pvalb Lpl #117). Conversely, some clusters are largely absent from hippocampus (HIP; including Lamp5 Pax6, Sncg Serpinf1 and Sst Crh supertypes, and Sst Nmbr #87-89 and Sst Sfrp2 #90-93 clusters), while others appear to have CTX- or HIP-selective counterparts (e.g. Sst Myh8 in CTX versus Sst Myh8 HPF, Sst Etv1 #79-82 in CTX versus Sst Etv1 #83 in HIP). The greatest distinction in GABAergic interneuron type composition is between CTX and HIP itself. The RHP regions often contain both CTX and HIP clusters, with a few exceptions: Sst Myh8 HPF supertype is highly enriched in PPP-SP and HIP, Sst Ctsc #96-99 clusters are specific to HIP, and Pvalb Lpl #117 cluster is highly enriched in ENT.

Our CTX-HPF GABAergic taxonomy corresponds well with a previous scRNA-seq study of CA1 interneurons (Harris et al., 2018), particularly for some HPF-specific clusters identified here (**Fig. S6A**). In the CGE neighborhood, the HPF-enriched supertype Ndnf HPF is uniquely marked by *Ntng1* and does not express canonical pan-CGE marker *Prox1* nor its subclass markers *Vip, Sncg* or *Lamp5* (except for one cluster; **Fig. 2E, F, S9**). *Ntng1*+ cells in HIP are seen at the stratum radiatum/stratum lacunosum-moleculare (sr/slm) border, suggesting that they may be the trilaminar cells or radiatum-retrohippocampal neurons that project to RSP (Harris et al., 2018). The Lamp5 Lhx6 supertype may be neurogliaform cells that express typical MGE genes, *Lhx6* and *Nkx2.1*. Its cluster #3 includes both CTX and HPF cells, while clusters #4-6 are HPF specific and are marked by *Rxfp3* which is expressed specifically in CA3 (**Fig. 2E, F, S9**). Vip cells in HPF are largely intermixed with cortical Vip cells, except for cluster #53 marked by *Qrfpr*, which is expressed mainly in the stratum radiatum of CA3 (**Fig. S9**).

In the MGE neighborhood, the Sst subclass has multiple HPF-enriched clusters. Most of these clusters are marked by *Npffr1*, which is not expressed in cortical Sst cells. The HPF-enriched supertype Myh8 HPF and cluster Etv1 #83 are closely related to the cortical Myh8 and Etv1 supertypes that have few or no cells from HIP (**Fig. 2G-J**). Our Patch-seq study showed that the cortical Myh8 and Etv1 cells are L5 T-shaped Martinotti cells (Gouwens et al., companion manuscript submitted). Thus, the HPF Myh8 and Etv1 cells may correspond to the Martinotti-like oriens lacunosum-moleculare (OLM) cells with perpendicular axonal projections to the innermost layer (stratum lacunosum-moleculare, SLM) (Leão et al., 2012).

Sst Ctsc is a highly distinct HPF-specific supertype, which is located to CA3 and the polymorphic layer of DG (DG-po, also known as the hilus) (**Fig. 2K-L, S10**). Clusters #96-98 are marked by *Rxfp3* which is expressed in CA3, while clusters #99-100 are marked by *Cxcr4* expressed in DG-po. The *Cxcr4*+ cells (#99-100) may correspond to the hilar performant path-associated (HIPP) or DG somatostatin-expressing-interneurons (DG-SOMIs) described previously (Hosp et al., 2014; Myers and Scharfman, 2009; Raza et al., 2017; Savanthrapadian et al., 2014). Finer subtypes with different projection patterns and functional roles have been described (Yuan et al., 2017), which might be correlated with the transcriptomic types described here.

### Glutamatergic cell type taxonomy

The cell type landscape within the glutamatergic excitatory neuronal class is much more complex than the GABAergic class. To navigate this landscape, we first examined the major branches in a top-down direction within the glutamatergic portion of the taxonomy tree (**Fig. 1B, Table S3**). Two types that branch out before the main glutamatergic/GABAergic split are the highly distinct Cajal-Retzius cells (CR #1), and cells of the 3^rd^ ventricle (V3d). Excluding these two types, we defined 28 subclasses within the main glutamatergic branch and visualized them in a taxonomy tree (**Fig. 1B**), UMAP plots and a constellation plot (**Fig. 3A-C**), together with marker gene expression (**Fig. S11**).

**Figure 3.**
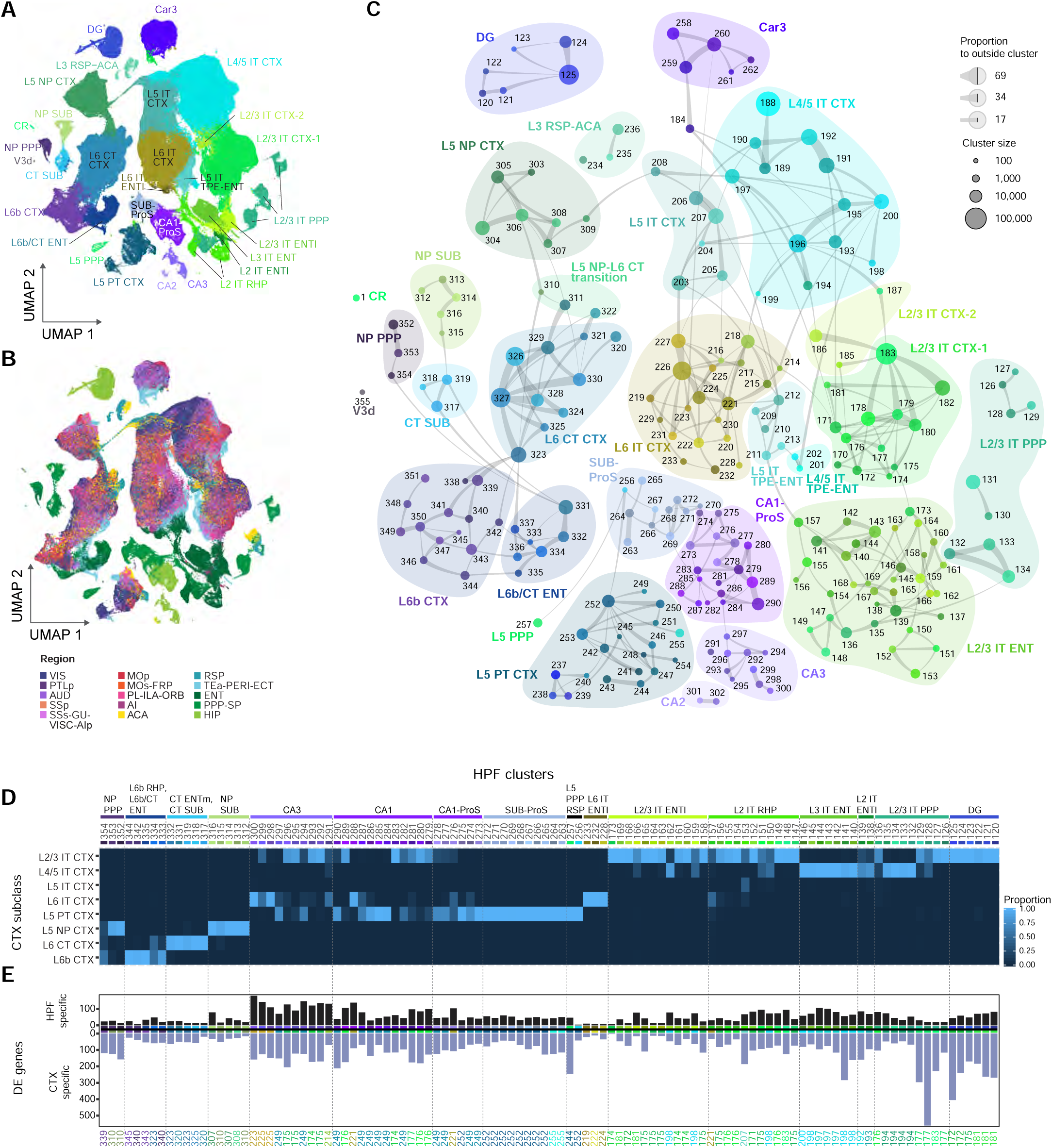
Glutamatergic cell types of CTX and HPF. **(A-B)** UMAP representation of glutamatergic types colored by subclass (**A**) or region (**B**). This UMAP was calculated for cells in the glutamatergic class based on 59 PCs derived from 4,963 DE genes (**Methods**). **(C)** Constellation plot of the global relatedness between glutamatergic types. Each cluster is represented by a dot, positioned at the cluster centroid in UMAP coordinates in **A**, with dot size proportional to the number of cells in the cluster. Connecting lines represent the proportion of nearest neighbors between pairs of clusters. Clusters are grouped into subclasses. **(D)** Correspondence of HPF clusters to CTX subclasses. Gene expression for each HPF cell (*n* = 1,781 genes) was correlated with average expression of each CTX cluster, in imputed common space. The CTX cluster with highest correlation was selected as the match, and the matches were aggregated by CTX subclass. The correspondence is represented as a proportion of total matches. **(E)** Same as in **D**, but the best-matched individual CTX cluster was selected, and the number of differentially expressed genes was calculated for the highest correlated HPF-CTX cluster pair.

The first branch within the main glutamatergic branch separates the DG granule cell subclass (clusters #120-125), and does not include any cortical counterparts, suggesting a highly distinct molecular nature of DG granule cells. The second major branch includes both CTX and HPF cells that are divided into 7 subclasses: CTX L5 near-projecting (NP), L6 corticothalamic (CT) and L6b neuron subclasses and related subclasses from HPF (clusters #303-354). The relatedness displayed within this branch suggests the existence of cell types in HPF regions homologous to these known isocortical deep-layer cell types. The third major branch comprises clusters that are specifically located in CA1, CA2, CA3, ProS and SUB and are divided into 4 subclasses, SUB-ProS (#263-272), CA1-ProS (#273-290), CA3 (#291-300), and CA2 (#301-302). The SUB-ProS and CA1-ProS subclasses both contain clusters from ProS, suggesting ProS contains cell types similar to either SUB or CA1, consistent with our previous study (Ding et al., https://www.biorxiv.org/content/10.1101/2019.12.14.876516v1). The next major branch represents the Car3 subclass (#258-262), which includes neurons from the claustrum (CLA) and from L6 of many lateral cortical areas. In our previous study, these *Car3*+ CLA and L6 neurons were shown to have extensive intracortical axon projections (Wang et al., https://www.biorxiv.org/content/10.1101/675280v1). The next major branch (#234-257) contains the subclass of CTX L5 pyramidal tract (PT) neurons (also known as subcerebral projection neurons, SCPNs) and two related subclasses. The final and largest branch (#126-233) is composed of intratelencephalic (IT) and related neuronal types from all layers of all CTX regions as well as RHP regions, again suggesting homology between isocortical IT neurons and many layer-specific neuronal types in the retrohippocampal regions. Within this major branch, 12 IT subclasses are defined that correspond well to specific layers (L2-6) and/or regions.

Given the existence of many distinct subclass branches for different hippocampal and retrohippocampal regions, some in parallel with corresponding isocortical subclasses, we further explored the relatedness between cell types from HPF (archicortex) and CTX (neocortex) by searching for gene expression covariation despite of regional difference. To do this, we separately defined DE genes within CTX or HPF, then intersected those gene sets to arrive at a common set of DE genes; we used these genes to correlate gene expression for each HPF cell to the average expression of each CTX cluster in an imputed common space (**Methods**). The CTX cluster with the highest correlation was selected as the match, and the matches were aggregated by CTX subclass (**Fig. 3D**). This approach revealed that most HPF cell types match a specific CTX subclass. We then calculated the number of DE genes between the highest correlated HPF-CTX cluster pair, which could indicate the overall degree of relatedness or similarity of the pair (the fewer DE genes, the more related; **Fig. 3E**).

We found that, based on the lower numbers of DE genes between the pairs, CTX NP, CT and L6b subclasses have the greatest similarity with their counterparts in ENT, PPP and SUB, so do CTX L2/3 and L6 IT subclasses with clusters in ENT (**Fig. 3E**). These similarities are consistent with our marker gene-based annotation of HPF clusters into corresponding layers, providing a mutual confirmation. Interestingly, L3 IT ENT (#140-146), L2 IT ENTl (#137-139) and part of L2/3 IT PPP (#132-135) were mapped to L4/5 IT CTX, revealing a novel relationship, which is supported by marker genes (for example, L2 IT ENTl expresses L4/5 markers *Endou* and *Grik1*, but not L2/3 marker *Cux2*). We further uncovered resemblance, albeit distant (i.e., many more DE genes, **Fig. 3E**), of SUB-ProS and HIP cell types to isocortical cell types. All SUB and ProS clusters are most related to L5 PT CTX (supported by marker genes *Bcl6*, *Pou3f1* and *Npr3*), so is cluster L5 PPP #257 (though more remotely). Intriguingly, individual clusters in both CA1 and CA3 are differentially mapped to 3 different cortical subclasses: L2/3 IT CTX, L6 IT CTX and L5 PT CTX. The implication of all these similarities is further investigated in a later section. Finally, DG granule cells are remotely related to L2/3 IT CTX as well.

### Continuous gene expression variation in glutamatergic neuron types across layers and regions of isocortex

We examined the composition of each glutamatergic cluster to find that most clusters predominantly contain cells derived from either CTX or HPF (**Fig. 1B, 3B, S12-13**). Examining cell distribution in different clusters one region at a time (**Fig. S12B**) showed that within CTX, sensory (i.e., AUD, VIS, SSp, SSs, PTLp), motor (i.e., MOp, MOs) and prefrontal (i.e., PL-ILA-ORB, MOs-FRP, AI) cortical areas have largely similar cell type compositions, whereas the lateral (i.e., TEa-PERI-ECT) and especially medial (i.e., ACA, RSP) cortical areas have additional unique cell types. Surprisingly, most CTX glutamatergic clusters contain cells from multiple or all cortical areas (**Fig. S13**), in contrast to the finding in our previous study focused on VISp and ALM (Tasic et al., 2018). This discrepancy could reflect a difference between discrete sampling of isocortex in the previous case where we may have sampled only the ends of one or more gene expression gradients, and now - by sampling cells from all intervening areas we reveal the gene expression continuity that prevents the separation of these cells into discrete types. To test this possibility, we investigated CTX IT and PT cell types in detail.

The isocortical IT neurons form 6 subclasses (**Fig. 1B, 3A, C**): L2/3 IT CTX-1 (clusters #170-183), L2/3 IT CTX-2 (#185-187), L4/5 IT CTX (#188-202), L5 IT CTX (#203-208), L5 IT TPE-ENT (#209-213), and L6 IT CTX (#214-231). We built a UMAP and a constellation plot (**Fig. 4A, D**) to represent gene expression-based relationships among cells in these subclasses, except for L5 IT TPE-ENT as it contains predominantly ENT cells. The UMAP revealed a gradual transition of the subclasses along the cortical depth, from L2/3 to L6. To further define this continuum, we computed one dimensional UMAP for all the IT cells based on the PCs in imputed space (**Methods**), which corresponded very well with cortical depth, and from this we calculated a pseudo-layer dimension and colored the IT UMAP according to this dimension **(Fig. 4B)**. The distribution of cells in each subclass along the pseudo-layer dimension showed that the subclasses fall along a gradient (**Fig. 4C**). Correspondence between the pseudo-layer dimension and actual cortical layers was established by calculating the expression of layer-specific marker genes (e.g., *Otof*, *Rspo1*, *Fezf2*, *Rxfp1, Osr1)* for cells ordered along the pseudo-layer dimension (**Fig. 4C right panel, S14A**). We found that collectively the clusters transition gradually across cortical depth from superficial to deep. There is a more discrete separation between L2/3 and L4/5 IT cell types, but L4/5, L5 and L6 IT types are distributed largely continuously (**Fig. 4C**). Similar observations were made when examining each isocortical region individually (**Fig. S14B**).

**Figure 4.**
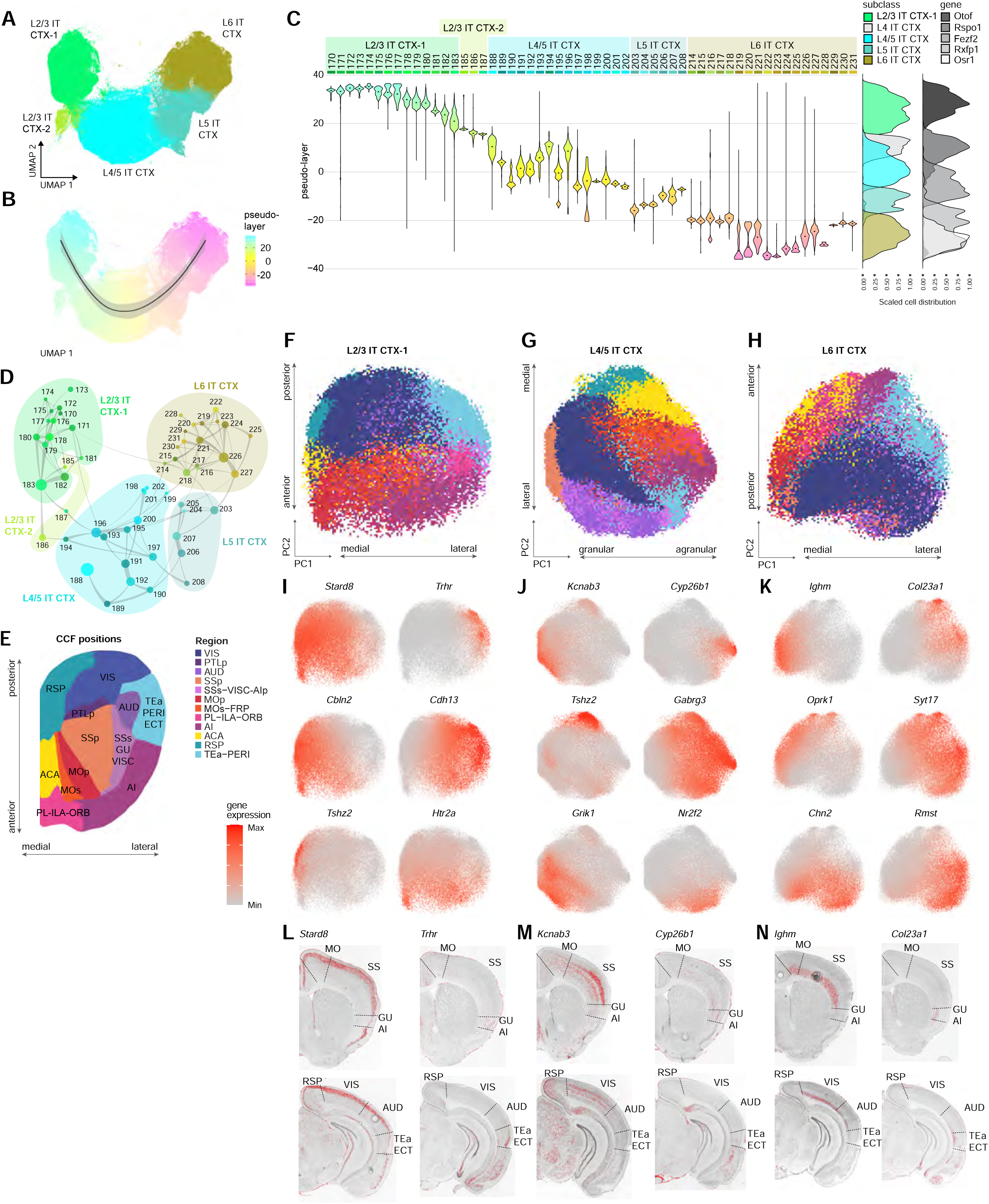
Multi-dimensional gradients of the distribution of IT cell types in isocortex. **(A)** UMAP representation of CTX IT cells colored by subclass reveals a prominent 1^st^ UMAP dimension that corresponds to cortical depth. The UMAP was calculated based on 59 PCs derived from 4,963 DE genes. CTX IT cells included: 10xv2: *n* = 499,190; SSv4: *n* = 25,355. **(B)** The same 2D UMAP as in **A** colored by pseudo-layer dimension, which corresponds to 1D UMAP dimension based on 59 PCs in imputed space for cells and genes in **A** (**Methods**). The black line represents the best fit with 95% confidence interval (gray). **(C)** The distribution of cells for each CTX IT cluster along the pseudo-layer dimension defined in **B**, visualized by a violin plot (dots are medians). The two density plots on the right represent the distribution of cells from IT subclasses or the expression of select layer-specific marker genes along the same pseudo-layer dimension, normalized for each subclass or gene, respectively. **(D)** Constellation plot of the global relatedness between CTX IT types. Each cluster is represented by a dot, positioned at the cluster centroid in UMAP coordinates in **A**, with dot size proportional to the number of cells in the cluster. Clusters are grouped by subclass. **(E) A** 2D flatmap representation of isocortical regions according to their positions in CCFv3. **(F)** PCA plot of L2/3 IT CTX**-**1 subclass cells based on 349 genes selected to highlight regional differences (**Methods**). To make the plot more visually interpretable, the cells were binned into 100 equal bins along each PC. The bins were colored based on the region with the highest representation within a given bin. PC1 and PC2 mostly correspond to medial-lateral and anterior-posterior spatial dimensions. L2/3 IT CTX-1 cells included: 10xv2: *n* = 115,215; SSv4: *n* = 5,857. **(G)** PCA plot of L4/5 IT CTX subclass cells based on 556 genes, selected to highlight regional differences (**Methods**), with cells binned like in **F**. PC1 corresponds to a granular-agranular regional transition and PC2 mostly corresponds to a lateral-medial spatial dimension. L4/5 IT CTX-1 cells included: 10xv2: *n* = 253,399; SSv4: *n* = 11,490. **(H)** PCA plot of L6 IT CTX subclass cells based on 497 genes, selected to highlight regional differences (**Methods**), with cells binned like in **F**. PC1 and PC2 mostly correspond to medial-lateral and anterior-posterior spatial dimensions. L6 IT CTX cells included: 10xv2: *n* = 78,702; SSv4: *n* = 4,980. **(I-K)** Expression gradients (with no binning) of selected genes corresponding to the above PCA plots (**I**, **J** and **K** correspond to **F**, **G** and **H**, respectively). **(L)** RNA ISH images for genes shown in the top row of panel **I**, highlighting the regional distribution of cells in the L2/3 IT CTX-1 subclass. **(M)** RNA ISH images for genes shown in the top row of panel **J**, highlighting the regional distribution of cells in the L4/5 IT CTX subclass. **(N)** RNA ISH images for genes shown in the top row of panel **K**, highlighting the regional distribution of cells in the L6 IT CTX subclass.

The number of cells in each cluster varies dramatically, and there is often a dominant cluster within each subclass, e.g. cluster #183 in L2/3 IT, #188 in L4/5 IT and #226 in L6 IT (**Fig. S13**). Cluster #188 is the layer 4 specific cell type marked by gene *Rspo1*, presumably representing the granular layer containing spiny stellate or star pyramid neurons. Nearly all clusters contain cells from multiple cortical areas. Although enrichment to specific areas is seen for some clusters, there is no one-to-one correspondence between clusters and regions. To investigate global gene expression variation across all cortical regions (**Fig. 4E**), we performed principal component analysis (PCA) on all cells within an IT subclass based on genes selected to highlight regional differences (**Methods**). We found that the first two PCs defined this way correspond to medial-lateral and anterior-posterior spatial dimensions for the L2/3 IT (**Fig. 4F**) and L6 IT (**Fig. 4H**) subclasses. For the L4/5 IT subclass, they correspond to L4-to-L5 (granular-agranular) and medial-lateral dimensions (**Fig. 4G**). Genes that contribute to these PCs have various gradient expressions across the cortical sheet and corresponding *in situ* patterns from the RNA ISH data (**Fig. 4I-N**). We did not include the analysis for the L5 IT subclass as the main axes of gene expression variation corresponded to cluster diversity instead of regional gradients (data not shown). Overall, our results show that isocortical IT cell types are often shared among multiple cortical areas and can exhibit gradient gene expression variations across these areas.

The cortical PT neurons (marked by *Bcl6*) could be segregated into three region-specific subclasses, L3 RSP-ACA (#234-236, marked by *Scnn1a*), L5 PT CTX (#237-256, marked by *Fam84b*) and L5 PPP (#257, marked by *Ptgfr*) (**Fig. 5**). Within the L5 PT CTX subclass, individual types or supertypes also show differential regional enrichment (**Fig. 5A-D**). The Chrna6 supertype (#237-239) is enriched in posterior sensory areas. In the Cdh13 supertype (#249-254), cluster #249 is highly specific to medial anterior cortical areas, PL and ILA, based on marker genes *Ndnf* and *Nnat* (**Fig. 5D-E**). The biggest cluster, #252, includes cells from most cortical areas. By applying the same analysis as described in **Fig. 4F-H**, we identified regional diversity within this cluster with the top two PCs corresponding to medial-lateral and anterior-posterior spatial axes (**Fig. 5F**), as well as the key genes driving this diversity (**Fig. 5G**). Within the C1ql2 supertype (#255-256), cluster #255 is enriched in RSP-ACA, whereas #256 is located in the posterior part of RSP bordering PPP based on co-expression of *C1ql2* and *Nnat* (**Fig. 5D-E**). The three ALM L5 PT types identified in our previous studies (Economo et al., 2018; Tasic et al., 2018) are contained in clusters #246-247 (thalamus-projecting cells) in the Npnt supertype (#240-248) and cluster #250 (medulla-projecting cells) in the Cdh13 supertype, respectively. It will be interesting to see if cells in other cortical areas belonging to these two supertypes have similarly differential projection patterns.

**Figure 5.**
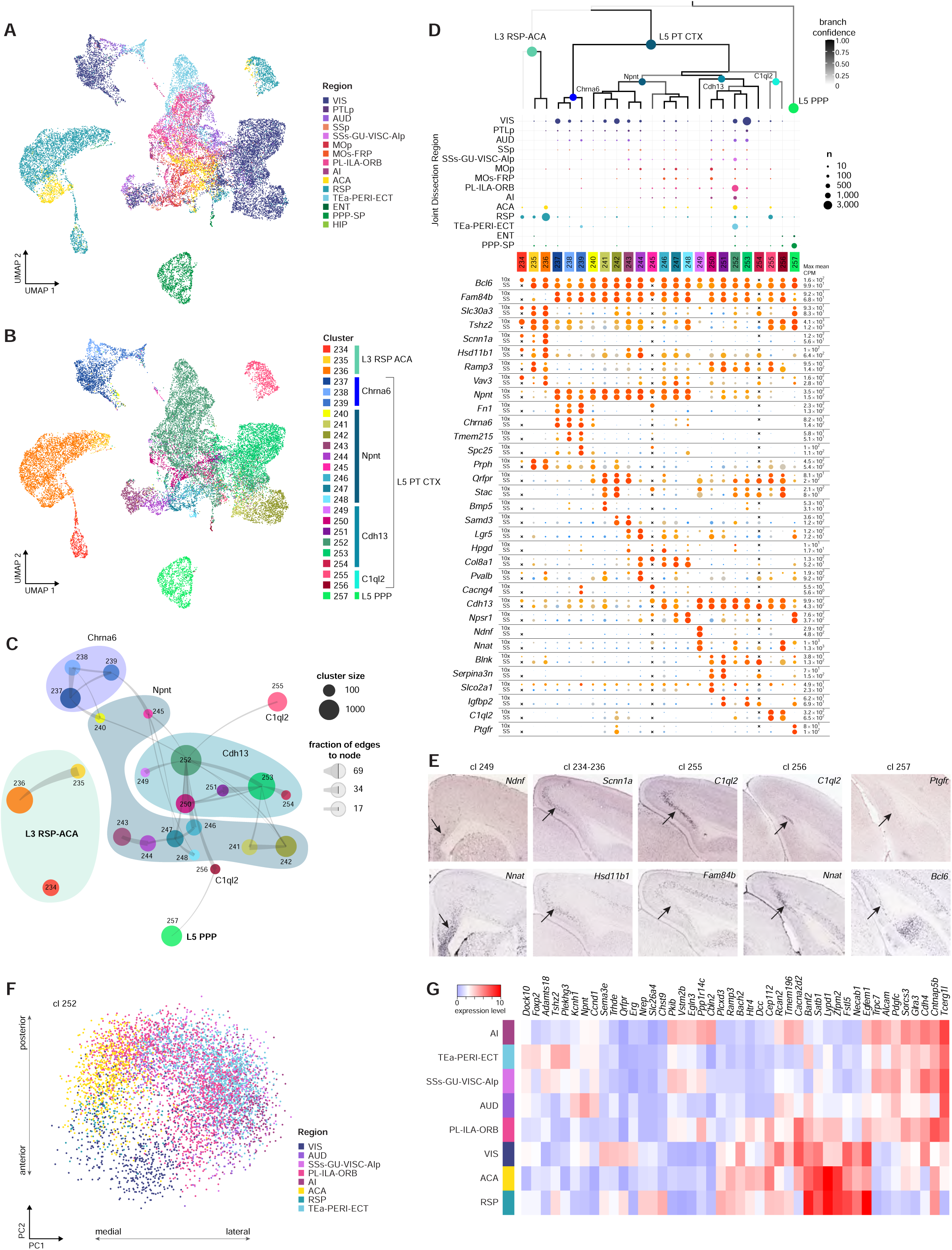
L5 PT and related cell types in isocortex. **(A-B)** UMAP representation of cells in the PT neighborhood colored by brain region (**A**) or cluster (**B**). This UMAP was calculated using 14 PCs based on 4,963 DE genes. Cells included: 10xv2: *n* = 22,237; SSv4: *n* = 2,221. **(C)** Constellation plot of the global relatedness between L5 PT and related types. Each cluster is represented by a dot, positioned at the cluster centroid in UMAP coordinates in **A-B**. Boundaries are drawn for L3 RSP-ACA subclass and supertypes in L5 PT CTX subclass, each labeled by their shared marker gene. **(D)** Dendrogram of 20 PT types and 4 PT-like types followed by the number of cells (dot size) within each cluster derived from each region noted on the left. The dendrogram is a part of the dendrogram in Fig. 1B. Below the regional dot plot, the gene expression dot plot illustrates marker gene expression in each cluster from 10x (top row) and SSv4 (bottom row) datasets. Dot size and color indicate proportion of expressing cells and average expression level in each cluster, respectively. **(E)** RNA ISH images for prominent cluster marker genes. Markers for clusters #255-257 highlight the transition of clusters from RSP to PPP. **(F)** PC plot of cells within cluster #252, colored by brain region. The first two PCs correspond to medial-lateral and anterior-posterior gradients. **(G)** Heatmap of genes associated with the top two PCs driving the diversity within cluster #252. Expression of genes is split by region.

L3 RSP-ACA (#234-236, marked by *Scnn1a*) is an unusual subclass. It expresses PT marker gene *Bcl6* but not *Fam84b*; it also expresses a pan-IT marker *Slc30a3*, as well as L4 IT-specific markers *Rspo1* and *Scnn1a* but not *Rorb* (**Fig. 5D**). RNA ISH images show that the *Scnn1a*+ L3 RSP-ACA subclass is more superficial than the L5 PT C1ql2 supertype in RSP (**Fig. 5E**). Projection mapping experiments in Allen Mouse Brain Connectivity Atlas (http://connectivity.brain-map.org/; experiments 166269090, 166458363 and 181860879) (Oh et al., 2014) showed that neurons labeled via Scnn1a-Tg3-Cre transgene in RSP have long-range projections to both cortical and subcortical targets. The gene expression makeup, layer specificity and projection pattern altogether suggests that the L3 RSP-ACA subclass has an intermediate cell type identity between IT and PT.

### Glutamatergic cell types in hippocampal formation in comparison with those in isocortex

As mentioned above, the glutamatergic cell types in different regions of HPF appear to be discretely different from those in CTX and from each other, while at the same time they also demonstrate correspondence with CTX glutamatergic subclasses (**Fig. 1B, 3**). Here we investigate them in further detail in three separate groups, IT, NP/CT/L6b and SUB/HIP, and uncover striking correlation between molecular/transcriptomic organization and spatial/anatomical organization of these cell types.

As shown in the UMAP containing all putative IT-projecting cells from cortex and hippocampus (**Fig. 6A-B**), CTX IT cell types are more similar to each other and aggregate in the middle of the UMAP, while HPF IT cell types are separated into two distinct groups, one mainly for ENT and the other mainly for PPP. In the ENT IT group (**Fig. 6C-D, F-G**), the ENT IT cell types are arranged in a layer-selective manner, in the order of L2, L2/3, L3, L5 and L6, consistent with their correspondence with the CTX L2/3-L6 IT cell types, and they show further differential distribution along the anterior-posterior axis. The spatial distribution (**Fig. 6D, G**) starts with the L3 IT ENT subclass in the caudal part, including the Plch1 Fn1 supertype (clusters #140-143) specific to ENTm and the Fign supertype (#144-146) specific to ENTl. Then it transitions to the L2/3 IT ENTl subclass marked by *Penk* in the middle part of ENTl, including the Fign supertype (#158-162) specific to L3 ENTl and the Penk Ndst4 supertype (#163-169) specific to L2/3 ENTl. The L2 IT RHP subclass is split into 3 supertypes, with the Lef1 supertype (#150-153) being L2 ENTm cells, the Dcn Pcsk5 supertype (#155-157) assigned to the superficial layer of ProS and HATA regions (Ding et al., https://www.biorxiv.org/content/10.1101/2019.12.14.876516v1), and the Cfap58 supertype (#147-149, 154) located in PAR (marked by *Calb1*, **Fig. 6G**). The L2 IT ENTl subclass (#137-139) marked by *Grik1* is located at the border between L1 and L2 in the rostral ENTl and likely continues into the piriform cortex. The L6 IT ENTl subclass (#232-233) marked by *Dlk1* is specifically located in L6 of the rostral and middle parts of ENTl. Finally, the L5 IT TPE-ENT subclass (#209-213) marked by *Dcn* contains L5 cells in ENTl and caudal ventral part of ENTm.

**Figure 6.**
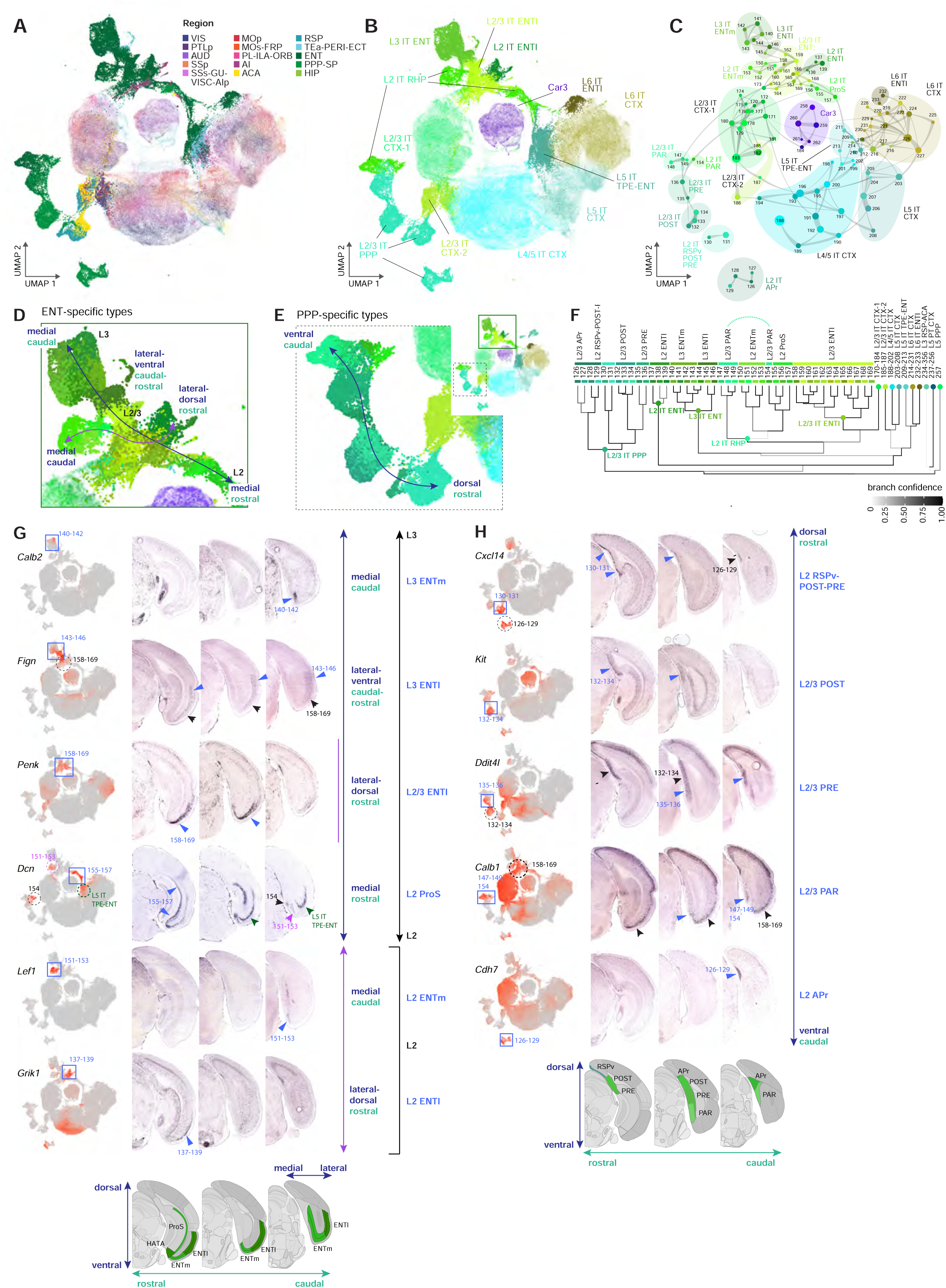
Transcriptomic relationship and anatomical distribution of IT-like cell types in retrohippocampal regions. **(A-B)** UMAP representation of IT neurons from CTX and HPF, colored by region (**A**), with non-HPF neurons faded out, or by subclass (**B**). The Car3 subclass contains cells from both CTX and claustrum (CLA). This UMAP was calculated using 59 PCs based on 4,963 DE genes. Cells included: 10xv2: *n* =375,658; SSv4: *n* =47,813. **(C)** Constellation plot of the global relatedness between IT types from CTX and HPF. Each cluster is represented by a dot, positioned at the cluster centroid in UMAP coordinates in **A-B**. Clusters are grouped by subclass or supertype. **(D)** Enlarged view of UMAP representation in **A** of ENT-specific types colored by cluster. The blue arrow marks a medial-rostral to medial-caudal trajectory from L2, L2/3, to L3 of ENT clusters. The purple arrow marks a medial-caudal to lateral-dorsal trajectory of ENT clusters. **(E)** Enlarged view of UMAP representation in **A** of PPP-specific types colored by cluster. Blue arrow marks rostral-dorsal to caudal-ventral trajectory of clusters within L2/3 IT PPP. **(F)** Dendrogram of ENT-PPP clusters with branches annotated by subclass and supertype. **(G)** Anatomical annotation of supertypes showing relationships of various ENTl and ENTm supertypes marked in panel **D**. UMAP representations, as in panel **A**, show expression of select supertype marker genes in red. RNA ISH images of supertype marker genes along 3 rostral to caudal sections reveal specific locations of the different supertypes. Colored boxes/circles mark expression of gene marker in UMAP space and match colored arrowheads marking location in RNA ISH images. **(H)** Anatomical annotation of supertypes showing PPP transition marked in panel **E**. UMAP representations, as in panel **A**, show expression of select supertype marker genes in red. RNA ISH images of supertype marker genes along 3 rostral to caudal sections reveal specific locations of the different supertypes and how they transition from RSPv through PPP. Colored boxes/circles mark expression of gene marker in UMAP space and match colored arrowheads marking location in RNA ISH images.

The PPP IT group (**Fig. 6C, E-F, H**) contains mainly the L2/3 IT PPP subclass (clusters #126-136) and is most closely related to the unique L2 IT RSP-ACA supertype (#185-186) in the L2/3 IT CTX-2 subclass, consistent with the anatomical proximity of RSP and PPP. The supertypes within L2/3 IT PPP follow a rostral dorsal to caudal ventral transition (**Fig. 6E, H**), starting with the Pdlim1 supertype (#130-131) in L2 of RSPv, POST and PRE, then the Kit supertype (#132-134) in L2/3 of POST, and the Wfs1 Prlr supertype (#135-136) in L2/3 of PRE, which is followed by the Cfap58 supertype (#147-149, 154) in PAR. The Cdh7 supertype (#126-129) is specific to L2/3 of Apr.

The NP/CT/L6b neighborhood contains paired sets of L5 NP, L6 CT and L6b subclasses for CTX and HPF (**Fig. 7**). The L5 NP CTX subclass (clusters #303-311) is closely related to the NP SUB subclass (#312-316, marked by *Ly6g6e*) and more remotely related to the NP PPP subclass (#352-354, marked by *Nts*). The L6 CT CTX subclass (#320-330) is closely related to the CT SUB subclass (#317-319, marked by *Rorb*) and also related to the Col5a1 supertype (#331-332, L6 CT ENTm clusters) of the L6 CT L6b ENT subclass. The L6b CTX subclass (#338-351) also contains cells from ENT, PPP and SUB in most of its clusters, including several ENT-enriched clusters (i.e. #342, 344), and it is related to the Cplx3 supertype (#333-337) of the L6 CT L6b ENT subclass. In addition, we observed a parallel continuous transition between L6 CT and L6b cell types for both CTX and ENT (**Fig. 7A-C**). As seen in the IT cells (**Fig. 4, 6**), the CTX L5 NP, L6 CT and L6b clusters are largely shared across all cortical areas (**Fig. 7B, S13**), with the exception of several ACA-RSP enriched clusters (i.e. #307-310 in the NP subclass, and #320-322 in the CT subclass), whereas the HPF NP and CT cell types are highly distinct among SUB, ENT and PPP. We identified multiple marker genes confirming the regional specificity of these HPF subclasses (**Fig. 7D-E**).

**Figure 7.**
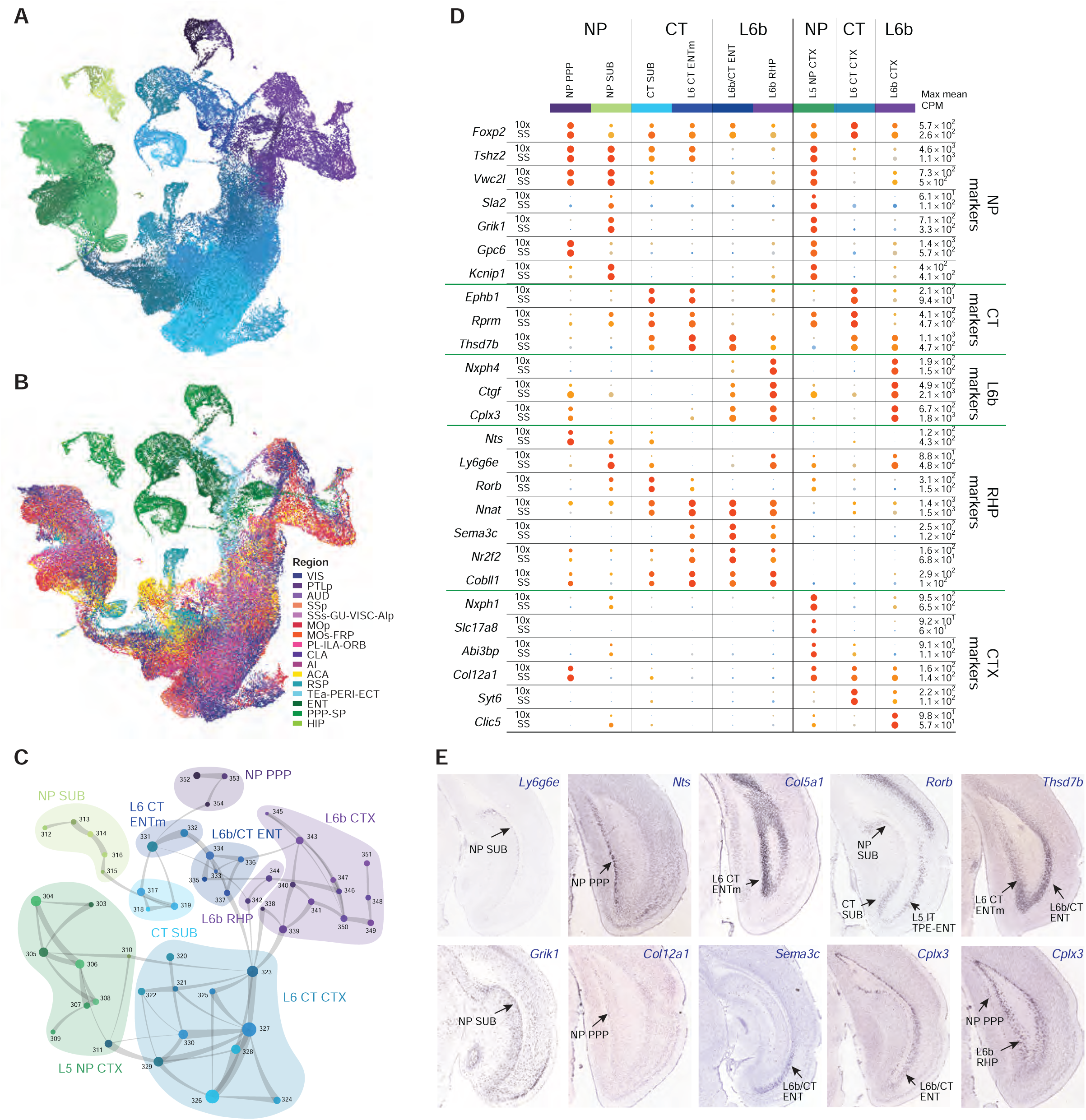
Parallel sets of NP CT L6b related cell types in isocortex and hippocampal formation. **(A-B)** UMAP representation of the NP/CT/L6b neighborhood from CTX and HPF, colored by cluster (**A**) or region (**B**). This UMAP was calculated using 17 PCs based on 4,963 DE genes. Cells included: 10xv2: *n* = 101,486; SSv4: *n* =12,019. **(C)** Constellation plot of the global relatedness between the NP/CT/L6b clusters. Each cluster is represented by a dot, positioned at the cluster centroid in UMAP coordinates in **A**. Clusters are grouped by subclass or supertype. **(D)** Dot plot illustrating marker gene expression in corresponding CTX and HPF cell types for 10xv2 (top row) and SSv4 (bottom row) datasets. Dot size and color indicate proportion of expressing cells and average expression level in each subclass, respectively. **(E)** RNA ISH images of supertype marker genes marking the specific locations of NP, CT and L6b cell types in ENT, PPP and SUB.

### Multidimensional variation of cell type distribution in the hippocampal and subicular regions

UMAP and constellation plots for the SUB/HIP neighborhood, including SUB, ProS, CA1, CA2, CA3 and DG regions (**Fig. 8A-D**), show that the DG, CA2 and CA3 subclasses are highly distinct, whereas the SUB-ProS and CA1-ProS subclasses together largely form a continuum. In the CA3 subclass, we found that the distinct cluster #291 represents hilar mossy cells (Scharfman and Myers, 2012) based on multiple marker genes (e.g. *Gal, Rgs12, Glipr1*, *Necab1*, *Calb2*) (**Fig. S15**).

**Figure 8.**
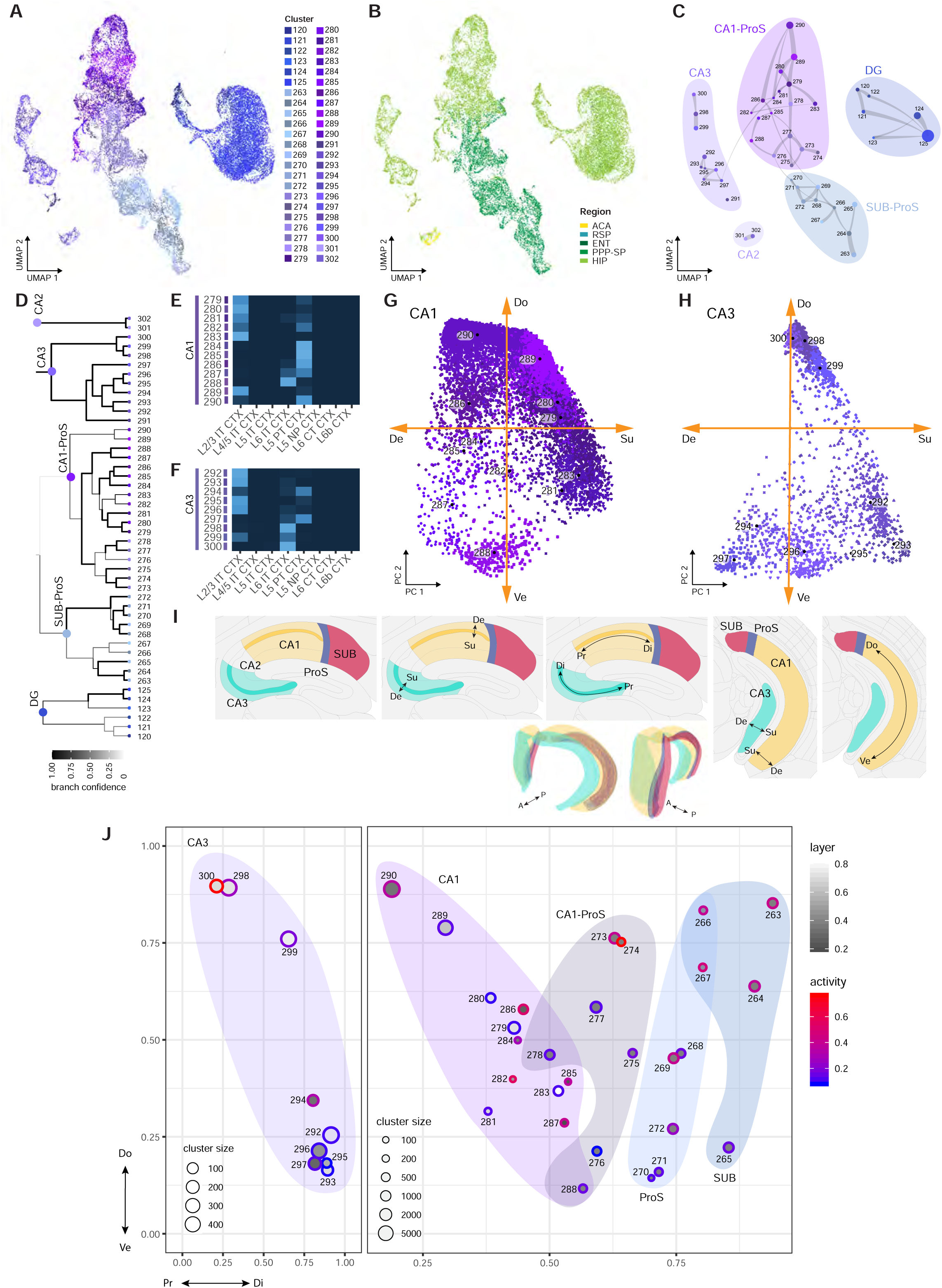
Multi-dimensional distribution of glutamatergic cell types in the hippocampus and subiculum. **(A-B)** UMAP representation of the SUB/HIP neighborhood, colored by cluster (**A**) or region (**B**). The UMAP was calculated using 17 PCs based on 4,963 DE genes. Cells included: 10xv2: *n* = 23,626; SSv4: *n* = 3,761. **(C)** Constellation plot of the global relatedness between the SUB/HIP clusters. Each cell type is represented by a dot, positioned at the cluster centroid in UMAP coordinates in **A**. Clusters are grouped by subclass. **(D)** Dendrogram of 46 clusters in the SUB/HIP neighborhood. **(E-F)** Correspondence of CA1 (**E**) or CA3 (**F**) clusters to CTX subclasses (part of Fig. 3D). **(G-H)** 2D PCA plots for cells in CA1 (**G**) or CA3 (**H**) clusters. The two axes correspond to the dorsal-ventral (Do-Ve) and superficial-deep (Su-De) dimensions, respectively. Each cell is colored by its cluster membership using different symbols, and cluster centroids are shown as a black dot next to each cluster number. **(I)** Structure of the mouse hippocampal and subicular regions. Images are rendered from Allen CCFv3. The 2D schematics show coronal and sagittal sections marking the spatial dimensions within the hippocampus and subiculum: proximal–distal (Pr–Di), superficial–deep (Su–De), and the dorsal–ventral (Do–Ve). **(J)** Summary of Pr-Di, Su-De, Do-Ve and activity dimensions at the cluster level for CA3, CA1, ProS and SUB. Each circle represents a cluster, for which we computed the average values for its cell members along each of the four dimensions: X axis for the Pr-Di dimension, Y axis for Do-Ve, filling color for Su-De (light colors correspond to the upper layer, and dark colors to the deep layer; only computed for cells in CA1 and CA3), and ring color for activity (red activated, blue not activated).

The DG subclass, marked by *Prox1*, has a dominant cluster, #125, which contains the vast majority of the granule cells (**Fig. S13**). Clusters #120-123 express a number of immediate early genes (IEGs), such as *Tmem2*, *Bdnf*, *Gadd45b*, *Arc*, and *Fosb*, suggesting they contain high-activity cells. Clusters #123-124, marked by *Hap1*, *Nr2f2* and *Trhr*, appear to be enriched in posterior/ventral DG. We did not find clusters strongly related to adult neurogenesis in DG (Gonçalves et al., 2016), indicating that immature neurons or progenitors might not be well labeled by the pan-neuronal or pan-glutamatergic Cre lines we used for this study. The CA2 subclass (#301-302) also contains cells from the ACA region. Marker gene expression of *Ntf3*, *Ramp3*, etc., shows that cells from the small structures, IG (underneath ACA) and FC, also belong to these clusters (**Fig. S15**).

Previous studies showed that CA1, CA3 and SUB all have gradual gene expression and connectivity changes (either gradient or discrete depending on the study) along multiple dimensions – superficial-deep, proximal-distal and dorsal-ventral (Cembrowski and Spruston, 2019). To understand the patterns of variation among the CA/SUB cell types identified in our study and their correlation with previously described dimensions, we performed UMAP and PCA. First, we noticed that there appeared to be one main axis that drove CA1, ProS and SUB variation. To extract this axis, we computed one dimensional UMAP of all the cells in the CA1-ProS and SUB-ProS subclasses. The distribution of cells along this axis for each cluster is shown in **Fig. S16A**, in which clusters are sorted based on the average values. Then we computed genes that either strongly correlate with this axis, which specify two ends of the spectrum, or whose expression is confined within a narrow range in the middle of the spectrum (**Fig. S16A**). ISH images of the selected genes (**Fig. S16B**) indicate that this axis corresponds to proximal-distal gradient, where the cell types transition from the proximal end of CA1 (marked by *Lct*), to distal CA1, CA1/ProS transition zone (marked by *Glipr*, *Dlk1*, *Dcn*), ProS (marked by *S100b*, *Klhl1*), and finally to distal SUB (marked by *Fn1*, *Cyp26b1*).

To examine other axes of variation, we performed PCA for CA1 and CA3 (excluding the hilar mossy cell cluster) separately. In each case, we observed that the top PC corresponded to a dorsal-ventral gradient (**Fig. 8G, H**). Closer examination revealed significant overlap between the genes that specify this gradient in either region (**Methods**), which we hypothesize is the core program for dorsal-ventral gradient specification. Using this core set of genes, we computed the top PC for cells in both regions and found it highly concordant with the original top PC for each region separately. We used the same gene set to compute the top PC in SUB/ProS and DG regions and again observed segregation of clusters along this gradient, which also corresponds to dorsal-ventral axis in SUB/ProS/DG based on ISH of key marker genes. The distribution of cells from all regions along this axis for each cluster is shown in **Fig. S16C,** segregated by subclass, along with a subset of genes in the core program (**Fig. S16C**) and ISH images of selected genes that are dorsal or ventral specific in some but not all the regions (**Fig. S16D**). Notably, *Nr2f2* is one of the most conserved ventral-specific genes across all the HPF regions and is also enriched in the TEa region (located at the posterior ventral part of the isocortex) for IT cells. As a transcription factor previously known to be involved in regulation of differentiation and patterning (Polvani et al., 2019), it is possible that *Nr2f2* is a master regulator for specifying dorsal-ventral variation in both cortical and hippocampal development. We also identified two members of the Wnt signaling pathway, *Wnt7b* and *Wnt2*, to be ventral and dorsal specific respectively, suggesting a potential role of this pathway in specifying dorsal-ventral axis in hippocampus. Besides the genes in the core program, each region also has a specific set of genes that contribute to this axis (**Fig. S16C**), e.g., *Coch* is ventral-specific only in CA3, and *Gpc3* only in CA1/ProS (likely in HATA), while *Rxfp1* and *Elfn1* are dorsal-specific only in SUB/ProS, *Wfs1* in CA1, and *Rph3a* in CA3. All the genes shown in the heatmap have ISH images from ABA that support their specificity along this axis, though only a subset of them are shown in **Fig. S16D**.

In both CA1 and CA3, we also observed that key genes contributing to the second PC correspond to the superficial-deep radial axis (**Fig. 8G-H, S17**). For example, in CA1, *Nptx2*, *Sulf2*, *Lpl* are expressed in the deep layer, while *Pde11a*, *Anln* are in the superficial layer (**Fig. S17A-C**). Consistent with the spatial layer distribution, the superficial CA1 cell types (#279-281, 283, 289) are most similar to L2/3 IT types, while deep CA1 cell types are most similar to L5 PT (for clusters #284-287, 290) or L6 IT (for #288) types (**Fig. 8E**). Similarly, in CA3, we identified a set of markers including *St18*, *Hopx*, *Sgcd*, *Prss23*, which are expressed in a very thin deep layer, while *Nos1*, *Kctd4*, *Kcnq5* have complementary expression pattern and label most of the CA3 cells (**Fig. S17D-F**). The separation of layer markers is more prominent in the ventral part of CA3. Clusters #294 and 297 located at the deep layer are most similar to L5 PT CTX cells, while the other clusters at the ventral superficial layer (#292, 293, 295, 296) are most similar to L2/3 IT CTX (**Fig. 8F**). Thus in both CA1 and CA3, superficial and deep clusters are similar to L2/3 IT CTX and L5 PT CTX, respectively. This finding supports the notion of an inside-out radial migration for the development of CA1 and CA3 pyramidal neurons similar to that in the isocortex (Khalaf-Nazzal and Francis, 2013).

The dorsal-ventral distribution of CA1 and CA3 clusters follow the taxonomy branches well (**Fig. 8D, G-H**). In CA1, clusters #289-290 are in the most dorsal location, followed by #279-286, and #287-288 most ventral. In CA3, clusters #298-300 (supertype Iyd) are in the dorsal location, and #292-297 more ventral. In the less diverse part of CA1 (most ventral) and CA3 (most dorsal), the superficial-deep distinction is also less obvious; correlated with this intriguingly, we note that in these parts, cluster #288 in ventral CA1 and clusters #298-300 in dorsal CA3 are all related to L6 IT CTX.

Besides dorsal-ventral, proximal-distal, and superficial-deep gradients, we also observed an activity-dependent transcriptional signature shared across all DG/CA3/CA1/ProS/SUB cell types (**Fig. S16E-F**). It includes many well-established IEGs such as *Ier5*, *Arc*, *Fos*, *Egr4* and *Nr4a1*, which have been shown to label neuronal ensembles encoding memory traces (Minatohara et al., 2015). We also identified activity-dependent genes co-expressed with IEGs in a cell type-dependent manner. For example, *Gadd45b* is expressed highest in DG and was shown to be required for activity-induced DNA demethylation of genes critical for adult neurogenesis (Ma et al., 2009).

To better understand the relationship between all the hippocampal and subicular clusters and the three-dimensional spatial structure of these regions (**Fig. 8I**), we plotted the average values for each cluster along all four dimensions of variation together, X-axis for the proximal-distal dimension, Y-axis for dorsal-ventral, color scale (gray, fill-in) for superficial-deep, and color scale (red-blue, outline) for activity (**Fig. 8J, Methods**). This summary plot shows how clusters differ along each dimension and the relationships between different dimensions. For example, while there is a large divergence between the dorsal clusters of SUB and CA1 along the proximal-distal axis, there is a convergence at the ventral parts of CA1 and ProS (and the HATA area). Also notably, the most dorsal clusters in CA1, CA3 and SUB show high levels of activity-dependent gene expression, whereas the most ventral ends of these regions show low levels (**Fig. S16E-F**). This distribution is consistent with a previous study of spatial distribution of *Arc* gene expression in the hippocampus, and could be associated with the differential response to spatial/nonspatial information along the dorsal-ventral axis (Chawla et al., 2018). Note that some genes show expression variation along multiple axes, e.g., some layer specific markers also vary in dorso-ventral direction, making it impossible to deconvolve their contribution to each axis, and our estimate of spatial distribution for each cluster should be validated by future spatial transcriptomic studies.

## DISCUSSION

In this study, we have conducted a large-scale and in-depth single-cell transcriptomic profiling of the entire isocortex and hippocampal formation in the adult mouse. This unprecedented dataset revealed remarkable cellular diversity, led to the generation of a catalog of transcriptomic cell types for all the regions contained within these two major brain structures, and allowed us to compare these cell types in various ways and uncover a new set of organizing principles.

One main goal in describing transcriptomic cell type diversity involves the deconvolution of various axes of gene expression variation, followed by discretization of cells into types, when possible. As previously observed in the cortex (Tasic et al., 2016, 2018), gene expression variation along these axes is frequently a combination of discreteness and continuity, likely reflecting the multiple facets of the cells’ identities, states and interrelationships. Here, for the first time, all regions of the cortex and hippocampus have been profiled in a systematic and standardized manner. This has enabled us to define both discrete and continuous axes of variation across the complete spatial landscape without significant gaps. We have thus identified many different patterns of cell type diversity.

At the molecular level, cell types can be organized in a hierarchical manner and have multi-scale and multidimensional relatedness with each other. Such molecular relationships correlate strongly with the spatial origins (both location and layer) of the cell types. Glutamatergic neuron types are much more diverse than GABAergic neuron types, both molecularly and spatially.

Some cell types are highly specific to a region, layer or location, while others are widely distributed and shared among multiple regions. The shared cell types can display gradients of gene expression variations across regions in various axes; such axes are consistent across different subclasses, suggesting a common underlying organizational plan. Within a region or within related regions, multiple cell types are organized in a coordinated manner, likely for the purpose of forming ordered functional circuits. Although highly distinct from each other, cell types from the hippocampal formation and isocortex also display remarkable homologous relationships through common gene expression variations and correspondences in spatial (laminar) specificity. Thus, evolutionary and developmental relationships between regions and cell types may also be inferred from transcriptomic cell type relationships.

Our previous VISp-ALM transcriptomic study had shown that GABAergic interneuron types are shared while glutamatergic neuron types are distinct between the two cortical areas, and we hypothesized that this dichotomy might be due to the differential developmental origins of these two classes of neurons (Tasic et al., 2018). During development, glutamatergic neurons are born within the ventricular and subventricular zones underneath the cortical plate, which is laid out into a protomap by a gradient or compartmentalized gene expression (Clowry et al., 2018; Rakic, 2009). In contrast, the GABAergic interneurons originate from ganglionic eminences in the subpallium, part of the ventral telencephalon, and migrate into cortex in tangential streams to populate all cortical regions (Lim et al., 2018).

Our study here extends the previous findings for both GABAergic and glutamatergic types. We show that GABAergic interneuron types are shared across all isocortical areas as well as retrohippocampal regions; most isocortical GABAergic types are also found in hippocampal regions (**Fig. 2**). We also find a subset of GABAergic types that are specific to HPF (**Fig. 2**). For example, the Ndnf HPF supertype may correspond to the trilaminar cells or the radiatum-retrohippocampal neurons projecting to RSP (Harris et al., 2018). The Sst Myh8 HPF supertype likely corresponds to the OLM cells (Leão et al., 2012), the hippocampal counterpart of cortical Martinotti cells, whereas one of the Sst Ctsc types may correspond to the HIPP cells of the dentate hilus (Raza et al., 2017). In general, hippocampal GABAergic interneurons also originate from ganglionic eminences like the cortical interneurons and follow the same migration streams (Khalaf-Nazzal and Francis, 2013; Lim et al., 2018). It will be interesting to investigate if these hippocampus-specific interneuron types have a unique developmental program.

The number of glutamatergic transcriptomic types has increased dramatically compared to our previous study (56 to 237 types) as the number of regions covered increased from 2 to ∼40, and we observed both region-specific types and types shared across multiple regions. The degree of distinction between glutamatergic cell types varies at multiple levels (**Fig. 1, 3**), and this may reflect the evolutionary distance between regions. The highest level is the distinction between isocortex and hippocampal formation, then between major regions within HPF, with largely non-overlapping subclasses at both levels. This may be due to the more ancient emergence of hippocampal formation and its subregions. For the more recently evolved isocortex, which has gone through accelerated expansion in both layers and areas, the dominant division of cells is by layers (or cortical depth). As such, the division of the 28 subclasses within the glutamatergic neuronal class is driven by a combination of layers and CTX/HPF regions. There is further differential regional specificity within the isocortex. The medial association areas, retrosplenial and anterior cingulate areas, have several unique cell types that are often shared between the two (e.g. the 131_L2 IT RSPv, 186_L2 IT RSP-ACA and 255_L5 PT RSP-ACA clusters, and the L3 RSP-ACA subclass). Likewise, the lateral association areas, temporal, perirhinal and ectorhinal (TPE) areas, share a unique cell type (184_L2/3 IT TPE), and share other types with the neighboring entorhinal cortex (i.e., the 201/202_L4/5 IT TPE-ENT cluster and the L5 IT TPE-ENT subclass). The rest of the CTX cell types are largely shared among multiple isocortical areas, with graded variation in gene expression and/or cluster distribution on the cortical sheet. These results suggest that medial and lateral association areas are more distinct from the other isocortical areas, again perhaps related to their evolutionary and developmental history.

Since all the regions within CTX and HPF have been profiled without gaps, multiple gradients (or continuous variations) of glutamatergic gene expression and cell type distribution have emerged from our dataset as a prominent feature of the molecular and cellular organization of these brain structures. In the isocortex, the major gradients are along the anterior-posterior or medial-lateral axis across cortical areas. These gradients have been observed in the L2/3, L4/5 and L6 IT subclasses, as well as the L5 PT and L6 CT subclasses (**Fig. 4, 5, 7, S13**). The complete characterization of the cortical sheet also reveals that when we only profiled VISp and ALM previously (Tasic et al., 2018), we only sampled the ends of the anterior-posterior gene expression gradient, which resulted in the separation of glutamatergic cells from these two areas as discrete clusters. With full profiling of the spatial axis, we now find that these cells belong to the same type that has continuous gene expression variation across regions. In addition, there is a continuous transitioning among L2/3, L4/5, L5 and L6 IT subclasses in the superficial-deep direction along the cortical depth (**Fig. 4**). This continuous transitioning is consistent, with slight variations, across multiple isocortical areas.

In the hippocampal formation, the glutamatergic subclasses of the hippocampal and subicular regions are joined in a major branch that is highly distinct from other glutamatergic subclasses. Within this major branch, we observed coordinated gradients of gene expression and cell type distribution from different regions that correspond to multiple spatial axes including dorsal-ventral, proximal-distal and superficial-deep (**Fig. 8**). Many previous studies had shown the existence of continuous or discrete subdivisions along these axes (Bienkowski et al., 2018; Cembrowski and Spruston, 2019). However, since these variations in multiple dimensions are intermingled in the convoluted hippocampal and subicular structures, in the absence of comprehensive and systematic transcriptomic data it was impossible to tease out the exact pattern of these variations. In this study, we have computationally extracted large sets of genes associated with the principal components (PCs) of the variations across cell types. Using the *in situ* expression patterns of these genes, we were able to recognize the correspondence between these PCs and spatial dimensions or anatomical locations. This has allowed us to derive a more complete picture of the multidimensional variation of gene expression and cell type distribution in the hippocampal and subicular regions (**Fig. 8, S16, S17**). Specifically, we find that glutamatergic cell types exhibit gradient distribution in the dorsal-ventral axis in all hippocampal-subicular regions examined (SUB, ProS, CA1, CA3 and DG); CA1, ProS and SUB cell types together form a proximal-distal transition zone; CA1 and CA3 cell types are also distributed along a superficial-deep division.

To compare the two major brain structures, the hippocampal formation (archicortex) and isocortex (neocortex), at cell type level, we computationally explored the relatedness between cell types and find that most HPF cell types match a specific CTX subclass (**Fig. 3D-E**). Strikingly, the resulting similarity based on transcriptomic profiles correlates well with the layer location (identified through marker genes) of HPF cell types, suggesting that related cell type pairs are also located in corresponding layers. We identified one-to-one correspondences of layer-specific cell types in all retrohippocampal regions, including ENT, PPP, SUB and ProS, with isocortical ones (it appears that deep L3 of ENT and PPP corresponds to L4/5 in CTX) (**Fig. 3, 6, 7**). Moreover, it is remarkable to see that the superficial and deep layer cell types in CA1 and CA3 correspond to CTX L2/3 IT and L5 PT types, respectively, and that these regions have yet a third type of cells resembling CTX L6 IT types (**Fig. 3, 8**). These similarities likely reflect the homologous relationships rooted in the evolutionary history of these regions, and raise the intriguing possibility that the axon projection patterns of the HPF glutamatergic neuron types may follow similar rules for those of isocortical neurons (e.g., corticocortical, corticofugal, corticothalamic, or local/near projections for IT, PT, CT or NP subclasses, respectively). Indeed, such rules have been observed in several cases. For example, ENTm and ENTl L2 and L3 neurons mainly project into the hippocampus and within ENT, whereas ENT L5 neurons mainly mediate the output projections (Nilssen et al., 2019), Similarly for SUB, it has been shown that superficial neurons mainly project within HPF whereas deep-layer neurons have subiculo-fugal or subiculo-thalamic projections (Bienkowski et al., 2018).

The glutamatergic neurons in hippocampal formation across the dorsal-ventral axis have shown differential cellular properties and connectivity patterns, and have been associated with different behavioral roles (Cembrowski and Spruston, 2019). A prominent function of the hippocampal formation is spatial navigation, with a number of functionally specific cell types identified in various HPF regions, such as grid cells in ENTm, head direction cells in PPP and ENTm, and place cells in CA1 (Moser et al., 2017). Underlying the functions, the input and output connections across many regions both within and outside HPF are highly complex yet highly organized (Bienkowski et al., 2018; van Strien et al., 2009). It will be of immense interest to examine the extent to which transcriptomic cell types are the nodes underlying specific connectional pathways and playing specific functional roles. For example, we hypothesize that the *Calb1*+ L3 ENTm cells and the *Reln*+ L2 ENTm cells may contain the pyramidal and stellate grid cells, respectively, based on the expression of these known grid cell marker genes (Ferrante et al., 2017; Nilssen et al., 2019). In addition, it has been shown that dorsal and ventral parts of the hippocampal and subicular regions preferentially form their respective interconnected networks (Bienkowski et al., 2018). The shared dorsal-ventral differentially expressed gene sets across these regions identified in our study suggest a common regulatory program for the development of these specific circuits.

As a last note, we acknowledge that our analysis of this large transcriptomic dataset is far from exhaustive. We have focused our analysis on the general organization of the transcriptomic landscape and its correspondence with the spatial organization of CTX and HPF through a variety of computational analyses and extensive anatomic annotations. Within the scope of this single paper, we have not fully investigated variations among individual clusters nor any correlations with projection targets or other cell type properties. Here we have assigned anatomical locations of major cell types (mostly at the supertype level) using existing RNA ISH data from the Allen Brain Atlas, and provided an estimate of the relative proportions of different cell types in each region using the 10xv2 data. However, the precise spatial distribution and relative proportion of various cell types should ultimately be established through future spatial transcriptomic studies using approaches such as multiplexed FISH or *in situ* sequencing (Eng et al., 2019; Lein et al., 2017; Moffitt et al., 2018; Rodriques et al., 2019; Wang et al., 2018).

Despite these limitations, our current study establishes the first blueprint of the molecular architecture that potentially reflects the developmental/evolutionary origins as well as the connectional/functional specificity of the isocortex and hippocampal formation and their subregions. This work also provides the roadmap to genetically target the numerous cell types newly discovered and categorized here, and lays the foundation for systematic, cell type-specific investigation of the structure and function of these brain circuits.

## ACKNOWLEDGMENTS

We are grateful to the Transgenic Colony Management, Neurosurgery & Behavior, Lab Animal Services, Molecular Biology and Histology teams at the Allen Institute for technical support. We thank Christof Koch, Ed Lein, Michael Hawrylycz and Allan Jones for their support and leadership. The research was funded by multiple grant awards from institutes under the National Institutes of Health (NIH), including award number R01EY023173 from The National Eye Institute to H.Z., U01MH105982 from the National Institute of Mental Health and Eunice Kennedy Shriver National Institute of Child Health & Human Development to H.Z., and U19MH114830 from the National Institute of Mental Health to H.Z. The content is solely the responsibility of the authors and does not necessarily represent the official views of NIH and its subsidiary institutes. This work was also supported by the Allen Institute for Brain Science. The authors thank the Allen Institute founder, Paul G. Allen (1953-2018), for his vision, encouragement, and support.

## AUTHOR CONTRIBUTIONS

Contribution to RNA-seq data generation: led by H.Z., K.A.S., B.T., N.D., with contributions from T.N.N., J.Goldy, D.B., T.C., K.C., O.F., A.G., J.Gray, D.H., M.K., K.L., B.L., D.M., T.P., C.R., N.S., J.S., M.T., A.T., H.T., K.W. Contribution to data analysis: led by Z.Y., with contributions from T.N.N., C.T.J.vV., J.Goldy, A.E.S-C., F.B., S-L.D., O.F., E.G., L.T.G., K.A.S., B.T., H.Z. Contribution to data interpretation: Z.Y., T.N.N., C.T.J.vV., S-L.D., Q.R., B.T., H.Z. Contribution to writing manuscript: led by H.Z., with contributions from Z.Y., T.N.N., C.T.J.vV., B.T., and inputs from other co-authors. Project/program management: S.M., S.M.S. Overall project supervision: H.Z., B.T., K.A.S., Z.Y.

## Methods

### Mouse breeding and husbandry

All procedures were carried out in accordance with Institutional Animal Care and Use Committee protocols at the Allen Institute for Brain Science. Animals were provided food and water *ad libitum* and were maintained on a regular 12-h day/night cycle at no more than five adult animals per cage. Animals were maintained on the C57BL/6J background, and newly received or generated transgenic lines were backcrossed to C57BL/6J. Standard tamoxifen treatment for CreER lines included a single dose of tamoxifen (40 μl of 50 mg ml^−1^) dissolved in corn oil and administered via oral gavage at postnatal day (P)10–14. Tamoxifen treatment for *Nkx2.1-CreERT2;Ai14* was performed at embryonic day (E)17 (oral gavage of the dam at 1 mg per 10 g of body weight), pups were delivered by cesarean section at E19 and then fostered.

*Cux2-CreERT2;Ai14* mice received tamoxifen treatment at P35 ± 5 for five consecutive days. Trimethoprim was administered to animals containing *Ctgf-2A-dgCre* by oral gavage at P40 ± 5 for three consecutive days (0.015 ml per g of body weight using 20 mg ml^−1^ trimethoprim solution). *Ndnf-IRES2-dgCre* animals did not receive trimethoprim induction, since the baseline dgCre activity (without trimethoprim) was sufficient to label the cells with the *Ai14* reporter (Tasic et al., 2018). We excluded any animals with anophthalmia or microphthalmia. We used 538 animals to collect 74,973 cells for SSv4 and 48 animals to collect 1,093,036 cells for 10xv2. Animals were euthanized at P53-59 (*n* = 530), P50-52 (*n* = 6), and P60-121 (*n* = 50). No statistical methods were used to predetermine sample size. All donors used in this study are listed in **Table S2**.

### Retrograde Labeling

We injected rAAV2-retro-EF1a-Cre (Tervo et al., 2016), RVΔGL-Cre (Chatterjee et al., 2018), or CAV-Cre (gift of Miguel Chillon Rodrigues, Universitat Autònoma de Barcelona) (Hnasko et al., 2006) into brains of heterozygous or homozygous Ai14 mice as previously described (Tasic et al., 2016, 2018). For ALM experiments, we also injected rAAV2-retro-CAG-GFP or rAAV2-retro-CAG-tdTomato (Tervo et al., 2016) into wild-type mice. We injected rAAV2-retro-EF1a-dTomato (Tervo et al., 2016) into *Gnb4-IRES2-CreERT2;Ai140* and *Cux2-CreERT2;Ai140* (Daigle et al., 2018) mice with the goal of collecting the Car3 cell types. We collected both singly positive (dTomato+) and double-positive (GFP+/dTomato+) cells when possible. We injected rAAV2-retro-EF1a-dTomato (Tervo et al., 2016) into *Ctgf-T2A-dgCre;Snap25-LSL-F2A-GFP* mice, with the goal of collecting L6b projection neurons. Stereotaxic coordinates were obtained from Paxinos adult mouse brain atlas (Franklin and Paxinos, 2008) (Supplementary Table 6 in (Tasic et al., 2018)). For two VISp experiments, we injected into SCs by inserting the needle through the cerebellum at a 45°-angle in the posterior to anterior direction. Injection information for each donor is available in **Table S2**.

### Retro-orbital Labeling

We delivered viruses that contain an enhancer element with putative specificity to L5 PT types (rAAV-mscRE4-minBGpromoter-FlpO-WPRE3) and L5 IT and L6 IT types (rAAV-mscRE10-minBGpromoter-FlpO-WPRE3, rAAV-mscRE16-minBGpromoter-FlpO-WPRE3) (Graybuck et al., https://www.biorxiv.org/content/10.1101/525014v2) into heterozygous or homozygous Ai65 mice into the retroorbital sinus as previously described (Chan et al., 2017). This approach allows the virus to cross the blood-brain barrier for brain-wide delivery of the viral particles. Due to the difficulty of isolating L5 PT neurons, we used this approach to enrich for labeling of specific types for more efficient cell isolation.

### Single-cell isolation

We isolated single cells by adapting previously described procedures (Hempel et al., 2007; Sugino et al., 2006; Tasic et al., 2016, 2018). The brain was dissected, submerged in ACSF (Tasic et al., 2018), embedded in 2% agarose, and sliced into 250-μm (SMART-Seq) or 350-μm (10x Genomics) coronal sections on a compresstome (Precisionary Instruments). Block-face images were captured during slicing. Regions of interest (ROIs) were then microdissected from the slices and dissociated into single cells with 1 mg/ml pronase (SMART-Seq before 28 June 2018, Sigma P6911-1G) and processed as previously described (Tasic et al., 2018). Fluorescent images of each slice before and after ROI dissection were taken from the dissecting scope. All these images were used to document the precise location of the ROIs using annotated coronal plates of CCFv3 as reference (see below).

We used Allen Mouse Brain Common Coordinate Framework version 3 (CCFv3) ontology (http://atlas.brain-map.org/) to define brain regions for profiling and boundaries for dissections. We covered all regions of CTX and HPF, and chose sampling at mid-ontology level with judicious joining of neighboring regions (**Table S1**). These choices were guided by the fact that microdissections are difficult for small regions and that joining was sometimes necessary to obtain sufficient numbers of cells for profiling, especially for 10xv2. For example, orbital area (ORB) covers ORBl, ORBm and ORBvl (see **Table S1** for full regional names), and primary somatosensory area (SSp) covers all SSp subfields. Sometimes neighboring small regions were combined to form a joint dissected region, for example, GU-VISC, PL-ILA and TEa-PERI-ECT were combinations of 2 or 3 cortical areas, respectively (**Table S1**). The PPP-SP joint region for 10xv2 includes para-, post-, and pre-subiculum, the subiculum proper and pro-subiculum (i.e. PAR-POST-PRE-SUB-ProS).

We used transgenic driver lines for fluorescence-positive cell isolation to enrich for neurons, with the vast majority being Cre driver lines crossed to the *Ai14*-tdTomato reporter (**Table S2**). A small fraction of SSv4 cells were labeled by retrograde tracing (Retro-seq) or retroorbital injection of AAVs (**Table S2**). All 10xv2 cells from all regions were isolated from the pan-neuronal *Snap25-IRES2-Cre* line (**Fig. S1H**). For SSv4, transgenic mice used, dissection scheme, and sampling rate varied by regions (**Fig. S1G, Table S2**). Our previously published VISp and ALM (part of MOs) SSv4 dataset (∼24,000 cells) (Tasic et al., 2018) were also included in the current study; this dataset had utilized a large number of driver lines with either broad or highly specific coverage of different cell types, and employed extensive layer-specific dissections.

Three other cortical regions, MOp, SSp and ACA, had 5,000-6,600 SSv4 cells each that were also from multiple driver lines and layer-specific dissections. The hippocampal region (HIP), which includes CA1, CA2, CA3 and DG, was divided into 4 anteroposterior (i.e. dorsoventral) segments, and each segment was profiled using both pan-glutamatergic and pan-GABAergic Cre lines, totaling ∼6,600 SSv4 cells. All the other CTX and HPF regions had 1,100-2,000 SSv4 cells each, profiled from pan-glutamatergic and pan-GABAergic Cre lines without layer-specific dissections. Finally, we also included ∼1,000 SSv4 cells from the claustrum (CLA), a region that does not belong to either CTX or HPF, because they co-cluster with some cortical cells into the Car3 subclass, and these cells came from various Cre lines or retrograde labeling (**Table S2**).

We used mice of both sexes, but didn’t achieved complete sex balance in sampling (**Fig. S1E-F**). The 10xv2 datasets had only male cells for MOp, PL-ILA-ORB, RSP, and only female cells for SSp. The SSv4 dataset had more equal sex coverage. There were only two small, sex specific clusters, #249 and #355, both male specific and almost all the cells in these two clusters were from PL-ILA-ORB and RSP 10xv2 data.

For all 10xv2 samples and for SSv4 samples after 28 June 2018, we improved our protocol with the following changes. Tissue pieces were digested with 30 U/ml papain (Worthington PAP2) in ACSF for 30 minutes at 30°C. Due to the short incubation period in a dry oven, we set the oven temperature to 35°C to compensate for the indirect heat exchange, with a target solution temperature of 30°C. Enzymatic digestion was quenched by exchanging the papain solution three times with quenching buffer (ACSF with 1% FBS and 0.2% BSA). Samples were incubated on ice for 5 minutes before trituration. The tissue pieces in the quenching buffer were triturated through a fire-polished pipette with 600-µm diameter opening, approximately 20 times. The tissue pieces were allowed to settle and the supernatant, which now contains suspended single cells, was transferred to a new tube. Fresh quenching buffer was added to the settled tissue pieces, and trituration and supernatant transfer were repeated using 300-µm and 150-µm fire polished pipettes. The single cell suspension was passed through a 70-µm filter into a 15-ml conical tube with 500 ul of high BSA buffer (ACSF with 1% FBS and 1% BSA) at the bottom to help cushion the cells during centrifugation at 100xg in a swinging bucket centrifuge for 10 minutes. The supernatant was discarded, and the cell pellet was resuspended in the quenching buffer.

All cells were collected by fluorescence-activated cell sorting (FACS, BD Aria II) using a 130-μm nozzle. Cells were prepared for sorting by passing the suspension through a 70-µm filter and adding DAPI (to the final concentration of 2 ng/ml). Sorting strategy was as previously described (Tasic et al., 2018), with most cells collected using the tdTomato-positive label. For SSv4, single cells were sorted into individual wells of 8-well PCR strips containing lysis buffer from the SMART-Seq v4 kit with RNase inhibitor (0.17 U/μl), immediately frozen on dry ice, and stored at −80°C. For 10x Genomics, 30,000 cells were sorted within 10 minutes into a tube containing 500 µl of quenching buffer. Sorting more cells into one tube dilutes the ACSF in the collection buffer, causing cell death. We also observed decreased cell viability for longer sorts. Each aliquot of sorted 30,000 cells was gently layered on top of 200 µl of high BSA buffer and immediately centrifuged at 230xg for 10 minutes in a centrifuge with a swinging bucket rotor.

The high BSA buffer at the bottom of the tube slows down the cells as they reach the bottom in order to minimize cell death. No pellet could be seen with this small number of cells, so we took out the supernatant and left behind 35 µl of buffer, in which we resuspended the cells. The immediate centrifugation and resuspension allowed the cells to be temporarily stored in a high BSA buffer with minimal ACSF dilution. The resuspended cells were stored at 4°C until all samples were collected, usually within 30 minutes. Samples from the same ROI were pooled, cell concentration quantified, and immediately loaded onto the 10x Genomics Chromium controller.

### cDNA amplification and library construction

For SSv4 processing, we performed the procedures with positive and negative controls as previously described (Tasic et al., 2018). We used the SMART-Seq v4 Ultra Low Input RNA Kit for Sequencing (Takara Cat# 634894) to reverse transcribe poly(A) RNA and amplify full-length cDNA. We performed reverse transcription and cDNA amplification for 18 PCR cycles (neurons) or 21 PCR cycles (non-neuronal cells) in 8-well strips, in sets of 12–24 strips at a time. All samples proceeded through Nextera XT DNA Library Preparation (Illumina Cat# FC-131-1096) using Nextera XT Index Kit V2 (Illumina Cat# FC-131-2001) and custom index sets (Integrated DNA Technologies). Custom index sets were validated to confirm the same performance as Nextera Index sets before being used on experimental samples. Nextera XT DNA Library prep was performed according to manufacturer’s instructions, with a modification to reduce the volumes of all reagents and cDNA input to 0.4× or 0.5× of the original protocol. Details are available in ‘Documentation’ on the Allen Institute data portal at: http://celltypes.brain-map.org.

For 10xv2 processing, we used Chromium Single Cell 3’ Reagent Kit v2 (10x Genomics Cat# 120237). We followed manufacturer’s instructions for cell capture, barcoding, reverse transcription, cDNA amplification, and library construction. We targeted sequencing depth of 60,000 reads per cell; the actual median achieved is 57,192 reads per cell across 162 libraries.

### Sequencing data processing and QC

Processing of SSv4 libraries was performed as described previously (Tasic et al., 2018). Briefly, libraries were sequenced on an Illumina HiSeq2500 platform (paired-end with read lengths of 50 bp) to a target read depth of 0.5M reads per cell (range 100,275-12,329,698, median 1,003,867). The Illumina sequencing reads were aligned to GRCm38.p3 (mm10) using a RefSeq annotation gff file retrieved from NCBI on 18 January 2016 (https://www.ncbi.nlm.nih.gov/genome/annotation_euk/all/). Sequence alignment was performed using STAR v2.5.3 (Dobin et al., 2013) in the two-pass mode. PCR duplicates were masked and removed using STAR option ‘bamRemoveDuplicates’. Only uniquely aligned reads were used for gene quantification. Gene counts were computed using the R GenomicAlignments package (Lawrence et al., 2013) and summarizeOverlaps function in ‘IntersectionNotEmpty’ mode for exonic and intronic regions separately. For the SSv4 dataset, we only used exonic regions for gene quantification. Cells that met any one of the following criteria were removed: < 100,000 total reads, < 1,000 detected genes (with CPM > 0), < 75% of reads aligned to genome, or CG dinucleotide odds ratio > 0.5. Doublets were removed by first classifying cells into broad classes of excitatory, inhibitory, and non-neuronal based on known markers. Reads that did not map to the genome were then aligned to synthetic constructs (i.e. ERCC) sequences and the *E. coli* genome (version ASM584v2) and were used as a QC metric.

10xv2 libraries were sequenced on Illumina NovaSeq6000 and sequencing reads were aligned to the mouse pre-mRNA reference transcriptome (mm10) using the 10x Genomics CellRanger pipeline (version 3.0.0) with default parameters. Cells that met the following criterion were filtered out for downstream processing in each 10x run: < 1,500 detected genes (with UMI count > 0). Doublets were identified using a modified version of the DoubletFinder algorithm (McGinnis et al., 2019) and removed when doublet score > 0.3. Doublets were further removed by first classifying cells into broad cell classes (neuronal versus non-neuronal) based on the co-expression of any pair of broad class marker genes.

### Clustering

Clustering for both SSv4 and 10xv2 datasets was performed independently using the in-house developed R package **scrattch.hicat** (available via github https://github.com/AllenInstitute/scrattch.hicat). In addition to classical single-cell clustering processing steps provided by other tools such as Seurat (Butler et al., 2018), this package features automatic iterative clustering while ensuring all pairs of clusters, even at the finest level, are separable by fairly stringent differential gene expression criteria (Tasic et al., 2018). For the 10xv2 dataset, we used q1.th = 0.4, q.diff.th = 0.7, de.score.th = 150, min.cells = 20; for the SSv4 dataset, we used q1.th = 0.5, q.diff.th = 0.7, de.score.th = 150, min.cells = 4. The package also performs consensus clustering by repeating the iterative clustering step on 80% subsampled set of cells 100 times, and then derives the final clustering result based on cell-cell co-clustering probability matrix. This feature enables us to both finetune clustering boundaries and to assess clustering uncertainty.

For the 10xv2 dataset, due to the large data size, we adapted the existing scrattch.hicat package to **scrattch.bigcat** package, which uses bigstatsr package as backend. Bigstatsr allows for manipulation of matrices that are too large to fit in memory through memory mapping to files on disk. This enables storage of the gene count matrix from the complete > 1M cells, while facilitating efficient random access of cells. During each iteration of clustering, the algorithm randomly samples up to 5,000 cells and loads them to the memory to perform high-variance gene selection and PCA, and then computes the reduced dimensions for the whole dataset by applying the same projection on all the cells. The reduced dimensions are used to compute the K nearest neighbors (KNN) using the RANN package (https://github.com/jefferislab/RANN), that are then used to perform Jaccard-Louvain clustering.

### Joint clustering between the 10xv2 and SSv4 datasets

To provide one consensus cell type taxonomy based on both 10xv2 and SSv4 datasets, we developed a novel integrative clustering analysis across multiple data modalities, now available via the **i_harmonize** function of the **scrattch.hicat** package. This method extends the clustering pipeline described above to incorporate datasets collected by different transcriptomic platforms. Unlike the Seurat CCA approach (Butler et al., 2018) and scVI (Svensson et al., 2020) which aim to find aligned common reduced dimensions across multiple datasets, this method directly builds a common adjacency graph using all cells from all datasets, then applies the standard Jaccard-Louvain clustering algorithm. We extended the cluster merging algorithm described above to ensure that all clusters can be separated by conserved DE genes across platforms. The **i_harmonize** function, similar to the **iter_clust** function in the single dataset clustering pipeline, applies the integrative clustering across datasets iteratively, while ensuring all clusters at each iteration are separable by conserved DE genes. This is an important feature of this method, as we aim to build a fine resolution taxonomy of increasing complexity, no clustering algorithm can provide proper resolution of cell types in one round.

To build the common graph that incorporates the samples from all the datasets, we first chose a subset of reference datasets from all available datasets, which either provides more sensitive gene detection and/or more comprehensive cell type coverage. For this study, as the 10xv2 dataset includes more cells while the SSv4 dataset provides more sensitive gene detection, both datasets were used as reference datasets.

The key steps of the pipeline are outlined below:

1. **Select anchor cells for each reference dataset.** For each reference dataset, we randomly sample 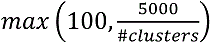
 anchor cells per cluster to achieve more uniform coverage of cell type. This is the only place during the joint clustering step that uses the platform specific clustering information.
2. **Select high variance genes.** Selecting high variance genes and PCA dimension reduction are performed using scrattch.hicat package. We defined several vectors that correspond to potential technical bias: the number genes detected in each cell, mitochondrial gene expression, and donor specific gene expression. Particularly, we identified a set of highly specific genes with significantly elevated gene expression in two out of three male donors used in TPE 10xv2 experiments. The top PC based on this gene set is used to help us track donor specific bias. PCs with more than 0.6 Pearson correlation with any of the technical bias vectors defined above were removed. For each remaining PCs, Z scores are calculated for gene loadings, and top 100 genes with absolute Z score greater than 2 are selected. The high variance genes from each reference datasets are pooled.
3. **Compute K nearest neighbors (KNN).** For each cell in each dataset, we compute its K nearest neighbors among anchor cells in each reference datasets based on the high variance genes selected above. The RANN package was used to compute KNN based on the Euclidean distance when the query and reference dataset is the same. To compute nearest neighbors across datasets, we used correlation as a similarity metric.
4. **Compute the Jaccard similarity**. For every pair of cells from all datasets, we compute their Jaccard similarity, defined as the ratio of the number of shared K nearest neighbors (among all anchors cells) over the number of combined K nearest neighbors.
5. Perform Louvain clustering based on Jaccard similarity.
6. **Merge clusters.** To ensure that every pair of clusters are separable by conserved differentially expressed (DE) genes across all datasets, for each cluster, we first identified the top 3 nearest clusters. For each pair of such close-related clusters, we computed the DE genes in each dataset (Tasic et al., 2018), and chose the DE genes that were significant in at least one dataset, while also having more than two-fold change in the same direction in both datasets. We then compute the overall statistical significance based on such conserved DE genes for each dataset independently. If any of the datasets pass our DE gene criteria described in the “clustering” section, the pair of clusters remain separated, otherwise they are merged. DE genes are recomputed for the merged clusters, and the process repeats until all clusters are separable by the conserved DE genes criteria. If one cluster has fewer than the minimal number of cells in a dataset (4 cells for SSv4 and 20 cells for 10xv2), then this dataset is not used for DE gene computation for all pairs involving the given cluster. This step allows detection of unique clusters only present in some data types.
7. Repeat steps 1-6 for cells within a cluster to gain finer-resolution clusters until no clusters can be found.
8. Concatenate all the clusters from all the iterative clustering steps, and perform final merging as described in step **6.**

This integrative clustering pipeline allows us to resolve clusters at fine resolution, while ensuring proper alignment between datasets by requiring presence of conserved DE genes. It also allows us to leverage the strengths of different datasets. For example, between clusters that are separated by weakly expressed genes, SSv4 dataset provides the statistical power for separation, and the relevant genes help to separate 10xv2 cells into clusters with consistent fold changes. On the other hand, for clusters that have very few cells in SSv4, the 10xv2 samples provide the statistical power for separation, and relevant genes are used to split SSv4 cells accordingly, and we can identify rare clusters that are present predominantly only in one platform.

### Excluding noisy clusters

We identified 408 clusters by the integrative clustering pipeline. There were three main categories of noisy clusters: clusters located outside areas of interest due to inaccurate dissection, clusters with significantly lower gene detection due to extensive drop out, and clusters due to doublets or contamination.

We identified 6 clusters that were outside of CTX and HPF, and located them to the striatum based on the expression of marker genes such as *Six3*, *Adora2a* and *Drd2*.

The remaining clusters were grouped at subclass level based on the taxonomy tree and correspondence with our previous cortical taxonomy (Tasic et al., 2018). While we observed differences in the number of genes detected at the subclass level, cells within a subclass had relatively homogeneous distribution. For every subclass, we assigned those clusters with average number of detected genes below a certain threshold (defined by mean and sd of the subclass) as putative low-quality clusters. If we also identified another cluster with more cells, at least 500 more detected genes, and no more than one down-regulated gene relative to the putative low-quality cluster, then we consider the putative low-quality cluster confirmed.

To identify doublet clusters, we searched for triplets of clusters A, B and C, wherein A is the putative doublet cluster, such that up-regulated genes of A relative to B largely overlap with up-regulated genes in C relative to B, and up-regulated genes in A relative to C largely overlap with up-regulated genes of B relative to C. This criterion ensures that A includes the most distinguished signature of B and C. To rule out the possibility that A is a transitional type between B and C, an additional requirement was that B and C cannot be closely related types, based on the correlation of their average gene expression of marker genes. After we systematically produced the list of all the candidate triplet clusters, the final determination was an iterative process that involved setting different thresholds and manual inspection.

### Marker gene selection

For each pair of clusters, we computed the conserved DE genes (at least significant in one dataset, and at least 2-fold change in the same direction in the other datasets). We selected the top 50 genes in each direction, and pooled such genes from all pairwise comparisons with a total of 4,963 genes as markers.

### Assessing concordance of joint clustering between 10xv2 and SSv4

We first compared the joint clustering results with the independent clustering result from each dataset. We then calculated the cluster means of marker genes for each dataset. For each marker gene, we computed the correlation between its average expression for each cluster across two different datasets to quantify the consistency of its expression at the cluster level between datasets.

### Imputation

To facilitate direct comparison, we projected gene expression of SSv4 dataset to the 10xv2 reference data and vice versa. To do that, we leveraged the KNN matrices computed during the iterative joint clustering step. During each iteration of the joint clustering, we used the average gene expression of the k nearest neighbors among the 10xv2 anchor cells as the imputed expression for each SSv4 cell. At the top-level clustering, we imputed the expression for all the genes. For each following iteration, we only imputed the expression of the high variance genes or the DE genes computed for the cells involved in the given iteration. We used this iterative approach for imputation because the nearest neighbors based on the genes chosen at the top level may not reflect the distinction between the finer types, and the imputed values for the DE genes that define the finer types consequently are not accurate based on these nearest neighbors.

Therefore, we deferred imputation of the DE genes between the finer types to the iteration when these types were defined. This method is now provided in the **impute_knn_global** function in the scrattch.hicat package. We computed the imputed gene expression matrix using either SSv4 or 10xv2 data as reference. Unless specified, we used 10xv2 imputed gene expression by default.

### Building cell type taxonomy tree

We computed the average expression of marker genes at the cluster level based on imputed gene expression using either SSv4 or 10xv2 data as reference. The two matrices were concatenated, and the tree was constructed using the **build_dend** function in the scrattch.hicat package as described in (Tasic et al., 2018).

### UMAP projection

We performed principal component analysis (PCA) on both 10xv2 and SSv4 based on imputed gene expression matrices of 4,963 marker genes, and selected the top 60 PCs for both. We removed the PCs with more than 0.6 correlation with any technical bias vectors. We used PCs from 10xv2 based imputed data as input to create 2D and 3D UMAP, using parameters nn.neighbors = 25 and md = 0.4.

### Assigning cluster names

We assigned cluster IDs based on the order of clusters in the taxonomy tree. Based on the topology of the taxonomy tree, we defined classes and subclasses following the convention from (Tasic et al., 2018). Based on the Allen Institute proposal for cell type nomenclature (https://portal.brain-map.org/explore/classes/nomenclature), we also assigned accession numbers to cell types (**Table S3**).

### Constellation plot

The global relatedness between cell types is visualized using constellation plots. These summarize the identity and relationship between clusters and were generated as follows. In the constellation plot, each transcriptomic cluster is represented by a node whose surface area reflects the number of cells within the cluster in log scale. The orientation of the nodes is based on the centroid positions of the clusters in UMAP coordinates. The relationship between nodes is indicated by edges between nodes and they were calculated as follows. For each cell its 15 nearest neighbors in reduced dimension space were determined and summarized by cluster. For each cluster, we then calculated the fraction of nearest neighbors that were assigned to other clusters. The edges connect two nodes in which at least one of the nodes has >5% of nearest neighbors in the connecting node. The width of the edge at the node reflects the fraction of nearest neighbors that are assigned to the connecting node and is scaled to the node size. We then determine for all nodes in the plot what the maximum fraction of “outside” neighbors is and set this as edge width = 100% of node width.

### Correspondence between CTX and HPF clusters

Glutamatergic cell types are highly distinct between CTX and HPF regions, but they also have intricate relationships according to the taxonomy tree, UMAP projections and constellation plots. To study their correspondence systematically, we first computed the top 50 DE genes in each direction for all pairs of glutamatergic clusters in CTX (2,469 genes in total) and HPF (2,389 genes) separately. Transitional clusters in L5 IT TPE-ENT and L2/3 IT PPP were excluded from this analysis. Interestingly, 1,781 or about 75% of genes were in common. Using this common set of DE genes that discriminate cell types in both structures, we mapped each cell in HPF clusters to the most correlated CTX cluster. Then we computed the frequency of the cells in each HPF cluster mapped to each CTX cluster or subclass (subclass level mapping is shown in Fig. 3D). Between each HPF cluster and its most correlated CTX cluster, we also computed the number of DE genes in each direction, shown in Fig. 3E.

### Gradient analysis for glutamatergic neurons in CTX and HPF regions

We used principal component analysis (PCA) and UMAP projection to extract major gradients that drive cell type diversities. To extract the most dominant gradient for isocortical IT types, we computed one dimensional UMAP for all the IT cells based on the PCs in the imputed space (see section **UMAP projection)**, which corresponded very well with cortical depth. UMAP instead of PCA was chosen here as we observed a nonlinear relationship. Particularly, in the PCA space, L6 IT types were more similar to L2/3 IT types than L5 IT types to L2/3 IT types, supported by many shared markers between L2/3 IT and L6 IT types. On the other hand, the transitions between adjacent layers were also obvious. One dimensional UMAP preserved such nonlinear transitions faithfully. Then for cells within each subclass, which had relatively homogenous distribution along the cortical depth, we categorized regional diversity using PCA based on regional specific genes, defined as the top 20 DE genes in each direction between all pairs of dissected regions in the imputed gene space. To determine a refined list of genes that have strong contribution to regional gradients, we chose up to 30 top genes, whose expressions were most correlated/anticorrelated with each PC, with absolute Pearson correlation greater than 0.3. We then recomputed PCA using this refined gene set.

One dimensional UMAP was also used to extract the Pr-Di gradient between CA1 and SUB, for which each stage of the transition was driven by a different set of genes. We then computed PCAs within CA1 and CA3 based on the DE genes between the clusters within each region. In both cases, the first PC corresponded to the Do-Ve axis, and the second PC largely corresponded to the Su-De axis, validated by ISH images of the key genes that drive each axis. We computed the correlation of the CA1 and CA3 marker genes with the top PC in CA1 and CA3, respectively, and selected a core set of genes with absolute correlation greater than 0.4 in both cases. These include 96 dorsal-specific genes and 216 ventral-specific genes that are shared by CA1 and CA3. This gene set was used to compute the top PC within each subclass, and scaled in range of [0,1] as shown in **Fig. S16C**, which corresponds to the Ve-Do axis not only in CA1 and CA3, but also in SUB, ProS and DG. We further noticed that many genes in this set still show variation along Pr-Di axis, particularly, ProS and CA1-ProS clusters marked by *Dcn* and *Dlk1* overall have stronger expression of ventral specific genes, pushing these clusters close to the ventral end of the axis. To recalibrate Do-Ve axis to be more faithful to the actual spatial Do-Ve location, we binned the clusters along Pr-Di axis shown in **Fig.S16A**: #263-265 for SUB, #266-272 for ProS, #273-277 for CA1-ProS, and #278-290 for CA1, and separately for CA3 and DG. Then within each bin, we used the core gene set to recompute the top PC, and scaled the values in the range of [0,1]. These rescaled values were used to compute the cluster coordinates along the Y-axis in Fig. 8J. For CA3, we didn’t identify the proximal-distal axis as a major PC. However, close examination revealed a previously known CA3 proximal gene, *Fmo1* (Thompson et al., 2008), that is expressed in dorsal clusters #298 and #300, but not #299. We computed the DE genes between clusters #298 and #299, most of which also separate clusters #298, 300 from #299 and other ventral CA3 clusters. The top PC based on this set of genes is defined as the CA3 proximal-distal axis, which also correlates heavily with the dorsal-ventral axis. This is consistent with the previously defined CA3 subdomains (Thompson et al., 2008), which divided CA3 into a series of diagonal bands oriented septal-distally (toward CA2) to temporal-proximally (toward DG). To define the Su-De axis in CA1, we simply used the second PC. For CA3, genes specific to the most ventral cluster #293, including *Tcerg1l*, *Bace2*, *Doc2b*, *Rnd2* and *Bves*, also contributes to the Su end of PC2. Dorsal clusters #298-300 do not show much layer separation. To recalibrate Su-De axis, we used the top PC based on the DE genes between clusters #294, 297 at the deep end, and more superficial clusters #292, 295, and 296, excluding the most ventral cluster #293, dorsal clusters #298-300, and activated cluster #294.

Again, all values were scaled in range of [0,1]. Additionally, we observed several pairs of adjacent clusters differ mostly by IEGs such as *Fos* and *Arc*. We curated the top DE genes from such pairs of clusters (**Fig. S16E, F**), and defined the top PC based on this gene set as the activity axis.

### Assessing correspondence to external datasets

The median gene expression for each cell type in 10v2 and SSv4 datasets was computed separately using the cell type marker genes defined in the “**Marker gene selection”** section. For each external dataset, we used the genes that intersect with our marker list. If the RNA-seq method of the external dataset was Smart-seq, then we used our SSv4 cells as the reference for mapping, and if it was 10x, we used our 10xv2 cells as reference. Correlation-based mapping was performed to find the cell type for each individual cell from each external dataset. Mapping for each cell was performed 100 times. In each iteration, 80% of the genes were selected randomly and the correlation of gene expression of that cell with each cluster median in our dataset was computed and the cluster with the highest correlation was chosen as the cluster for that cell in that round. After 100 iterations, the percentage of time a cell was mapped to a given cell type in our reference dataset was defined as the probability of mapping to that cluster for that cell. Finally the cell type with the highest probability of mapping was chosen as the corresponding cell type of that cell.

### Data analysis software and visualization tools

Analysis and visualization of transcriptomic data were performed using R v3.5.0 and greater {R Core Team}, assisted by the Rstudio IDE {RStudio Team} and the scrattch.hicat, scrattch.bigcat and scarttch.vis packages {REFs/links}

### Data and code availability

The raw sequencing data is deposited in the NeMO Archive (https://biccn.org/data). Full metadata for all samples are available in **Table S2, S3**. R packages for the iterative clustering method utilized in this analysis (scrattch.bigcat and scrattch.hicat) are available on GitHub at https://github.com/AllenInstitute/scrattch.hicat. The transcriptomic data can be visualized and analyzed using the Transcriptomics Explorer available on https://portal.brain-map.org/.

We also provide an accompanying website at https://taxonomy.shinyapps.io/ctx_hip_browser/, with a Cell Card for each cell type. The website can be browsed by cell type and provides information on specific markers, cell type metadata and relation to neighboring cell types.

## Supplemental Information

**Figure S1.**
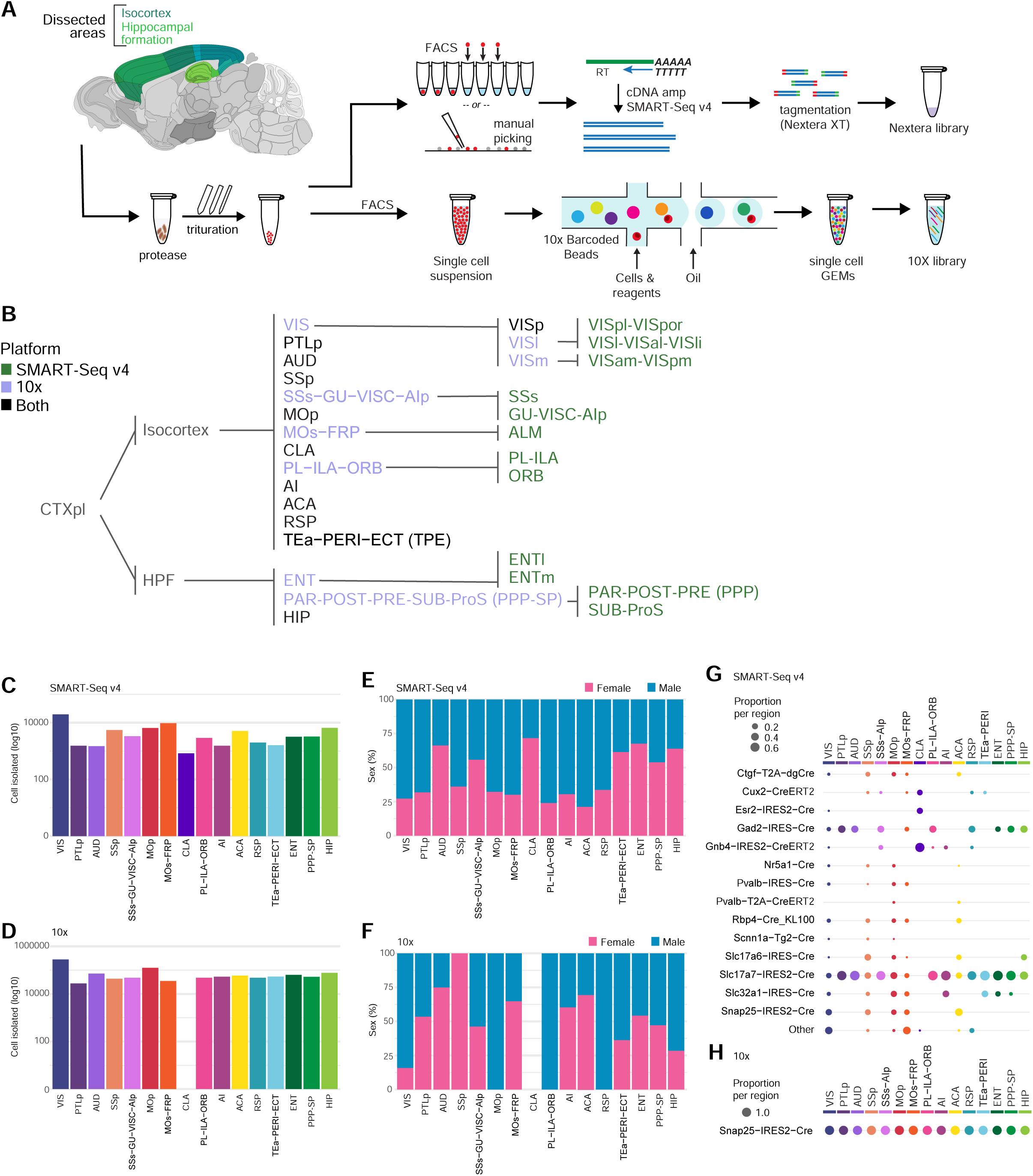
Experimental workflow and sample collection. **(A)** Schematic diagram illustrating cell isolation process for single cell RNA-sequencing. Brain regions were dissected according to the Allen CCFv3. Each sample was digested and triturated to obtain single cell suspensions. For SMART-Seq v4 (SSv4) processing, individual cells were sorted into 8-well strip PCR tubes by FACS or by manual picking. Cells were lysed, and SSv4 was used to reverse-transcribe and amplify full-length cDNAs from each cell. cDNAs were then tagmented by Nextera XT, PCR-amplified, and processed for Illumina sequencing. For 10x processing, debris was removed from single cell suspension by FACS, suspensions were loaded on the 10x Genomics Chromium™ Controller to create single-cell libraries, and libraries were processed for Illumina sequencing. **(B)** Ontology of dissected brain regions according to CCFv3. See Table S1 for each region’s full name and sampling. **(C-D)** The number of cells sampled from each dissection region for SSv4 (**C**) or 10xv2 (**D**). Bars are colored by region. **(E-F)** Sex sampling proportion per joint region for SSv4 (**E**) or 10xv2 (**F**). **(G-H)** Genotype sampling proportion per joint region for SSv4 (**G**) or 10xv2 (**H**), colored by region. Columns add up to 100%.

**Figure S2.**
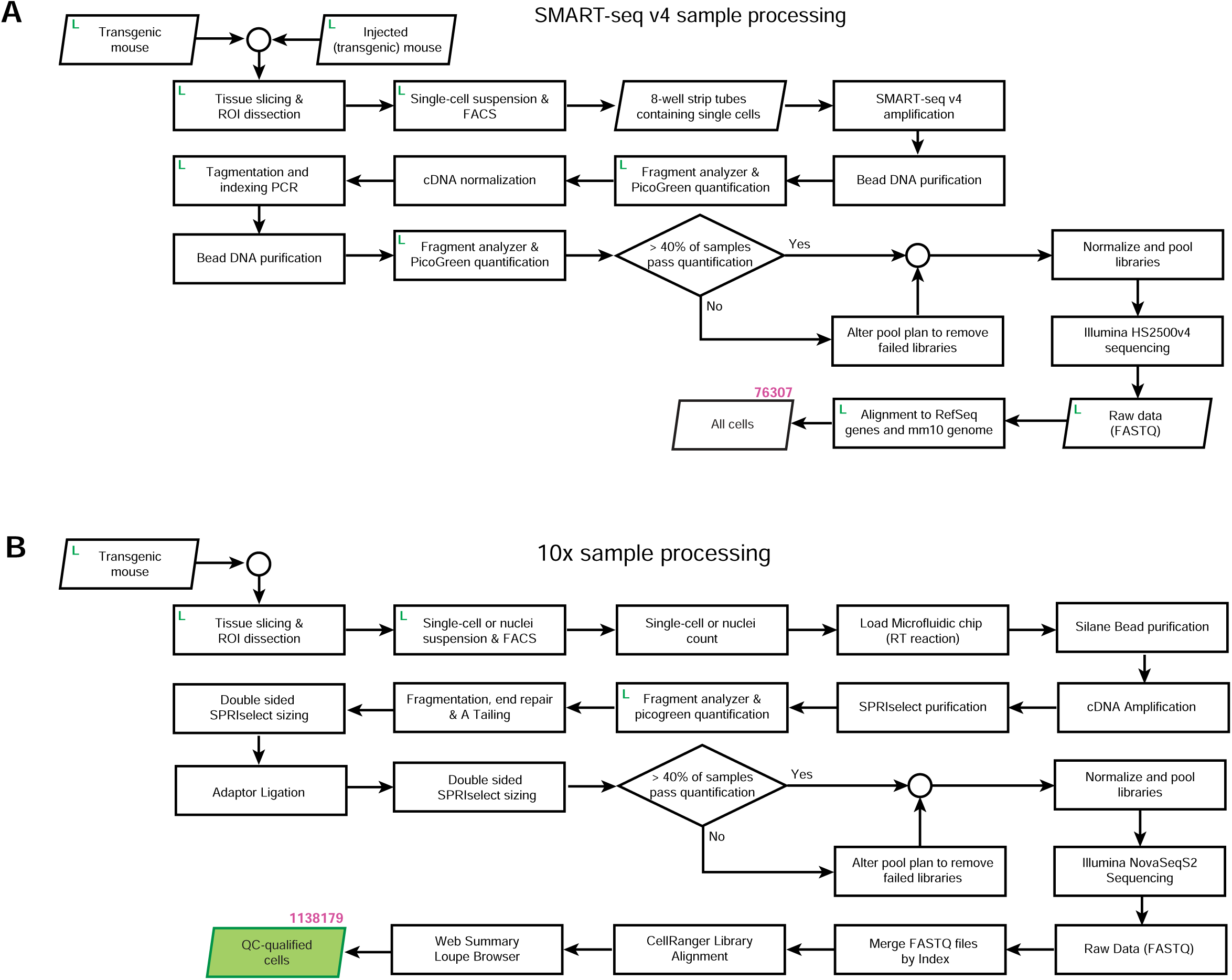
scRNA-seq pipeline and data preprocessing workflow. **(A-B)** Workflow outlining the path from individual experimental animals to quality control (QC)-qualified scRNA-seq data for SSv4 (**A**) or 10xv2 (**B**). At multiple points throughout sample processing, cell and sample metadata were recorded in a laboratory information management system (LIMS, labelled as L), which informs QC processes. Samples must pass QC benchmarks to continue through sample processing.

**Figure S3.**
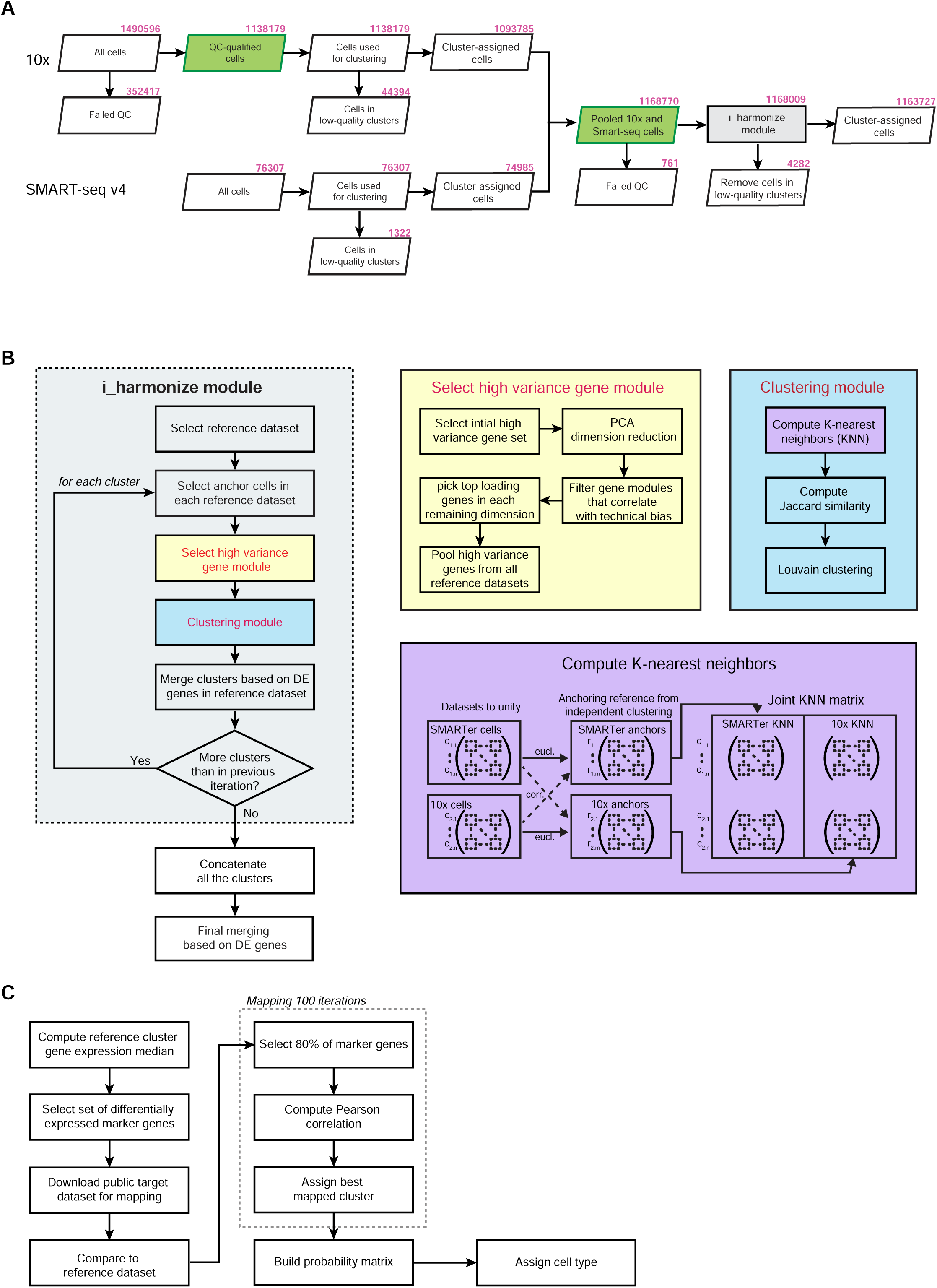
scRNA-seq data analysis workflow. **(A)** The number of cells at each step in the scRNA-seq data analysis pipeline. The identification of doublets and low-quality clusters is described in more detail in **Methods**. The 10xv2 and SSv4 data was first QC-ed and analyzed separately. After initial clustering the datasets were combined and QC-ed again before and after joint clustering. **(B)** Joint clustering procedure using the new i_harmonize function from the scrattch.hicat package (**Methods**). For this study, as the 10xv2 dataset includes more cells while the SSv4 dataset provides more sensitive gene detection, both datasets were used as reference datasets. For each reference dataset anchor cells were selected to achieve uniform coverage of all cell types. Based on these anchor cells high variance genes were selected (select high variance gene module, yellow box), and high variance genes from each reference dataset were pooled. Next, a common adjacency graph using all cells from all datasets was built (purple box) and the standard Jaccard-Louvain clustering algorithm was applied (clustering module, blue box). Resulting clusters were merged to ensure that all pairs of clusters, even at the finest level, were separable by conserved DE genes across platforms. This i_harmonize function applies the integrative clustering across datasets iteratively, while ensuring that all clusters at each iteration are separable by conserved DE genes. **(C)** To assess correspondence of cell types identified in this study to previously published datasets, cells from published datasets were mapped to our clusters using the nearest centroid classifier based on a set of shared markers that were detected in both datasets (expression > 0). To estimate the robustness of mapping, classification was repeated 100 times, each time using 80% of randomly sampled markers, and the probability for each cell to map to every reference cluster was computed.

**Figure S4.**
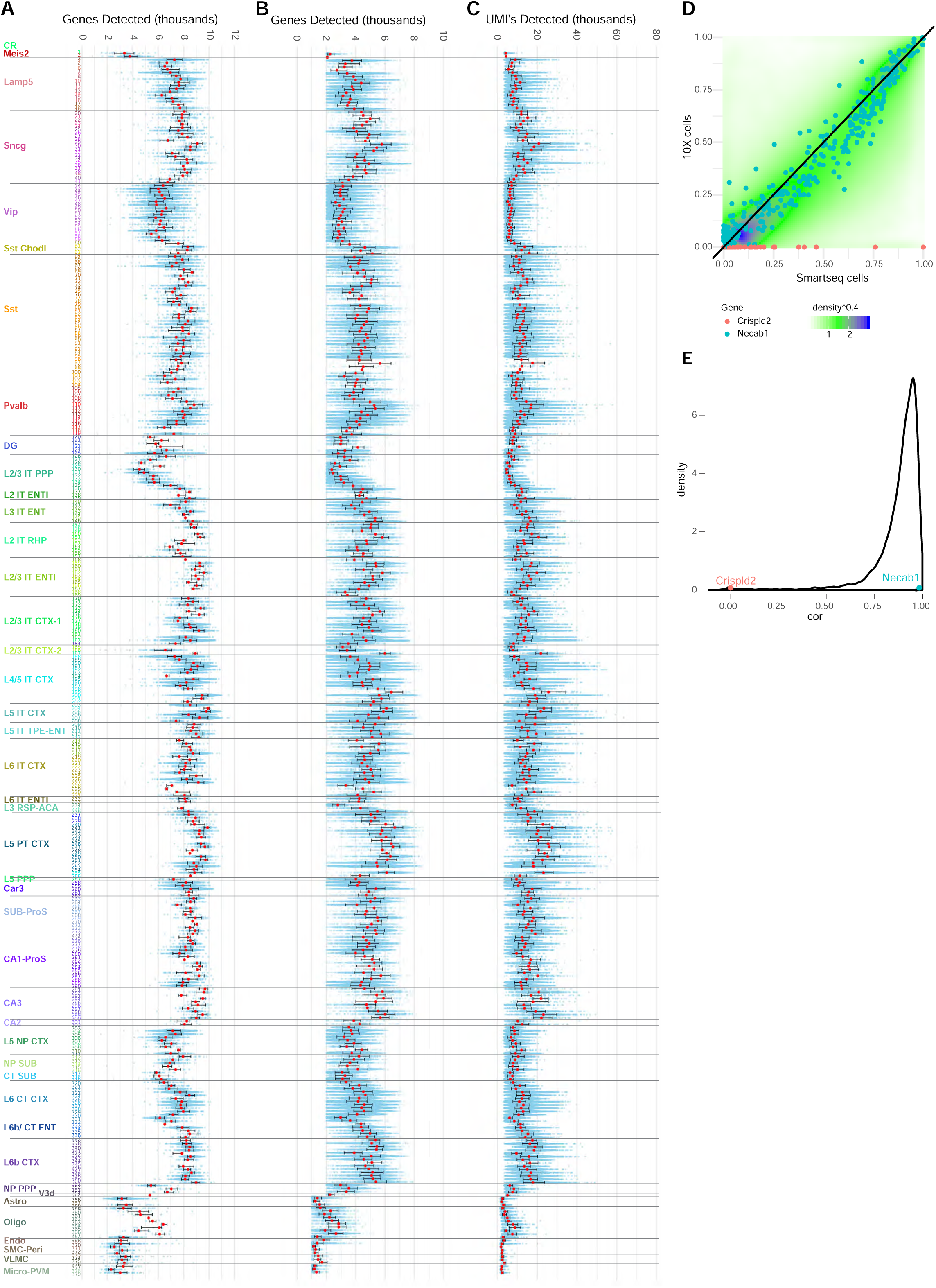
Gene and UMI detection rates in the datasets. **(A-B)** Number of genes detected per SSv4 (**A**) or 10xv2 (**B**) cell for each cluster. **(C)** Number of UMI’s detected per 10xv2 cell for each cluster. The average number of UMIs detected per cell in 10xv2 data was 11,668. The numbers of genes or UMIs detected were largely consistent within each class and subclass of cells; non-neuronal cells and the CR and Meis2 neurons had substantially lower numbers of genes detected than other neurons. **(D)** Comparison of the relative expression level of marker genes across all clusters between the SSv4 and 10xv2 datasets. Since the two datasets differ in experimental platform, gene expression quantification software and gene annotation reference, for each gene, we normalized the average log2(CPM+1) values at the cluster level in the range [0,1] by subtracting the minimum value and then dividing them by the maximum value for that gene. The smooth scatter plot represents normalized gene expression for all marker genes across all clusters in the two datasets. The areas with the highest density of data points are colored blue, and the lowest density white. Two example genes are shown, each dot representing the average expression level for each gene in each cluster. **(E)** Distribution of gene expression conservation between the two platforms. For each of 4,963 marker genes, we computed the correlation of its average expression across all overlapping cell types between 10xv2 and SSv4, and distribution of such correlation values (‘cor’) is shown in the density plot. Two example genes shown in panel **D** were highlighted: *Necab1* represents a gene with high correspondence between the two datasets, while *Crispld2* is detectable in SSv4 cells but not in 10xv2 cells.

**Figure S5.**
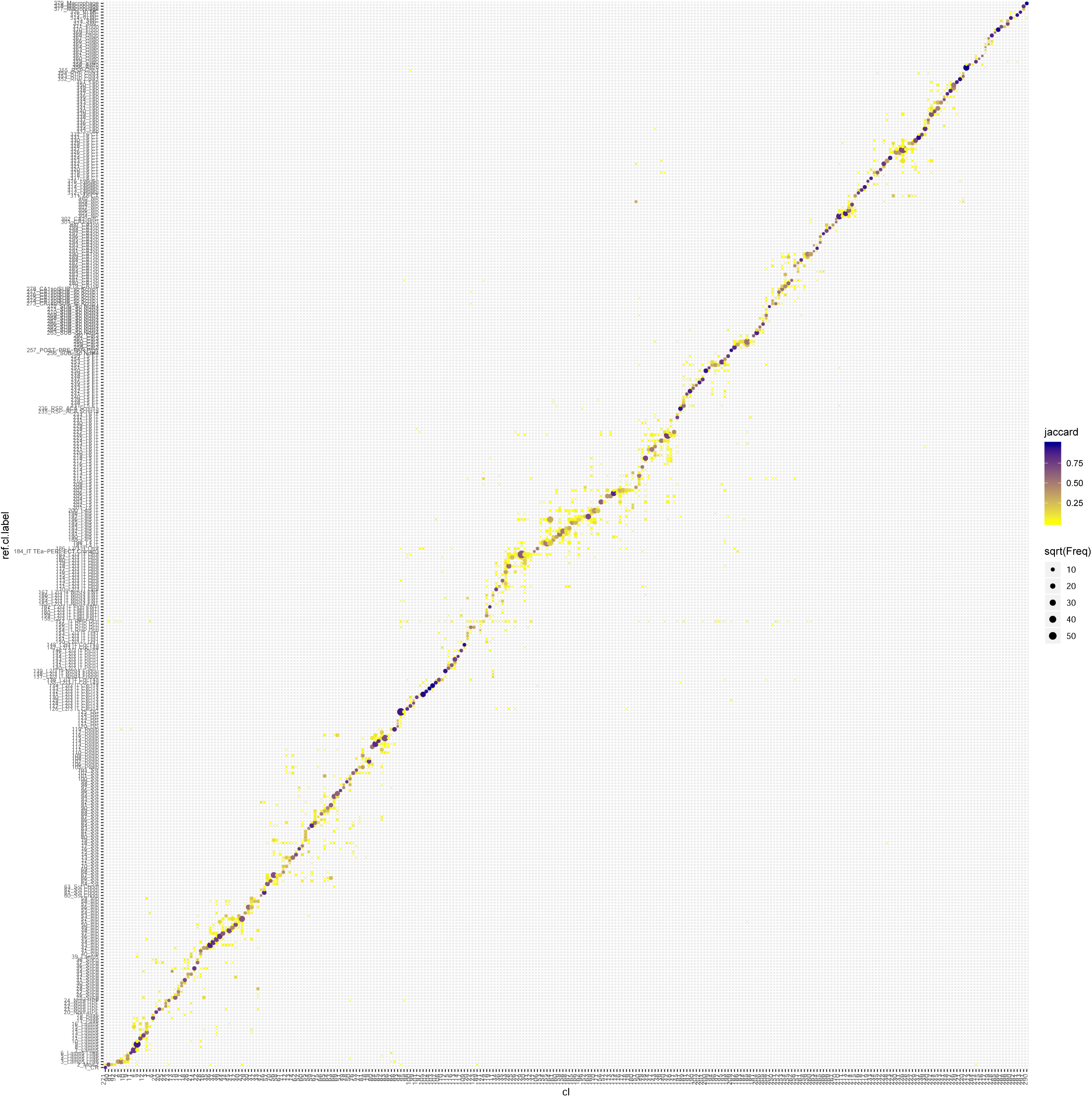
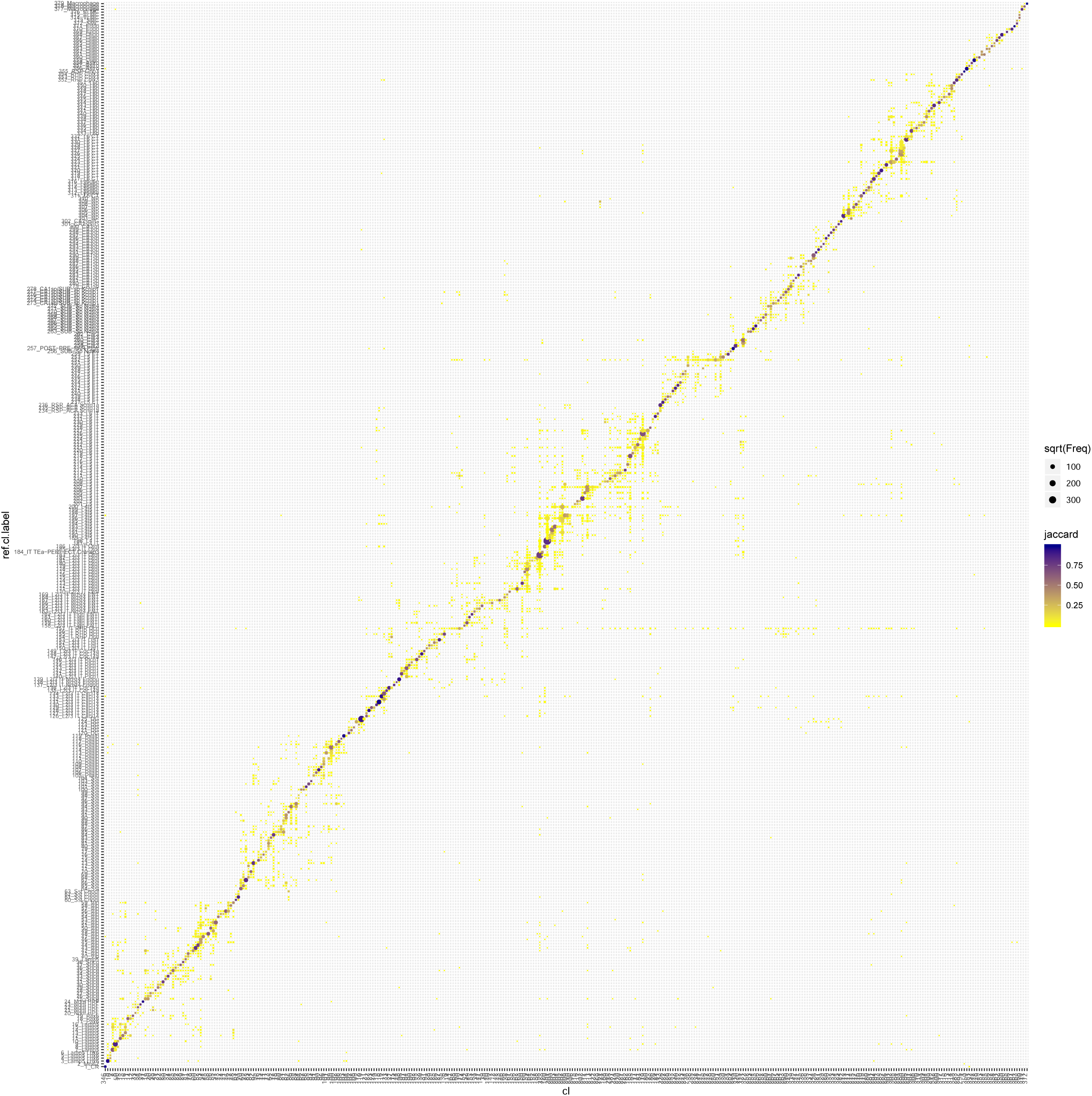
Correspondence between joint taxonomy and platform-specific taxonomy. **(A-B)** Cells from SSv4-only clusters (**A**) or 10xv2-only clusters (**B**) were mapped to the joint taxonomy using the method described in **Figure S3C** and **Methods**. Dot size represents the number of cells mapped to a specific cluster; color represents the similarity (Jaccard distance).

**Figure S6.**
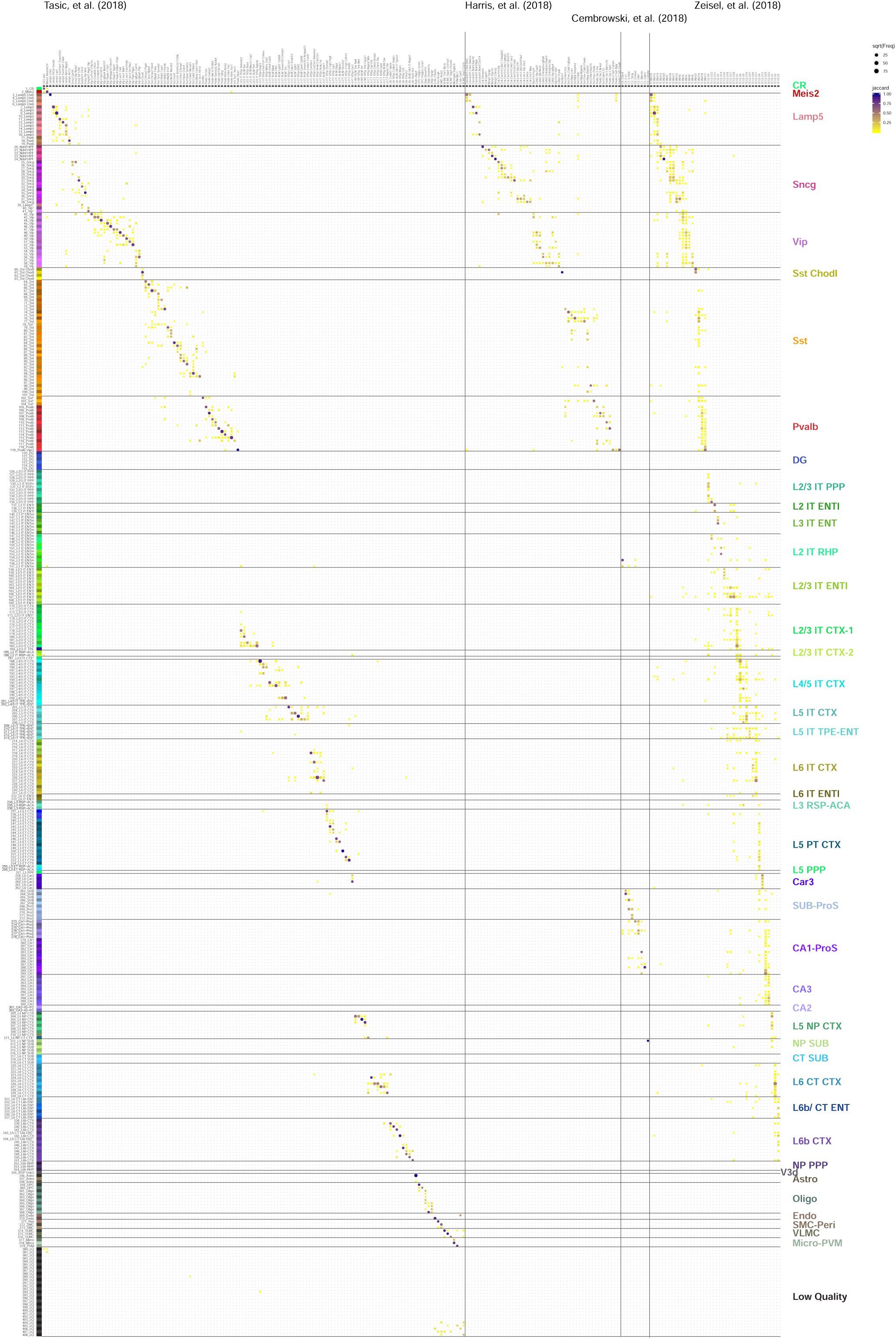
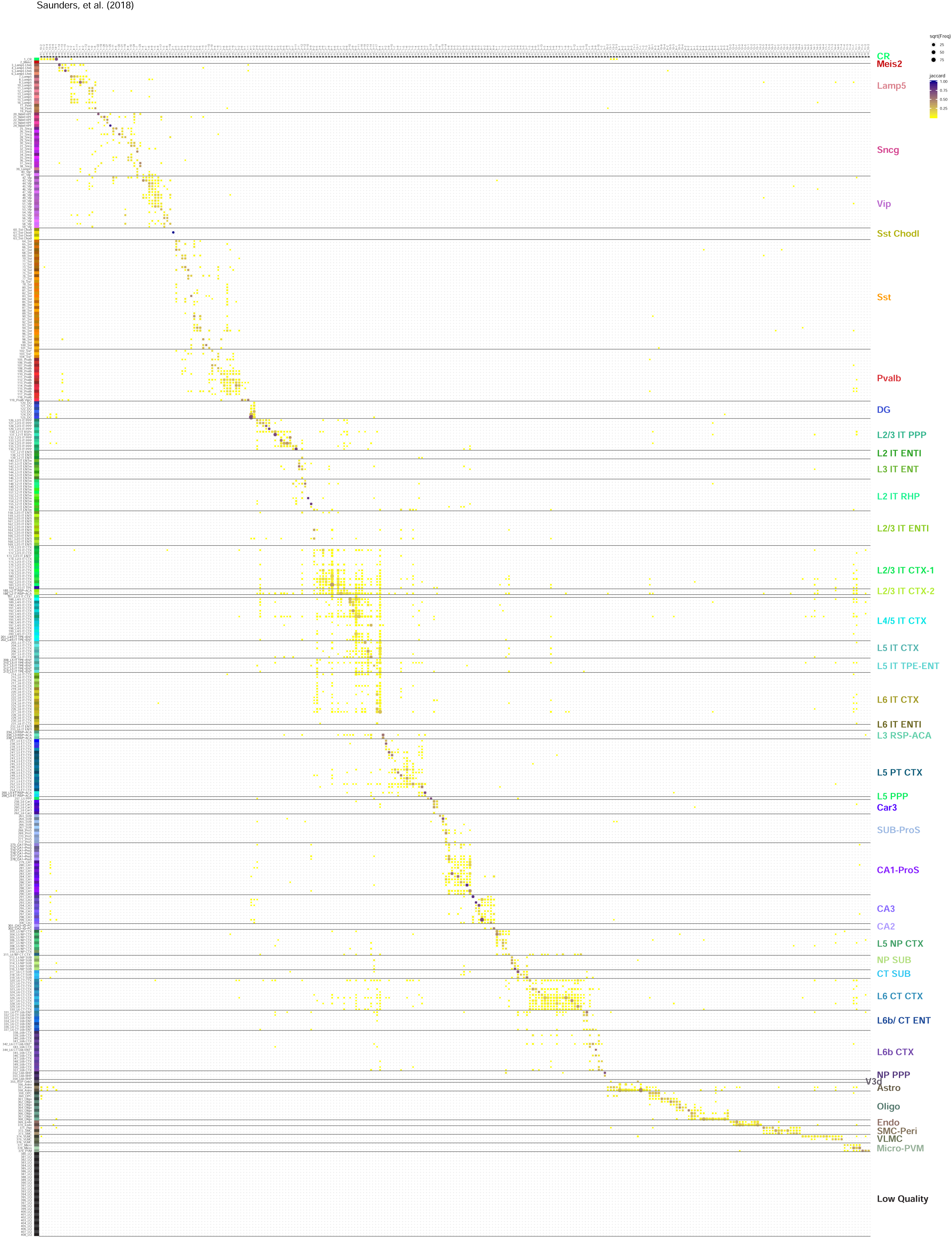
Correspondence between the current transcriptomic taxonomy and previously published ones. **(A-B)** Correspondence was determined by mapping cells from previously published datasets to the current taxonomy as described in **Figure S3C** and **Methods**. Published datasets include: **A**: Tasic et al 2018, Harris et al 2018, Cembrowski et al 2018, and Zeisel et al 2018; **B**: Saunders et al 2018.

**Figure S7.**
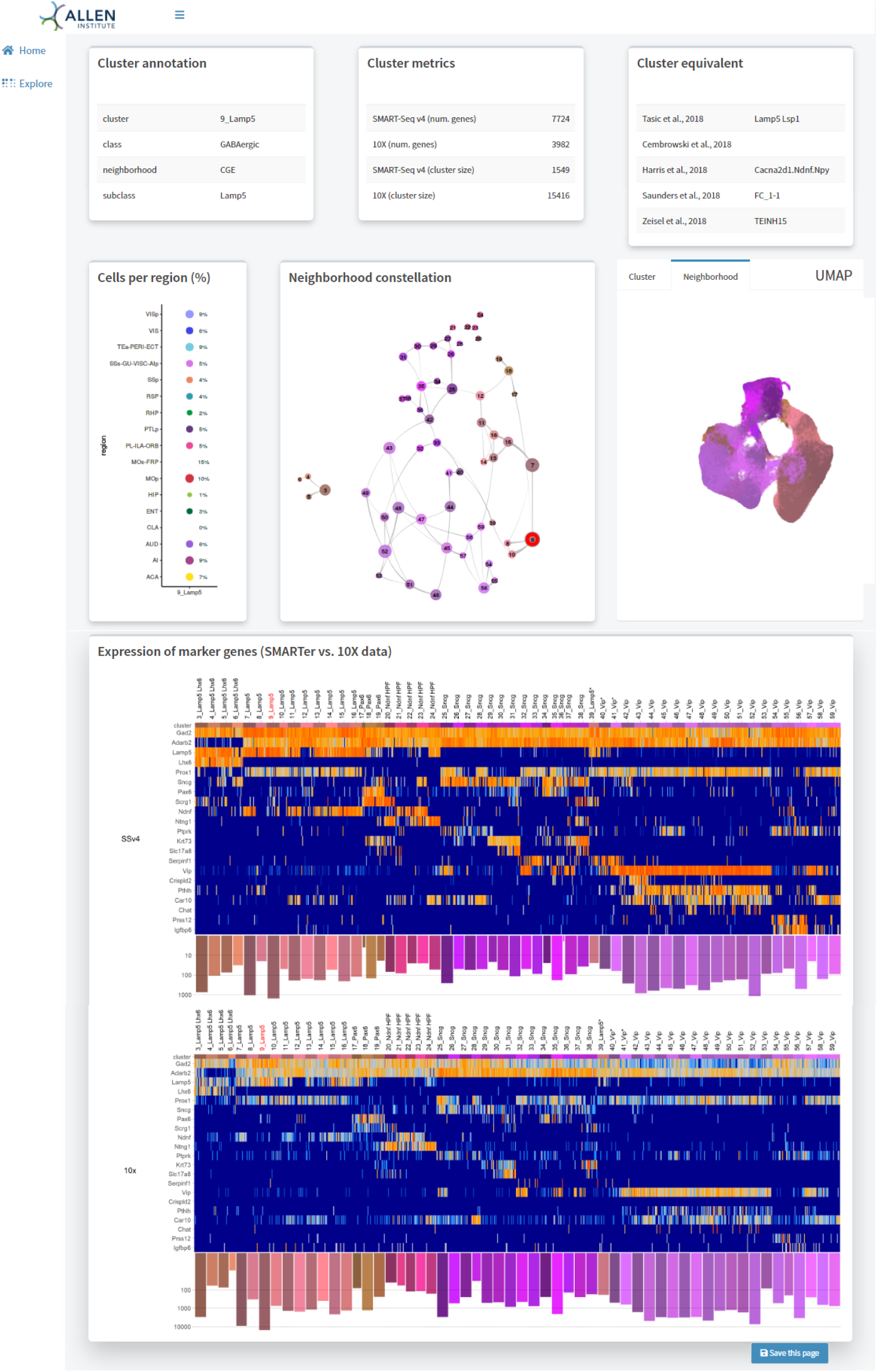
Cell type cards. Example of a cell type card, summarizing information for a specific type within a neighborhood. The cell type card includes information about cluster annotation, cell sampling, marker gene expression, relationship to other clusters in that neighborhood, and correspondence to previously published cell types.

**Figure S8.**
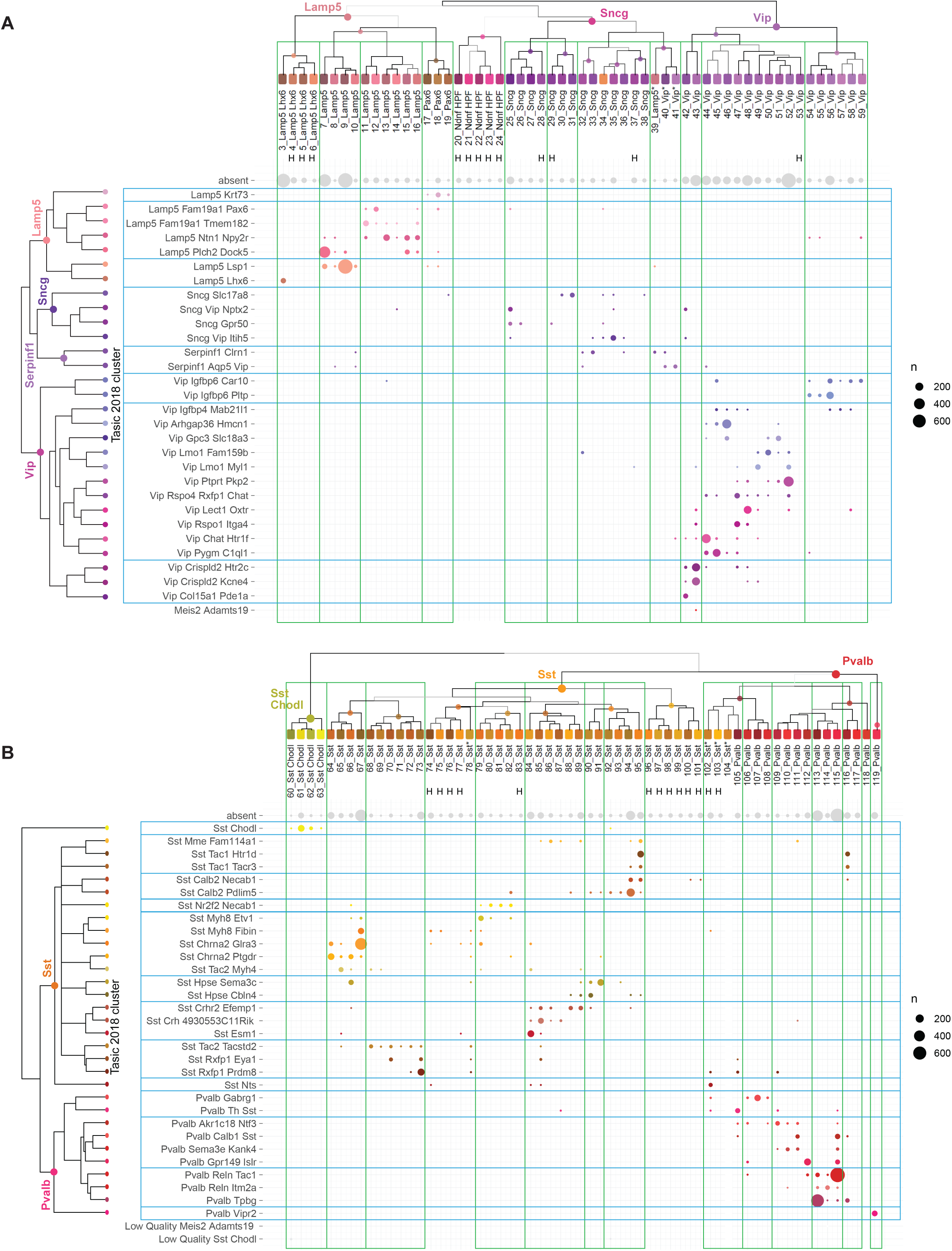
Correspondence between the current CTX-HPF GABAergic taxonomy and previous VISp-ALM taxonomy. **(A-B)** Correspondence between CGE types (**A**) or MGE types (**B**) in the current taxonomy and in the VISp-ALM taxonomy (Tasic et al., 2018). The cells included in the VISp-ALM taxonomy were also included in the current taxonomy and their cluster assignments in both taxonomies were compared in the dot plots. Unlike in **Figure S6A**, no additional mapping computation was performed here. Dot size represents the number of cells included.

**Figure S9.**
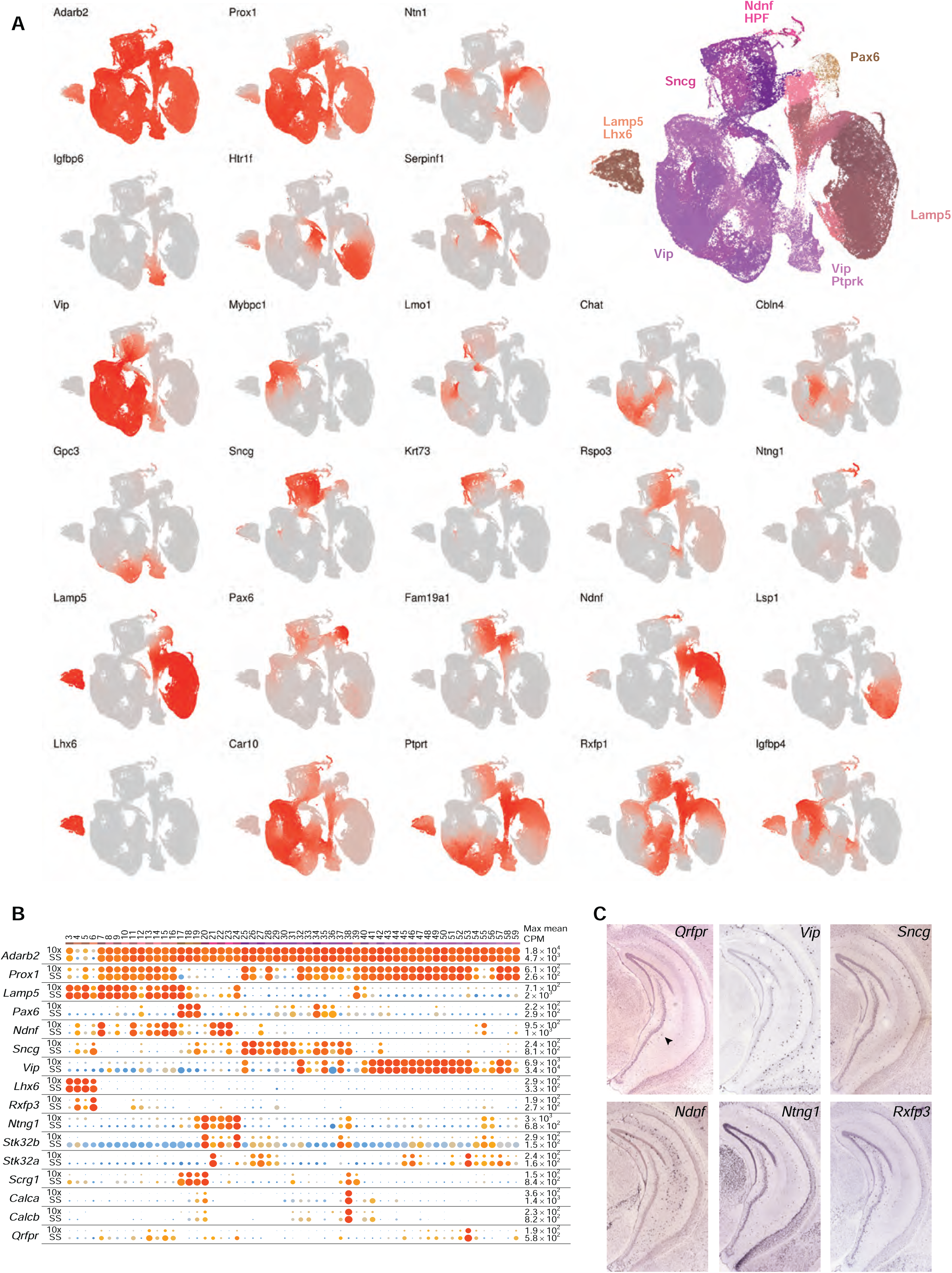
Marker gene expression in supertypes in the CGE GABAergic subclasses. **(A)** UMAP representation of CGE types showing expression of select supertype marker genes in red. The larger inset in the top right is a UMAP representation of CGE types colored by clusters and annotated with supertype labels. **(B)** Dot plot showing marker gene expression in clusters from the 10xv2 (top row) and SSv4 (bottom row) datasets. Dot size and color indicate proportion of expressing cells and average expression level in each cluster, respectively. **(C)** RNA ISH images for select markers expressed in the HPF-enriched CGE cell types.

**Figure S10.**
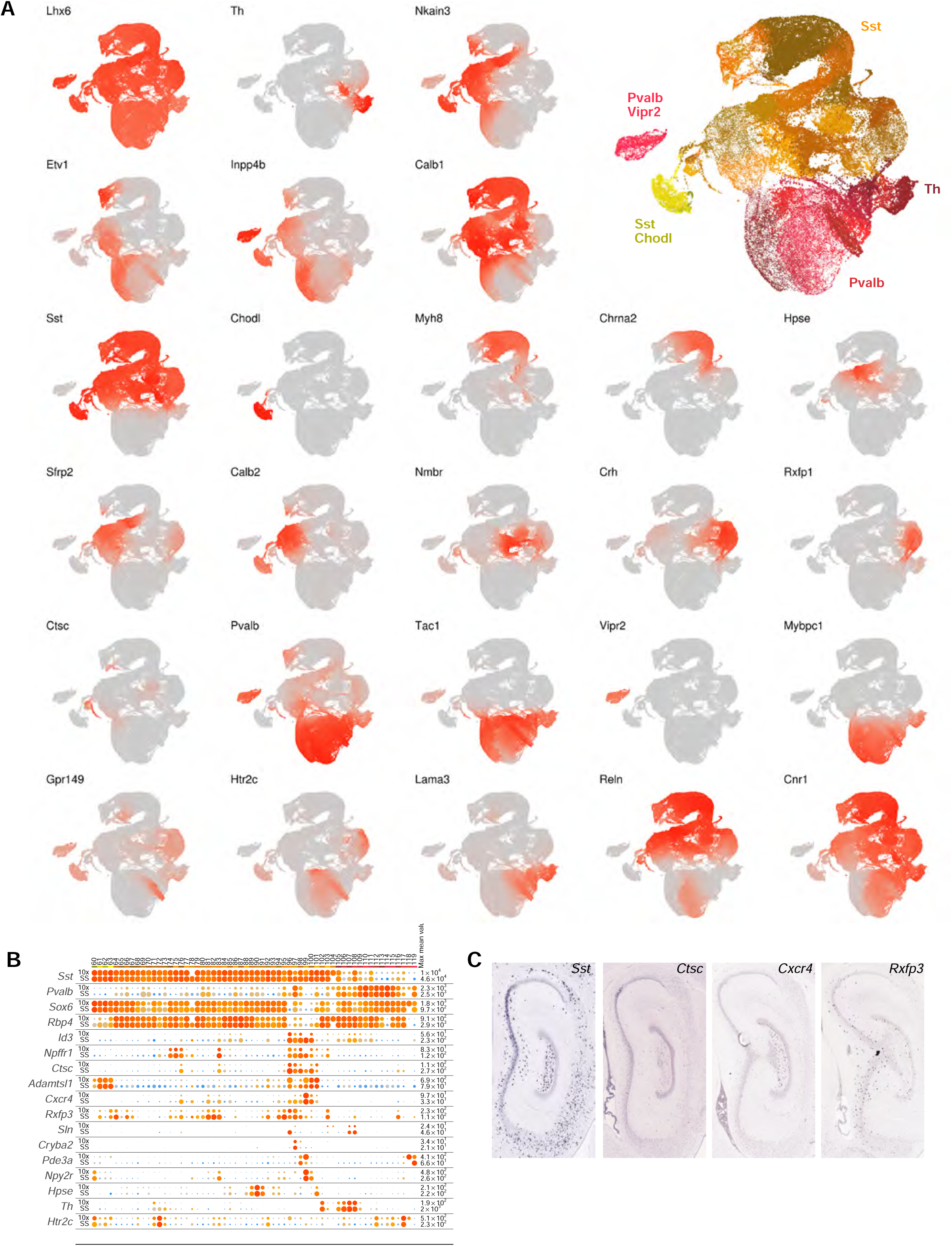
Marker gene expression in supertypes in the MGE GABAergic subclasses. **(A)** UMAP representation of MGE types showing expression of select supertype marker genes in red. The larger inset in the top right is a UMAP representation of MGE types colored by clusters and annotated with supertype labels. **(B)** Dot plot showing marker gene expression in clusters from the 10xv2 (top row) and SSv4 (bottom row) datasets. Dot size and color indicate proportion of expressing cells and average expression level in each cluster, respectively. **(C)** RNA ISH images for select markers expressed in the HPF-enriched MGE cell types.

**Figure S11.**
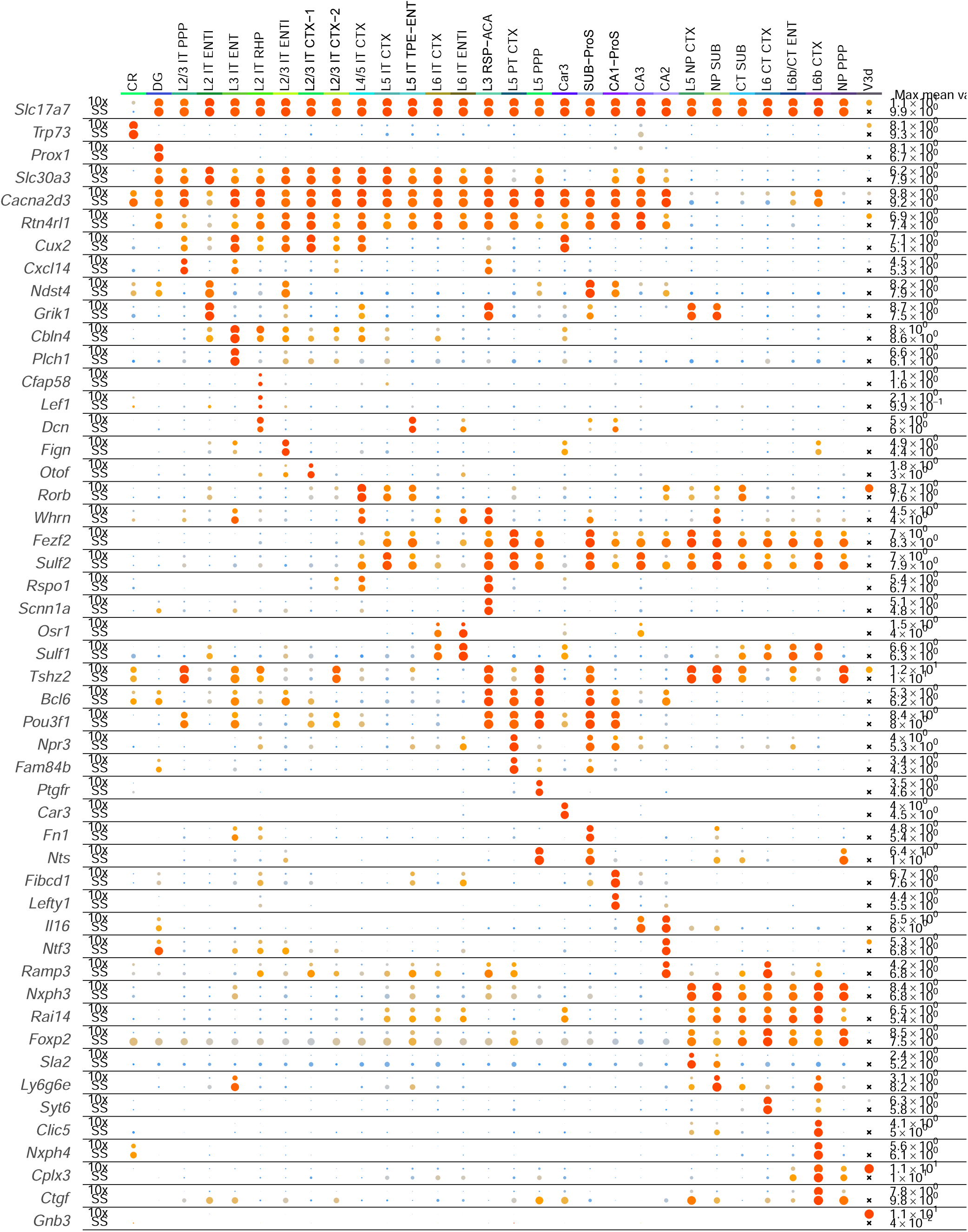
Marker gene expression among glutamatergic subclasses. The dot plot illustrates marker gene expression in glutamatergic excitatory subclasses from the 10xv2 (top row) and SSv4 (bottom row) datasets. Dot size and color indicate proportion of expressing cells and average expression level in each cluster, respectively.

**Figure S12.**
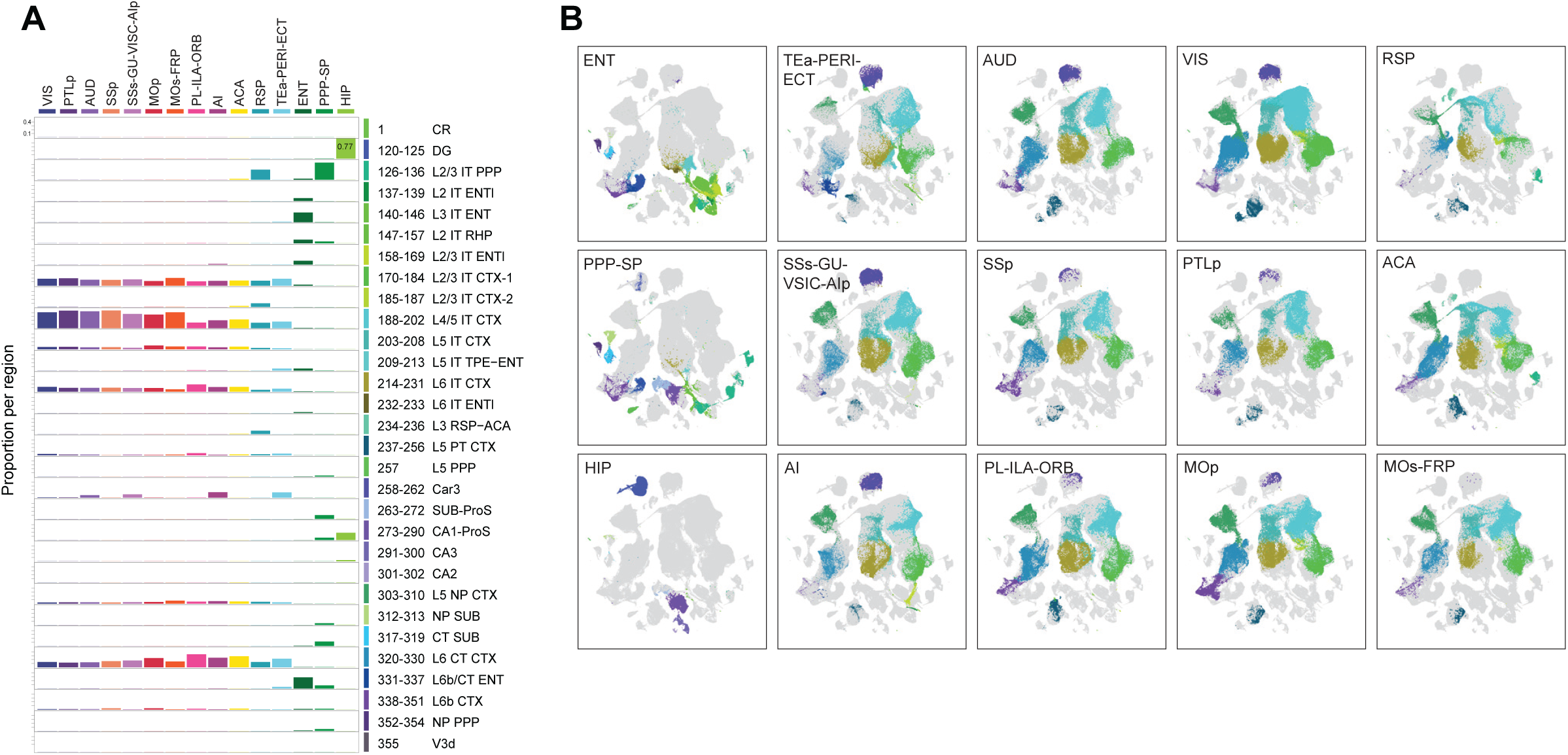
Regional distribution of glutamatergic subclasses. **(A)** The glutamatergic neuronal class was subdivided into 28 subclasses and 2 highly distinct types (CR and V3d). The bar graph shows the proportion of cells within each subclass derived from each region (the values in each region/column add up to 100%). Cells included: 10xv2: *n* = 909,541; SSv4: *n* = 49,198. **(B)** Same UMAP representations as in **Figure 3A**, colored by subclass and separated by region. The highlighted cells are derived from the specified region and cells from other regions are greyed out.

**Figure S13.**
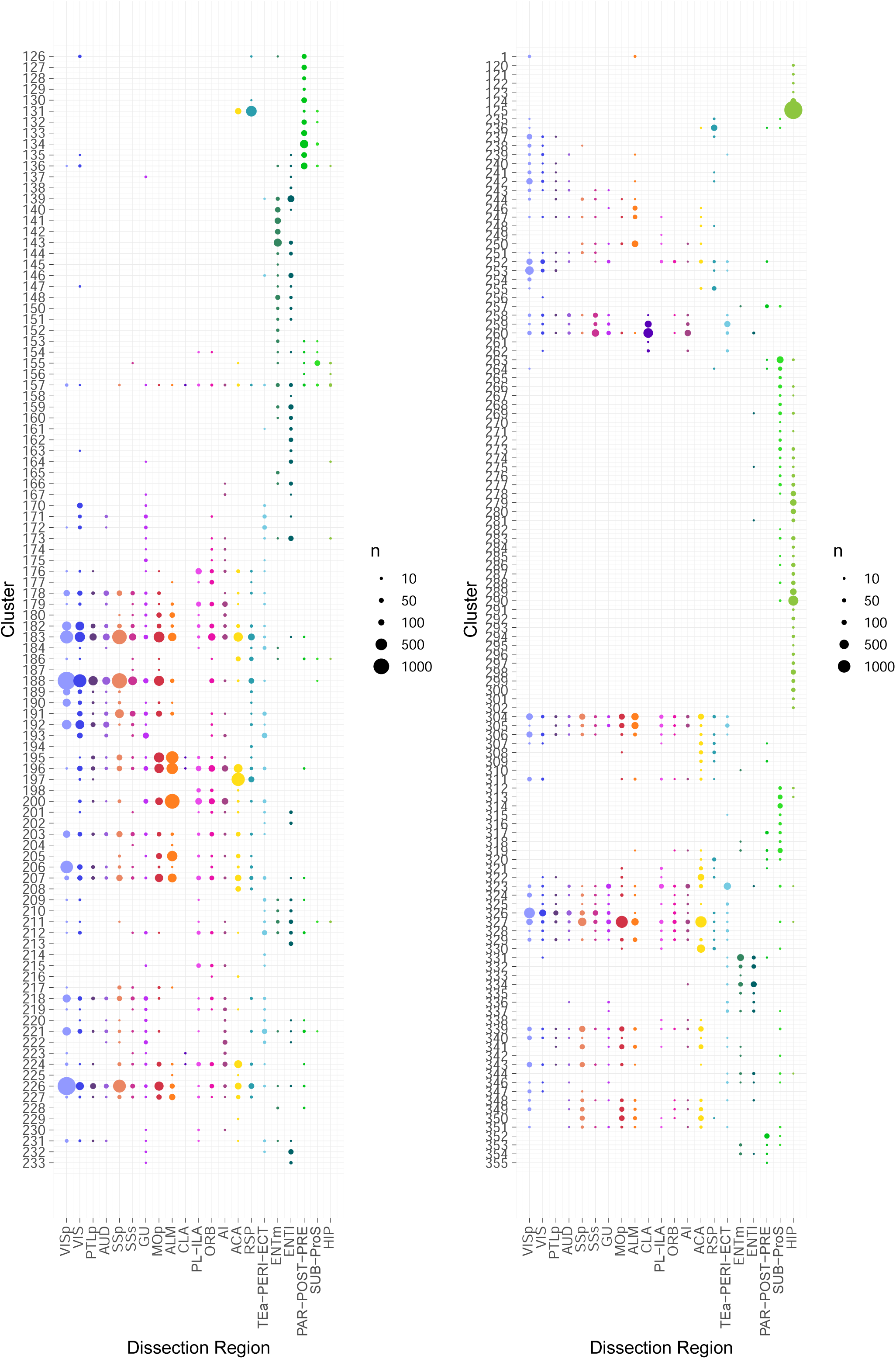
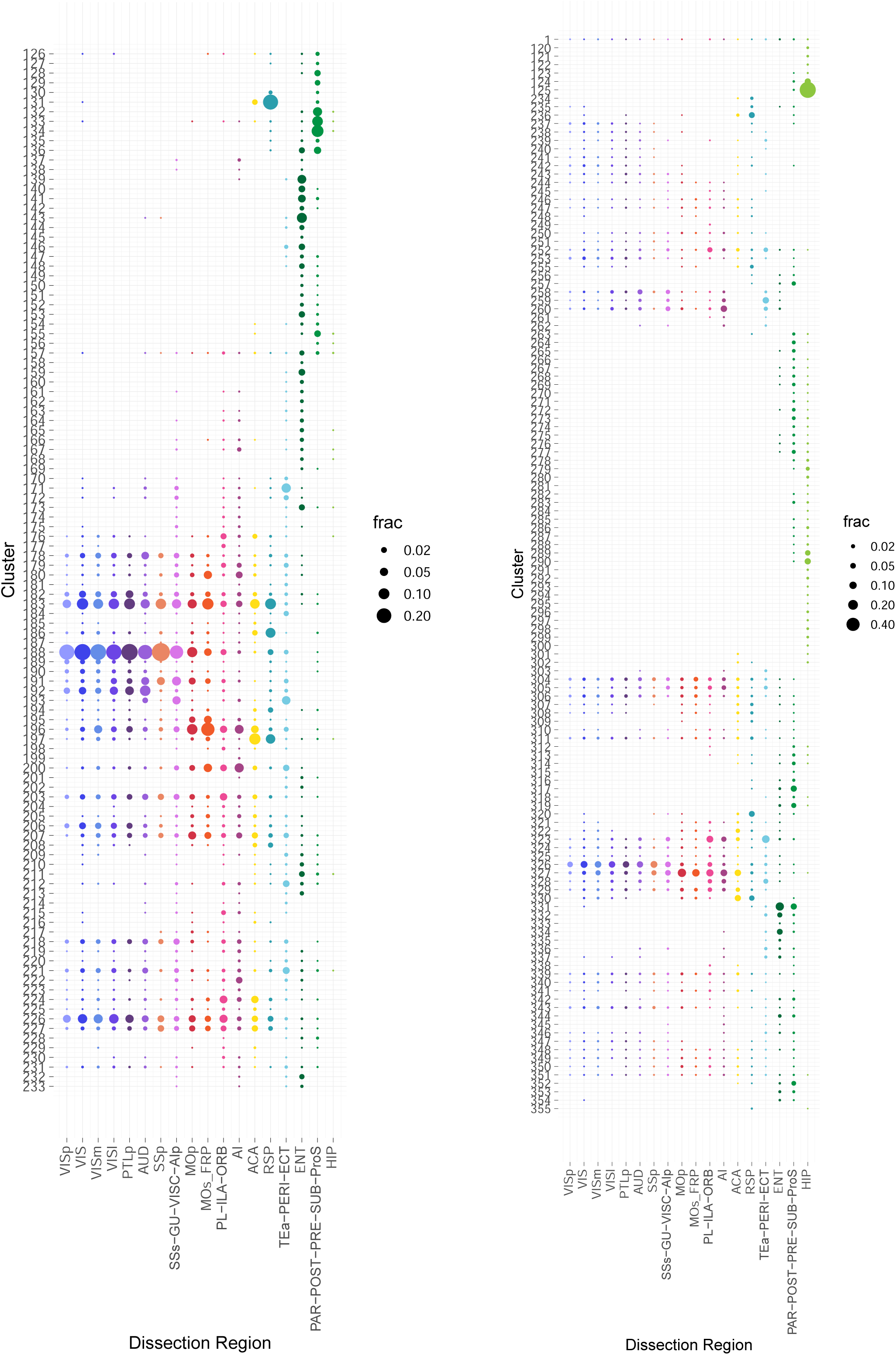
Regional distribution of glutamatergic types. **(A)** Number of SSv4 cells (dot size) for each glutamatergic excitatory cluster derived from each dissection region. **(B)** Fraction of 10xv2 cells (dot size) for each glutamatergic excitatory cluster derived from each dissection region (normalized per region).

**Figure S14.**
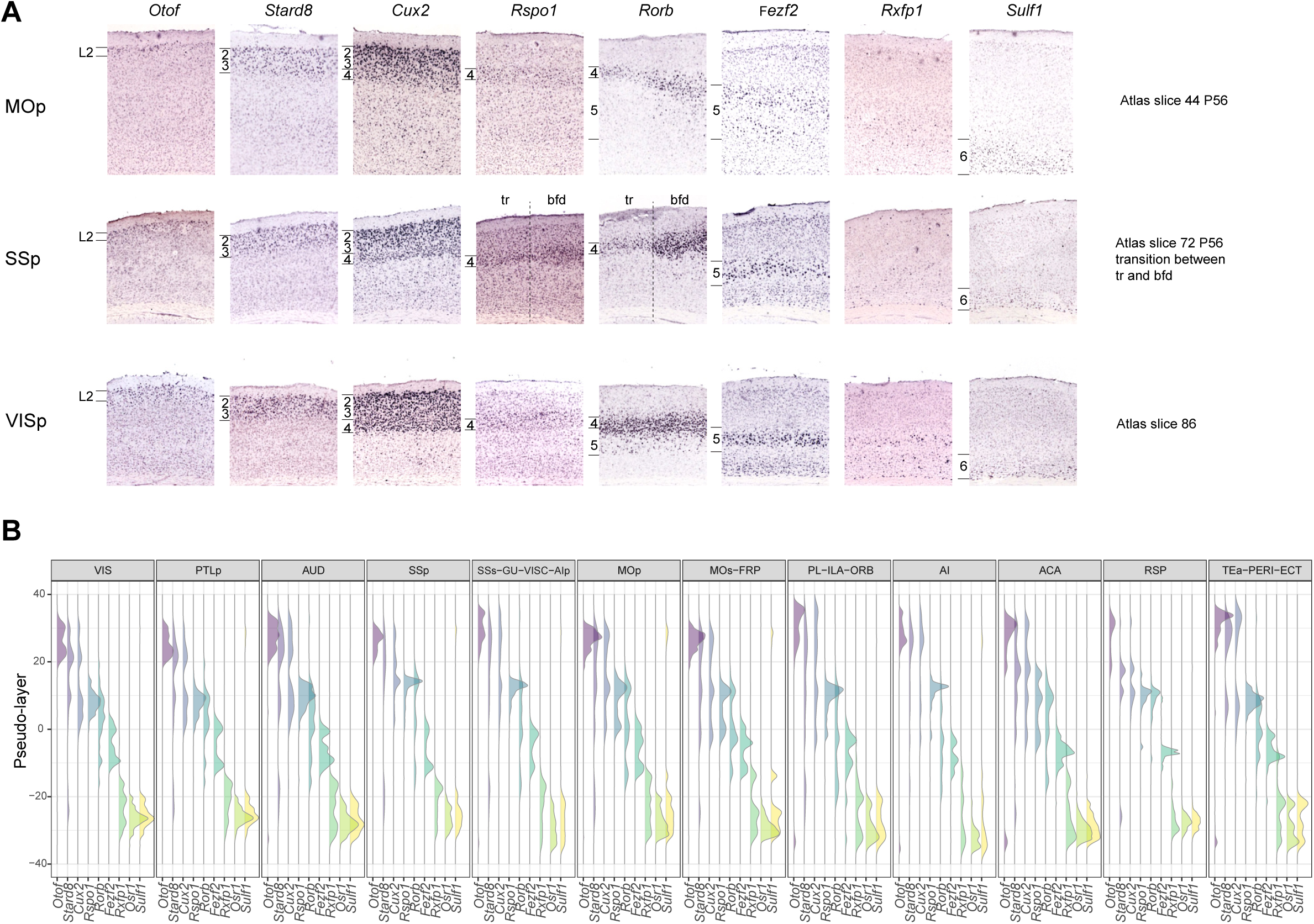
IT pseudo-layer scores and corresponding layer-specific marker genes. **(A)** RNA ISH images for select markers that are expressed by IT cells in specific layers of three isocortical areas: MOp, SSp and VISp. **(B)** Pseudo-layer distribution, calculated as described in **Figure 4B-C**, of markers in panel **A** in 12 different isocortical areas.

**Figure S15.**
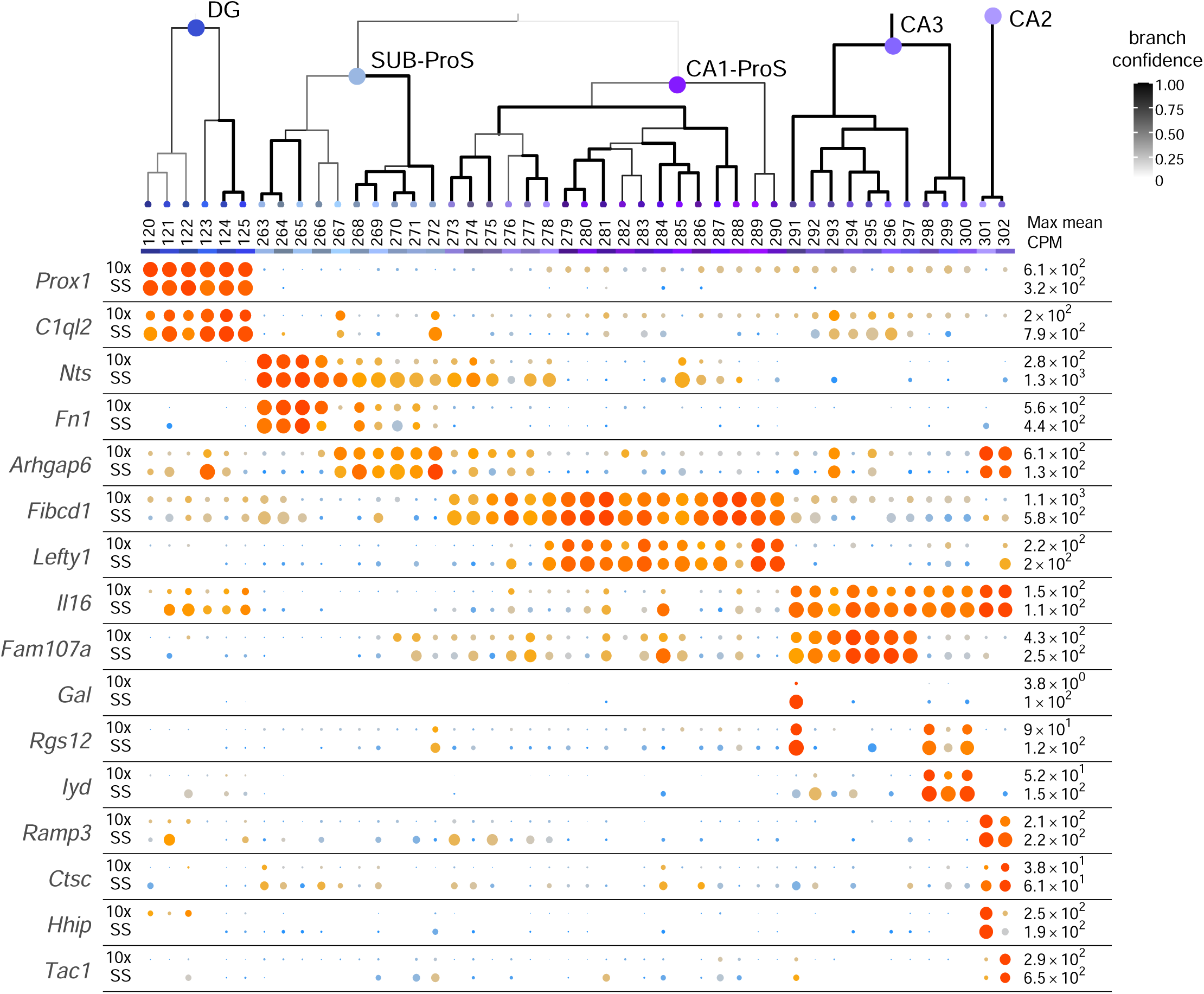
Subclass and supertype marker genes in the hippocampal and subicular glutamatergic types. Dendrogram of 46 clusters in the SUB/HIP neighborhood followed by dot plot illustrating marker gene expression in clusters from the 10xv2 (top row) and SSv4 (bottom row) datasets. Dot size and color indicate proportion of expressing cells and average expression level in each subclass, respectively.

**Figure S16.**
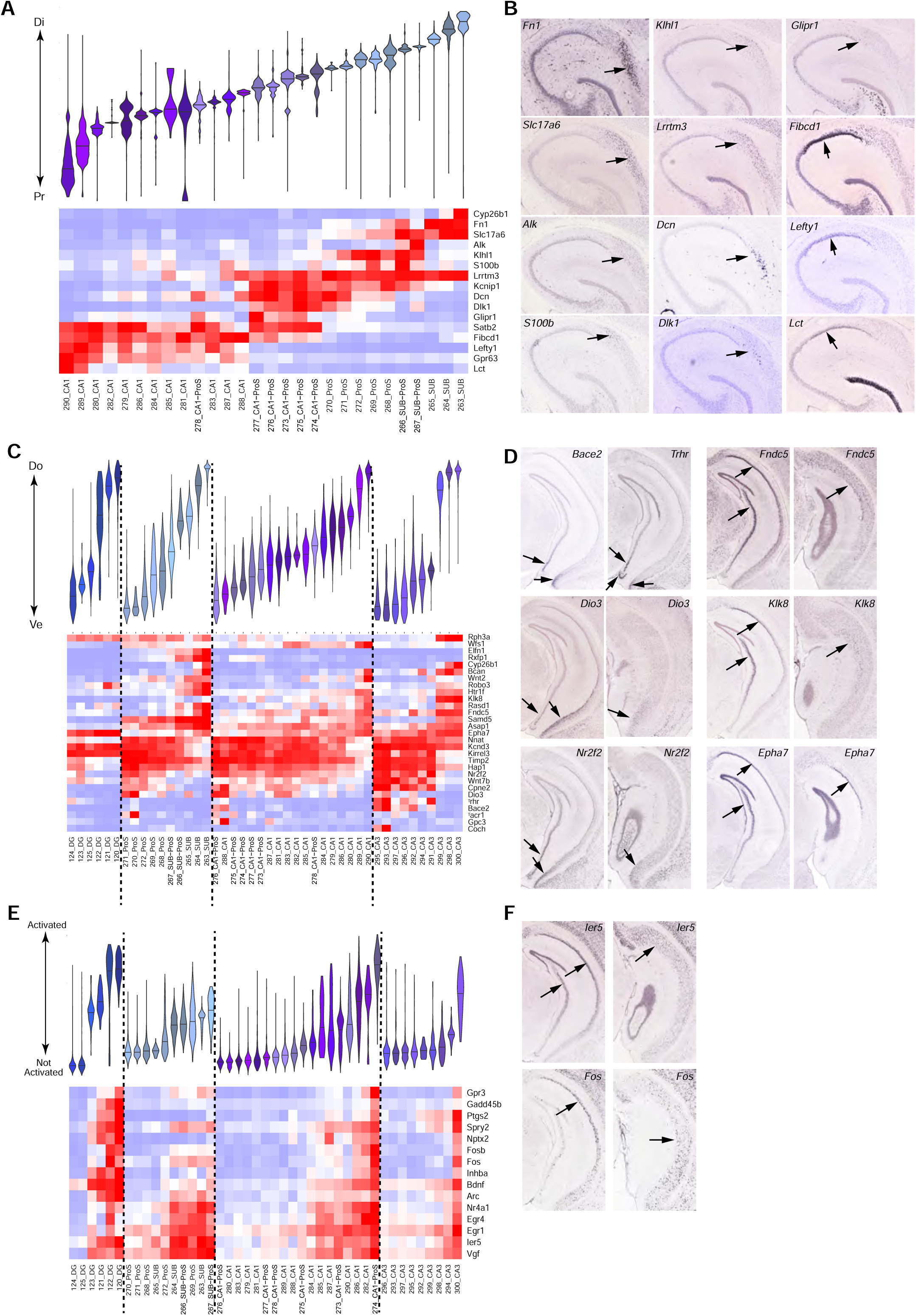
Multi-dimensional gradient distribution of hippocampal and subicular glutamatergic types. **(A)** Distribution of clusters along the proximal-distal axis in SUB/ProS/CA1. Clusters are sorted based on median value. Bottom panel shows the average expression in each cluster of select markers correlated with this gradient. Gene expression is normalized by dividing the maximal value per gene in range [0,1] from blue to red. **(B)** RNA ISH images for select markers showing the proximal-distal transition. **(C)** Distribution of clusters along the ventral-dorsal axis within each subclass (separated by dashed lines). Clusters are sorted within each subclass based on median value. Bottom panel shows the average expression in each cluster of select markers correlated with this gradient, normalized as in A. **(D)** RNA ISH images for select markers showing the ventral-dorsal transition. **(E)** Distribution of clusters along the activity axis within each subclass (separated by dashed lines). Clusters are sorted within each subclass based on median value. Bottom panel shows the average expression in each cluster of select markers correlated with this gradient, normalized as in A. **(F)** RNA ISH images for select markers that are expressed in activated cells.

**Figure S17.**
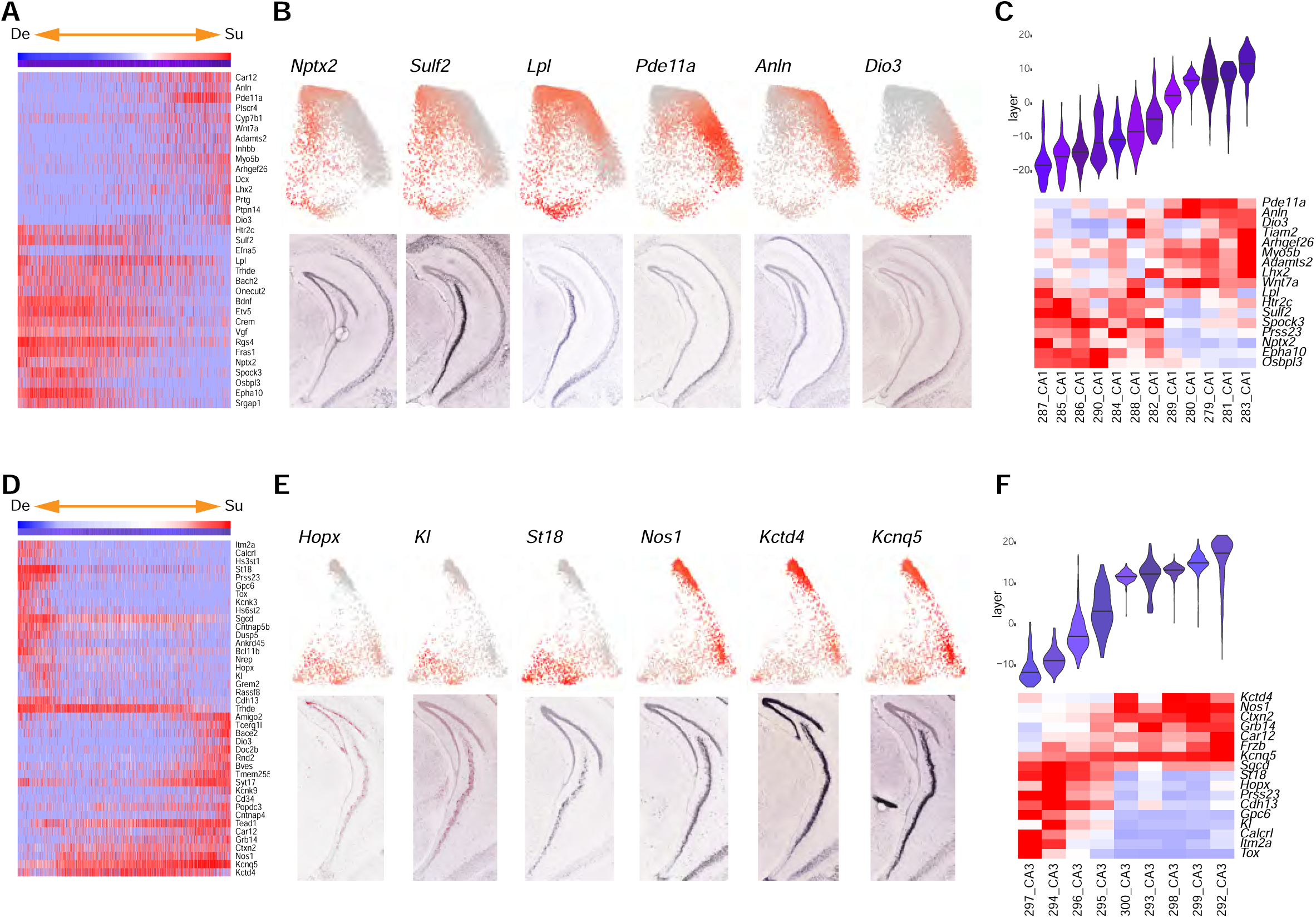
(A) Heatmap of genes that are highly correlated with the second PC of CA1 (2D PC plot shown in **Fig. 8G)** in both directions. Cells are sorted based on this PC, which corresponds to the superficial-deep axis. The top color bar corresponds to the PC value, and the second color bar corresponds to the cluster membership. Gene expression is shown in blue-red scale, with maximal value in red, and no expression in blue. **(B)** Top: imputed expression of select markers in CA1 2D PC plot. Maximal expression in red, no expression in gray. Bottom, ISH images of these makers. **(C)** Top: Distribution of clusters along the CA1 superficial-deep axis. Clusters are sorted based on the median value. Bottom: average expression of select markers in each cluster. **(D)** Similar to A. Heatmap of genes that are highly correlated with the second PC of CA3 (2D PC plot shown in **Fig. 8H**). **(E)** Top: imputed expression of select markers in CA3 2D PC plot. Maximal expression in red, no expression in gray. Bottom, ISH images of these makers. **(F)** Top: Distribution of clusters along the CA3 superficial-deep axis (Note that this axis was recomputed to eliminate the effect of confounding factors. See **Methods** for details). Clusters are sorted based on the median value. Bottom: average expression of select markers in each cluster.

**Table S1. Cell sampling per region** Number of cells sampled for SMART-Seq and 10x platforms, with full names and abbreviations for each region.

**Table S2. Specimen** All specimens used in this study are listed, with associated donor information (sex, age). When applicable, injection target and injection material are specified for specimens with retrograde or retro-orbital labeling. One donor may be in multiple rows (duplicate values highlighted in “donor_label” column) if multiple regions were dissected (“region_label”) or multiple FACS gating (“facs_population_plan”) were used.

**Table S3. Cluster annotation** Detailed information for each cluster, including membership in broader categories, marker genes, putative region, and sampling.

